# Quantitative and systematic NMR measurements of sequence-dependent A-T Hoogsteen dynamics uncovers unique conformational specificity in the DNA double helix

**DOI:** 10.1101/2024.05.15.594415

**Authors:** Akanksha Manghrani, Atul Kaushik Rangadurai, Or Szekely, Bei Liu, Serafima Guseva, Hashim M. Al-Hashimi

## Abstract

The propensities to form lowly-populated short-lived conformations of DNA could vary with sequence, providing an important source of sequence-specificity in biochemical reactions. However, comprehensively measuring how these dynamics vary with sequence is challenging. Using ^1^H CEST and ^13^C *R*_1ρ_ NMR, we measured Watson-Crick to Hoogsteen dynamics for an A-T base pair in thirteen trinucleotide sequence contexts. The Hoogsteen population and exchange rate varied 4-fold and 16-fold, respectively, and were dependent on both the 3’- and 5’-neighbors but only weakly dependent on monovalent ion concentration (25 versus 100 mM NaCl) and pH (6.8 versus 8.0). Flexible TA and CA dinucleotide steps exhibited the highest Hoogsteen populations, and their kinetics rates strongly depended on the 3’-neighbor. In contrast, the stiffer AA and GA steps had the lowest Hoogsteen population, and their kinetics were weakly dependent on the 3’-neighbor. The Hoogsteen lifetime was especially short when G-C neighbors flanked the A-T base pair. The Hoogsteen dynamics had a distinct sequence-dependence compared to duplex stability and minor groove width. Thus, our results uncover a unique source of sequence-specificity hidden within the DNA double helix in the form of A-T Hoogsteen dynamics and establish the utility of ^1^H CEST to quantitively measure sequence-dependent DNA dynamics.

## Introduction

The sequence of DNA plays crucial roles in many biochemical processes beyond coding for protein and RNA molecules. The sequence determines the position of nucleosomes and the accessibility of various genomic regions to the cellular machinery (1,2). It directs the binding of transcription factors to specific DNA motifs to regulate gene expression (3–5). Beyond these regulatory processes, many mutational processes linked to cancer strongly depend on the DNA sequence (6–8). Some DNA sequences are more susceptible to damage from environmental and endogenous factors (9–17), while others are more prone to replicative errors (18–24). The efficiency of damage repair can also vary depending on the sequence context of the damaged nucleotide (25–28).

There has been a long-standing interest in understanding the molecular origins of DNA sequence-specificity. Some processes directly act on the one-dimensional DNA sequence (29–31). Proteins, for example, can ‘read’ the DNA sequence through base-specific contacts. In addition, adjacent thymidine residues can form thymine dimers following UV exposure, resulting in sequence-specific damage (11,16,17). Specificity can also originate from the three-dimensional (3D) structure of the DNA, which in turn depends on the sequence (32,33). The sequence-specific 3D DNA structure can be specifically recognized by DNA-binding proteins (34,35) and can also determine the accessibility of the DNA and its susceptibility to damage (36).

Another mechanism for DNA sequence-specificity has recently come to light. Biochemical processes frequently act on alternative conformations of DNA base pairs (bps), which form transiently and in low abundance in the naked duplex (37–40). For example, damage repair often requires extra-helical flipping of nucleotides (40–43). Many proteins employ non-canonical Hoogsteen bps to bind DNA (44–46) (Figure 1A). Lowly-populated and short-lived Watson-Crick-like G-T mismatches can evade fidelity checkpoints and contribute to errors during replication, transcription, and translation (24,47–52) and possibly off-target CRISPR-Cas genome-editing (52,53). Because these rare conformational states entail changes in the stacking interactions with neighboring bps, their energetics and kinetics could vary substantially with sequence context (24,54), and they could therefore be another source of sequence-specificity in biochemical reactions.

**Figure 1.**
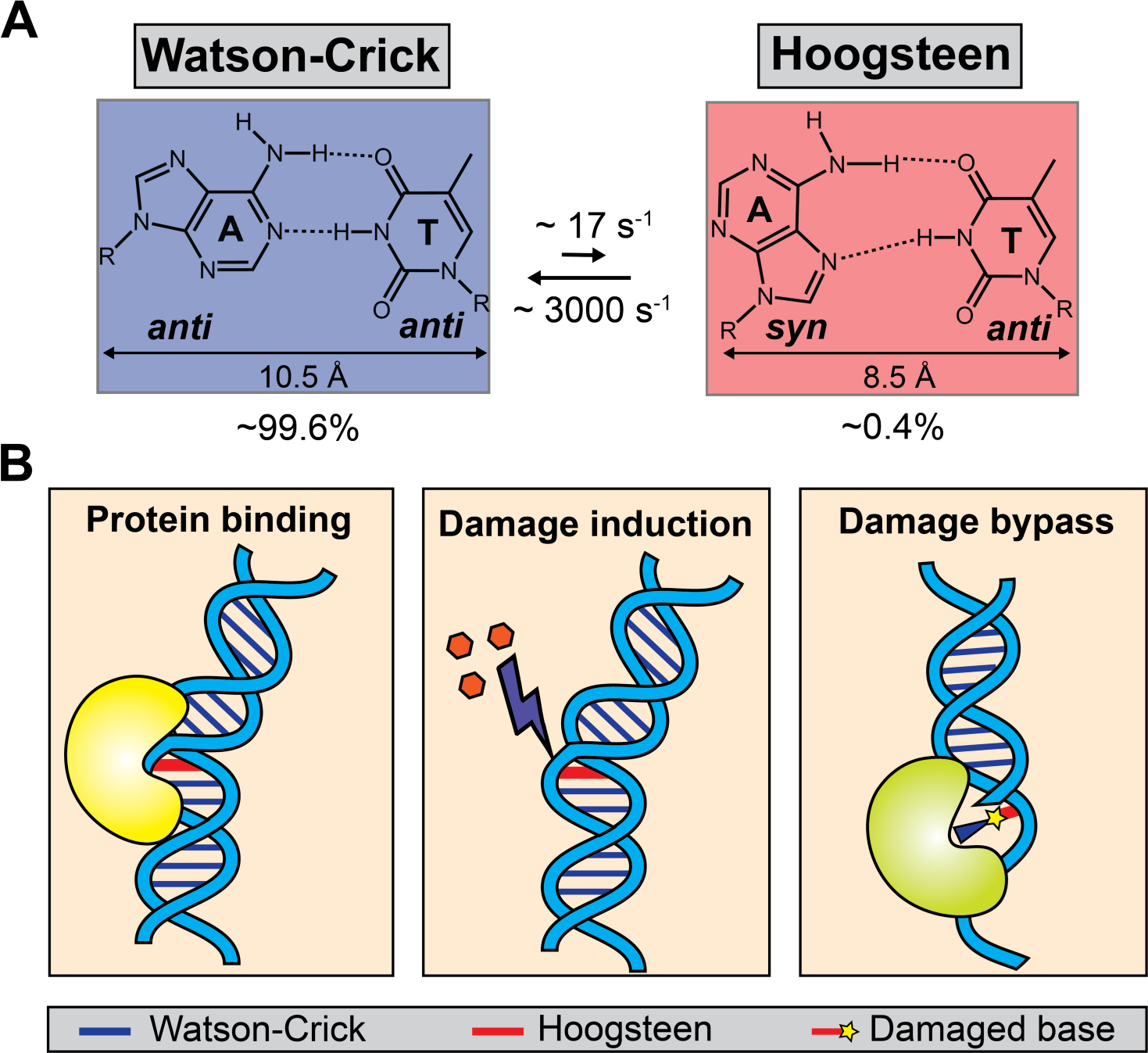
Watson-Crick to Hoogsteen dynamics play diverse roles in biochemical processes. (A) Conformational exchange between canonical A-T Watson-Crick and non-canonical Hoogsteen bps. The Hoogsteen bp forms through 180° rotation of the adenine base followed by constriction of the two bases by ∼ 2-2.5 Å (46). (B) Representative examples of biochemical processes involving the A-T Hoogsteen bp. A-T Hoogsteen bps are proposed to play roles in sequence-specific DNA-protein recognition, DNA damage induction, and damage bypass during replication (63).

NMR methods that measure chemical exchange, including spin relaxation in the rotating frame (*R*_1ρ_) and chemical exchange saturation transfer (CEST) (55–57), provide a rare opportunity to measure the kinetic rates and thermodynamic propensities to form these lowly-populated short-lived conformational states of DNA. However, these experiments are time-consuming and require ^13^C/^15^N isotopic enrichment, which can be cost prohibitive even for a handful of sequences. As a result, few studies have assessed how the dynamics of DNA bps vary with sequence quantitatively and systematically over the expanse of sequence space.

Building on the ^1^H SELective Optimized Proton Experiments (SELOPE) *R*_1ρ_ experiment developed to characterize conformational dynamics in unlabeled RNA (58–60), we recently introduced the high-power ^1^H CEST experiment to measure conformational exchange in unlabeled DNA duplexes by targeting the imino protons of guanine and thymine residues (61). Compared to conventional ^13^C and ^15^N experiments, ^1^H CEST is not only cost-effective, obviating the need for ^13^C/^15^N enrichment of the DNA, but it can also be carried out in higher throughput using sensitive 1D ^1^H experiments. Here, we leveraged ^1^H CEST to measure the sequence-dependence of Watson-Crick to Hoogsteen dynamics for the A-T bp in duplex DNA for all combinations of its neighboring base pairs.

In naked canonical duplex B-DNA, A-T and G-C Watson-Crick bps exist in a dynamic equilibrium with their Hoogsteen counterparts (Figure 1A) (62). The Hoogsteen bps are lowly-populated (population < 1 %) and short-lived (lifetime ∼ 1 ms) and appear to form robustly across a variety of sequence and positional contexts (54,62) (Figure 1A). However, Hoogsteen bps can become the dominant state when duplex DNA is bound by certain proteins (45,63–66) and drugs (67,68) (Figure 1B). Increasingly observed in crystal structures of protein-DNA complexes (64,66,69–71), Hoogsteen bps are proposed to play a role in sequence-specific DNA recognition by transcription factors (44,70,72), in the damage bypass during replication (73,74), as markers of DNA stress (64), and can also increase the susceptibility of DNA to damage (75,76). Initial efforts to assess how the propensities to form Hoogsteen vary with sequence used ^13^C and ^15^N *R*_1ρ_ experiments and isotopically enriched samples to examine a handful of largely AT-rich sequence motifs in varying duplexes and buffer conditions (54,62) or melting experiments that do not provide kinetic information (77).

We recently demonstrated the utility of ^1^H CEST for measuring Hoogsteen dynamics in DNA duplexes without the need for isotopic enrichment (61,78). Across a variety of duplexes and conditions, the Hoogsteen exchange kinetics measured using ^1^H CEST were shown to be in quantitative agreement with counterparts measured using well-established ^13^C and ^15^N *R*_1ρ_ experiments (61,78). Here, using ^1^H CEST and targeted ^13^C *R*_1ρ_ experiments, we quantitatively and systemically measured the Watson-Crick to Hoogsteen dynamics for an A-T bp positioned at the center of a 12-mer duplex in thirteen out of sixteen possible trinucleotide sequence contexts under two salt concentrations. Our results uncover a unique reservoir of conformational specificity hidden within the DNA double helix in the form of sequence-dependent A-T Hoogsteen dynamics and establish the utility of ^1^H CEST to quantitively measure sequence-dependent DNA dynamics.

## Results

### Sequence-dependent ^1^H chemical shifts

To measure the sequence-dependence of A-T Hoogsteen dynamics systematically and quantitatively, we designed sixteen duplexes in which the central A-T bp was embedded in all sixteen trinucleotide sequence contexts (Figure 2A). For all sixteen unlabeled duplexes, the imino resonances were assigned using 2D ^1^H-^1^H NOESY and natural abundance 2D ^1^H-^15^N HSQC experiments (Figure S1A). As expected, all the residues formed Watson-Crick bps as the dominant conformation. Increasing the salt concentration from 25 to 100 mM NaCl resulted in minor chemical shift perturbations (Figure S1B).

**Figure 2:**
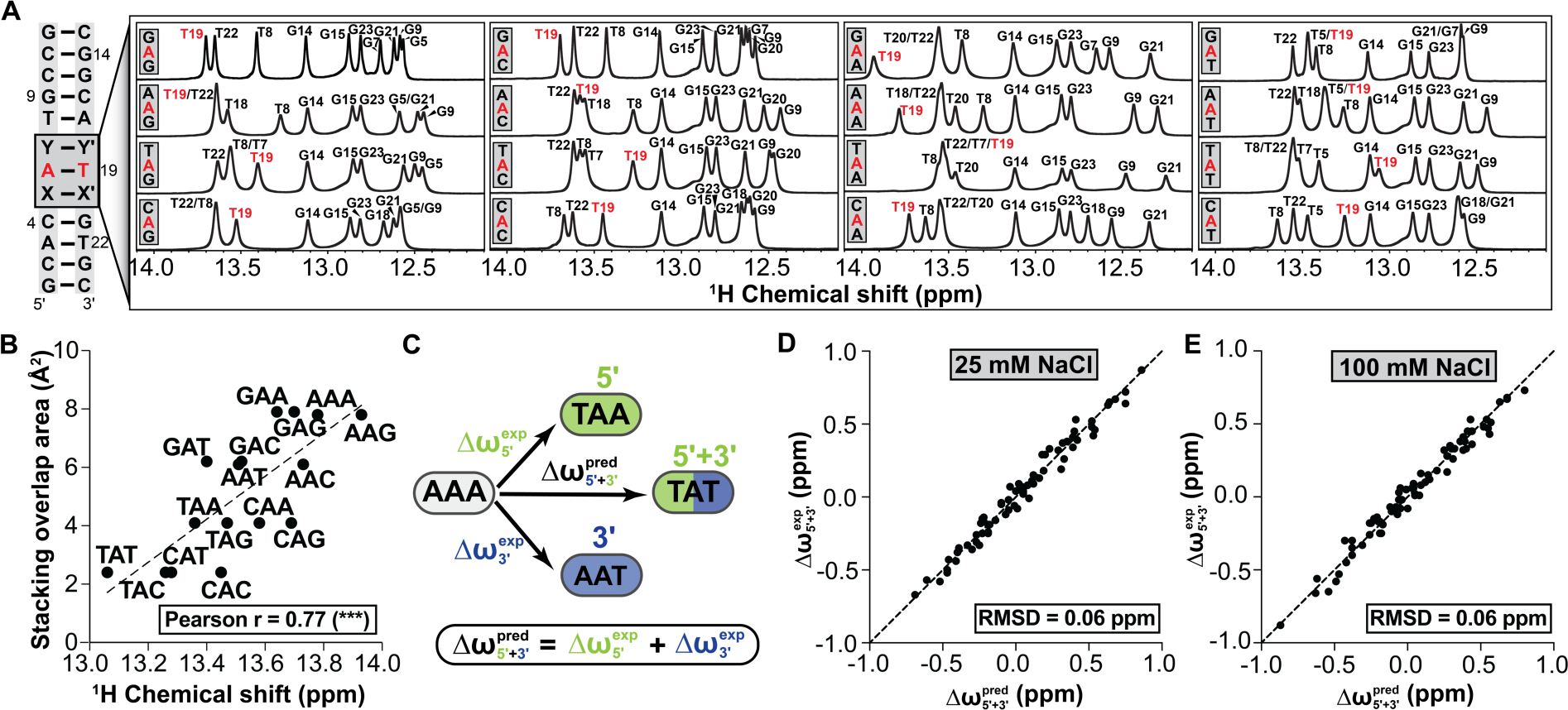
Imino ^1^H chemical shifts depend on the DNA trinucleotide sequence context. (A) The DNA constructs used in this study to examine the sequence dependence of A-T Hoogsteen dynamics. The central A-T bp of interest is highlighted in red. X-X’ and Y-Y’ represent the four Watson-Crick (A-T, T-A, G-C and C-G) neighbors. Also shown are the 1D ^1^H spectra measured at 25 mM NaCl, pH 6.8 and T=25°C. The trinucleotide sequence context is given as 5’-XAY-3’ (bottom to top) and spectra grouped according to the 5’ dinucleotide step. (B) Correlation plot between ^1^H chemical shift and the stacking overlap between the central A-T bp in the Watson-Crick conformation with its neighboring Watson-Crick bps (Methods). The Pearson correlation coefficient (r) and its statistical significance represented by the two-tailed p-value = 0.0005, are shown in the inset. (C) Representative mutational cycle used for testing additivity in the imino ^1^H chemical shifts. Starting from a reference sequence AAA, the 5’ and 3’ neighbors are changed individually to generate 5’ and the 3’ mutants TAA and AAT, respectively. Δω^exp^_5’+3’_ is the experimentally observed change in chemical shift when altering both the 5’ and the 3’ neighbors to generate TAT relative to the AAA reference. (D) Correlation plot between the predicted (Δω^pred^_5’+3’_) and experimentally observed (Δω^exp^_5’+3’_) ^1^H chemical shifts in 25 mM NaCl, pH 6.8 and (E) 100 mM NaCl, pH 8.0 at T=25°C, computed using 72 independent mutational cycles. The root mean square deviation (RMSD) between Δω^exp^_5’+3’_ and Δω^pred^_5’+3’_ is computed with respect to the y=x line.

No prior study has systematically examined how the NMR chemical shift of imino proton of a given bp varies with the identity of the neighboring bps in Watson-Crick DNA duplexes. The sequence-dependence of the T-H3 imino chemical shifts are of great interest as they could provide insights into the sequence-dependence of the ground-state Watson-Crick conformational ensemble (79,80) and could potentially even carry information concerning the propensities to form Hoogsteen bps.

Indeed, we found that changing the identity of the Watson-Crick neighbor resulted in large changes (up to ∼0.87 ppm) in the central T-H3 imino chemical shift (Figure 2A). The T-H3 chemical shifts depended on both the 5’ and 3’ neighbors (Figure 2A). T-H3 was downfield shifted when both the 5’ and 3’ neighbors were purine bases (AAA, AAG, GAA and GAG) whereas it was upfield shifted when both neighbors were pyrimidines (CAC, CAT, TAC and TAT). These significant sequence-specific variations in chemical shifts could arise from sequence-specific changes in the Watson-Crick conformational ensemble of the DNA duplex (79,80). However, similar sequence-specific variations have previously been reported for U-H3 in A-form RNA duplexes and were attributed to differences in sequence-specific ring-current contributions from the neighboring bps (79–82). Consistent with a ring current contribution, we observed a statistically significant correlation (Pearson r = 0.77) between the T-H3 chemical shift and the overlap area (methods) between the Watson-Crick A-T bp and its neighbors (Figure 2B).

To further dissect the sequence-dependence of the Watson-Crick T-H3 chemical shifts, we tested whether the variations arising from changing the 5’ or 3’ neighboring bp are independent of one another and additive, as might be expected for ring current contributions (81), or whether there are non-additive contributions possibly linked to sequence-specific changes in the Watson-Crick conformational ensemble. In particular, we asked whether the perturbation in the T-H3 chemical shift Δω^exp^_5’+3’_ relative to a reference sequence context (AAA) due to changes in the 5’ and 3’ neighbors (e.g. AAA→ TAT) can be decomposed into the sum (Δω^pred^_5’+3’_) of the individual perturbations arising from changing either the 5’ (AAA→TAA, Δω^exp^_5’_) or 3’ neighbor (AAA→TAA, Δω^exp^_3’_) (Figure 2C) (81).

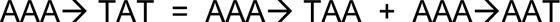

and

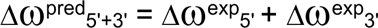

Thus, we compared Δω^pred^_5’+3’_ = Δω^exp^_5’_+Δω^exp^_3’_ to the values measured experimentally (Δω^exp^_5’+3’_) across the 72 independent mutational cycles. Indeed, we observed very good agreement between Δω^pred^_5’+3’_ and Δω^exp^_5’+3’_ with root-mean-square deviation (RMSD) of 0.06 ppm (Figure 2D), which is small compared to the chemical shift span of 0.87 ppm. Similar results (RMSD = 0.06 ppm) were obtained in 100 mM NaCl (Figure 2E). These results indicated that the sequence-specific variations in ground-state Watson-Crick T-H3 chemical shifts are likely dominated by ring current contributions (79–82). However, our analysis does not rule out sequence-specific changes in the Watson-Crick conformational ensemble which are not sensed by T-H3 or that involve the short-lived lowly-populated conformational states such as the Hoogsteen bps, which are invisible to the ground-state chemical shifts. Our sequence-specific database of T-H3 chemical shifts could facilitate assignment of imino resonances in other DNA duplexes and aid the development and application of structure-based approaches for predicting DNA chemical shifts(83,84).

### Measuring A-T Hoogsteen dynamics using ^1^H CEST

To quantitatively and systematically measure how the propensities to form the low-abundance short-lived A-T Hoogsteen bp vary with the trinucleotide sequence context, we measured ^1^H CEST profiles for all sixteen unlabeled DNA duplexes (∼1.0 mM) (Figure S2A). The ^1^H CEST experiments (methods) were performed at 600, 800 and 900 MHz NMR spectrometers equipped with cryogenic probes at T= 25°C and 25 mM NaCl and pH = 6.8 unless indicated otherwise. To minimize NOE contributions, we restricted the offset of the applied radio frequency (RF) field to −6 ppm – 6 ppm as described previously (61). Similar results were obtained when measuring the ^1^H CEST profiles at 25 versus 100 mM NaCl (Figure S2 and Table S1-S3).

For ten duplexes (AAA, AAC, AAG, CAC, CAG, CAT, GAC, GAG, GAT and TAC), the T-H3 imino resonance of the central A-T bp was well resolved and suitable for ^1^H CEST measurements (Figure 2A). Based on prior studies (54,61,62), we anticipated observing Hoogsteen exchange using ^1^H CEST for all ten sequence contexts. Indeed, for AAA, AAC, AAG, CAT, and GAT the ^1^H CEST profiles showed evidence for the expected Hoogsteen chemical exchange based on the statistically significant improvement in the fit when including 2-state chemical exchange (Figure 3A, 3B and S2). The fit yielded an upfield shifted (Δω = ω_HG_ – ω_WC_ = −1.7 – −1.2 ppm) T-H3 chemical shift (Table S1 and S3) for the minor state consistent with an A-T Hoogsteen bp (61). The population of the Hoogsteen bps (*p*_B_ = 0.17-0.47%) and rates of exchange (*k*_ex_ = *k*_forward_ + *k*_backward_ = 4000-8000 s^-1^, Table S1-S3) also fell within the range previously reported for transient A-T Hoogsteen bps (54).

**Figure 3.**
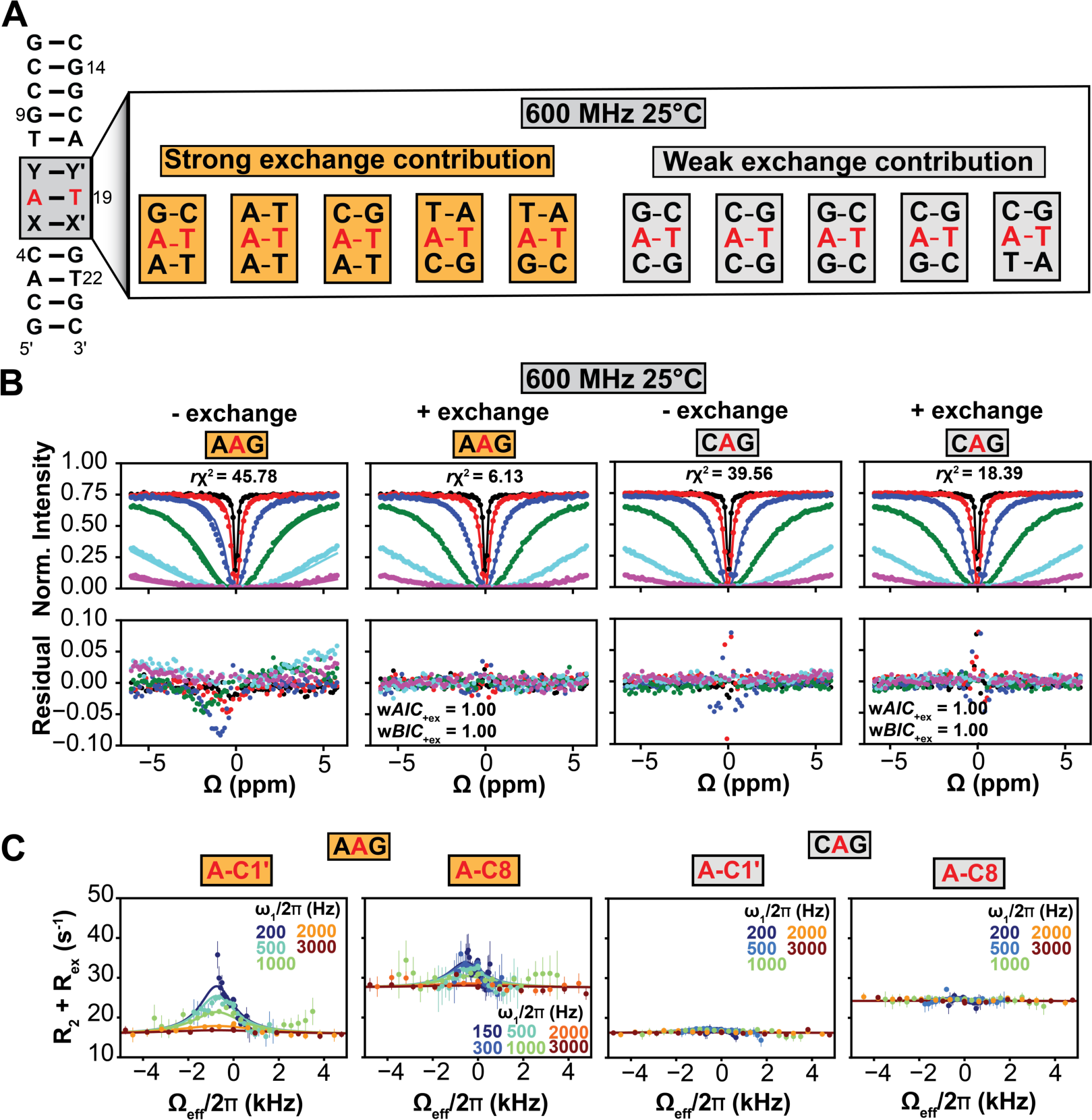
Measuring A-T Hoogsteen exchange using ^1^H CEST. (A) Trinucleotide sequence contexts showing strong and weak exchange contributions to the ^1^H CEST profiles measured at 25 mM NaCl, pH 6.8 and T=25°C at 600 MHz are indicated in orange and grey, respectively. (B) Representative ^1^H CEST profiles showing strong (AAG, left) and weak (CAG, right) exchange contributions. The solid lines represent the fit of the data to the Bloch-McConnell equations (94,95) with and without exchange, respectively. The bottom panels show the corresponding residuals of the fit. Also shown are the reduced chi-squares (*rχ*^2^), the Akaike (w*AIC*_+ex_) and Bayesian information weights (w*BIC_+ex_*) for the fits with exchange (96,97). The radio frequency powers used in the ^1^H CEST experiments are color-coded 30 Hz (black), 90 Hz (red), 270 Hz (blue), 810 Hz (green), 2430 Hz (cyan) and 5000 Hz (purple). The error bars for the ^1^H CEST data are obtained from the reference no RF irradiation experiment as described in Methods. The error bars are smaller than the data points. (C) Off-resonance C8 and C1’ ^13^C *R*_1ρ_ profiles for the central adenine in AAG (left) and CAG (right) sequence contexts measured at 25 mM NaCl, pH 6.8 and T=25°C at 800 and 700 MHz, respectively. The ^13^C labeled adenine is highlighted in red. The solid lines represent the fit of the ^13^C *R*_1ρ_ data to 2-state exchange in the Bloch-McConnell equations (94,95). The error bars represent the experimental uncertainty in the *R*_1ρ_ data estimated using a Monte-Carlo scheme as described previously (57).

Surprisingly, for CAC, CAG, GAC, GAG, and TAC, the inclusion of 2-state chemical exchange minimally improved the quality of the fit to the ^1^H CEST profiles as assessed by visual examination of the residuals (Figure 3B and S2). In addition, the 2-state fit to these data consistently yielded high *k*_ex_ > 10,000 s^-1^ values. We verified these results by ^13^C/^15^N-labeling four duplexes containing CAC, CAG, GAC, and GAG and performing off-resonance ^13^C *R*_1ρ_ experiments on the adeninine-C8 and/or C1’ (Figure 3C and S3A). We observed broad or nearly flat *R*_1ρ_ profiles characteristic of fast exchange kinetics (Figure 3C and S3A). As a positive control, the exchange parameters from ^13^C *R*_1ρ_ measurements were in agreement with those measured using ^1^H CEST for AAG sequence context (differences in exchange parameters < 2-fold, Table S1 and S4) (Figure 3C).

Finally, for six sequence contexts (CAA, TAA, TAG, TAT, AAT, and GAA), we were unable to record ^1^H CEST profiles for the central A-T bp due to spectral overlap. We were able to obtain data for CAA based on the ^1^H CEST profile measured for T20 in the AAA duplex, which was offset from the center by one bp. For TAA, TAG and TAT, we performed off-resonance ^13^C *R*_1ρ_ measurements on DNA duplexes in which the central adenine residue was ^13^C/^15^N-labeled (Figure S3B). In all three cases, we observed relaxation dispersion consistent with Hoogsteen dynamics (Figure S3B). 2-state fitting of the *R*_1ρ_ profiles yielded the downfield shifted C8 and C1’ (Δω = 2-3 ppm) (Table S4) expected for adenine in the *syn* Hoogsteen conformation (62). The exchange kinetics for TAG, which has one G-C neighbor was also comparatively fast with *k*_ex_ =18,000 ± 1,800 s^-1^ (Table S4).

### Fast Hoogsteen dynamics observed robustly in GC-rich contexts

Prior measurements of A-T Hoogsteen dynamics were performed on predominantly AT-rich sequences (54,62). However, our results revealed that when the A-T bp is flanked by G-C neighbors, the dynamics can be substantially faster. Indeed, four of the five duplexes (CAC, CAG, GAC, and GAG) with unusually fast exchange kinetics had both neighbors as G-C bps while the fifth (TAC) had one G-C neighbor. These GC-rich sequence contexts are also predicted (85) to be the thermodynamically most stable Watson-Crick conformations (Figure S6A). Interestingly, in one prior study, fast A-T Hoogsteen exchange kinetics (8,200 ± 1,000 s^-1^) was also reported for the TAC sequence context (54). These findings raised the possibility that A-T Hoogsteen dynamics are intrinsically faster when flanked by G-C neighbors.

To test this possibility, we embedded the TAG, GAG and CAC triplets into two distinct hairpins (Figure 4A). Once again, for GAG and CAC, we could not detect an exchange contribution to the ^1^H CEST profiles measured at 600 or 700 MHz and T=25°C (Figure 4B) while the ^13^C *R*_1ρ_ profiles measured for TAG yielded fast exchange kinetics with *k*_ex_ > 15,000 s^-1^ (Figure 4C).

**Figure 4.**
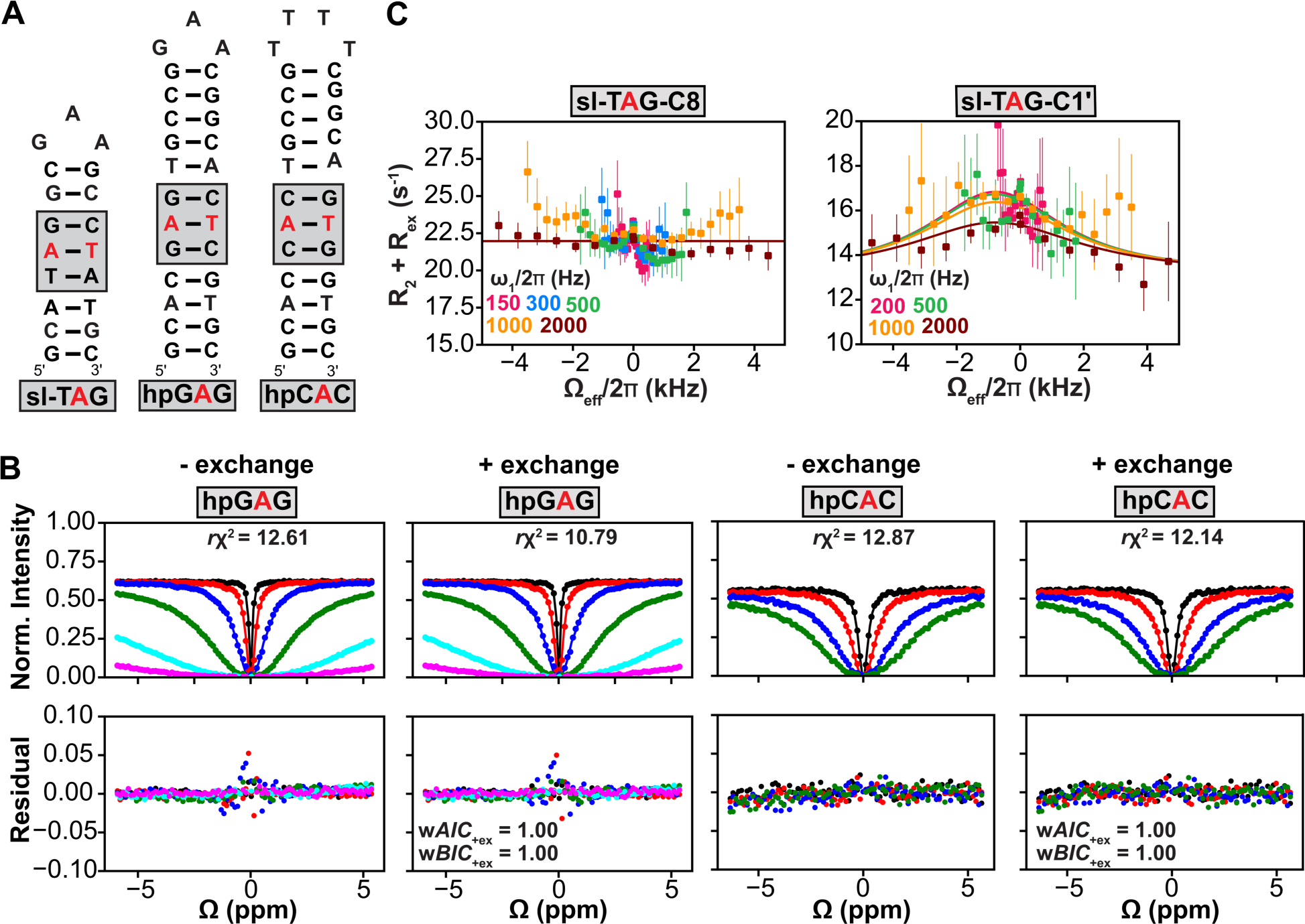
Testing robustness of fast A-T Hoogsteen exchange in GC-rich triplets. (A) Hairpin constructs with TAG, GAG and CAC trinucleotides. (B) ^1^H CEST profiles measured for hpGAG (left) and hpCAC (right).The solid lines represent the fit of the data to the Bloch-McConnell equations (94,95) with and without exchange, respectively. The bottom panels show the corresponding residuals of the fit. Also shown are the reduced chi-squares (*rχ*^2^), the Akaike (w*AIC*_+ex_) and Bayesian information weights (w*BIC_+ex_*) for the fits with exchange (96,97). The radio frequency powers used in the ^1^H CEST experiments are color-coded 30 Hz (black), 90 Hz (red), 270 Hz (blue), 810 Hz (green), 2430 Hz (cyan) and 5000 Hz (pink) for hpGAG and 100 Hz (black), 250 Hz (red), 600 Hz (blue) and 1000 Hz (green) for hpCAC. The error bars for the ^1^H CEST data are obtained from the reference no RF irradiation experiment as described in Methods. The error bars are smaller than the data points. (C) Off-resonance C8 and C1’ ^13^C *R*_1ρ_ profiles for the central adenine in sl-TAG hairpin measured at 25 mM NaCl, pH 6.8 and T=25°C at 800 and 700 MHz, respectively. The ^13^C labeled adenine is highlighted in red. The solid lines represent the fit of the ^13^C *R*_1ρ_ data to 2-state exchange in the Bloch-McConnell equations (94,95). The error bars represent the experimental uncertainty in the *R*_1ρ_ data estimated using a Monte-Carlo scheme as described previously (57).

### Resolving fast Hoogsteen dynamics in GC-rich contexts

For the five sequence contexts (CAC, CAG, GAC, GAG, and TAC) exhibiting fast exchange kinetics, we were able to better resolve the Hoogsteen exchange contribution by changing the NMR chemical shift timescale. We accomplished this by performing ^1^H CEST experiments at higher magnetic fields and/or by lowering the temperature (7-20°C). For CAC, CAG, GAG, and TAC, we resolved the Hoogsteen contribution at 900 MHz (Figure 5A and S4). The 2-state fit of the ^1^H CEST profiles yielded upfield shifted chemical shifts (Δω = - 0.9 - −2.3 ppm) (Table S1) consistent with a Hoogsteen bp. Based on a 2-state fit of the ^1^H CEST profiles, the Hoogsteen exchange rate in these sequence contexts was at least ∼5-fold faster relative to other sequences (Table S1-4).

**Figure 5.**
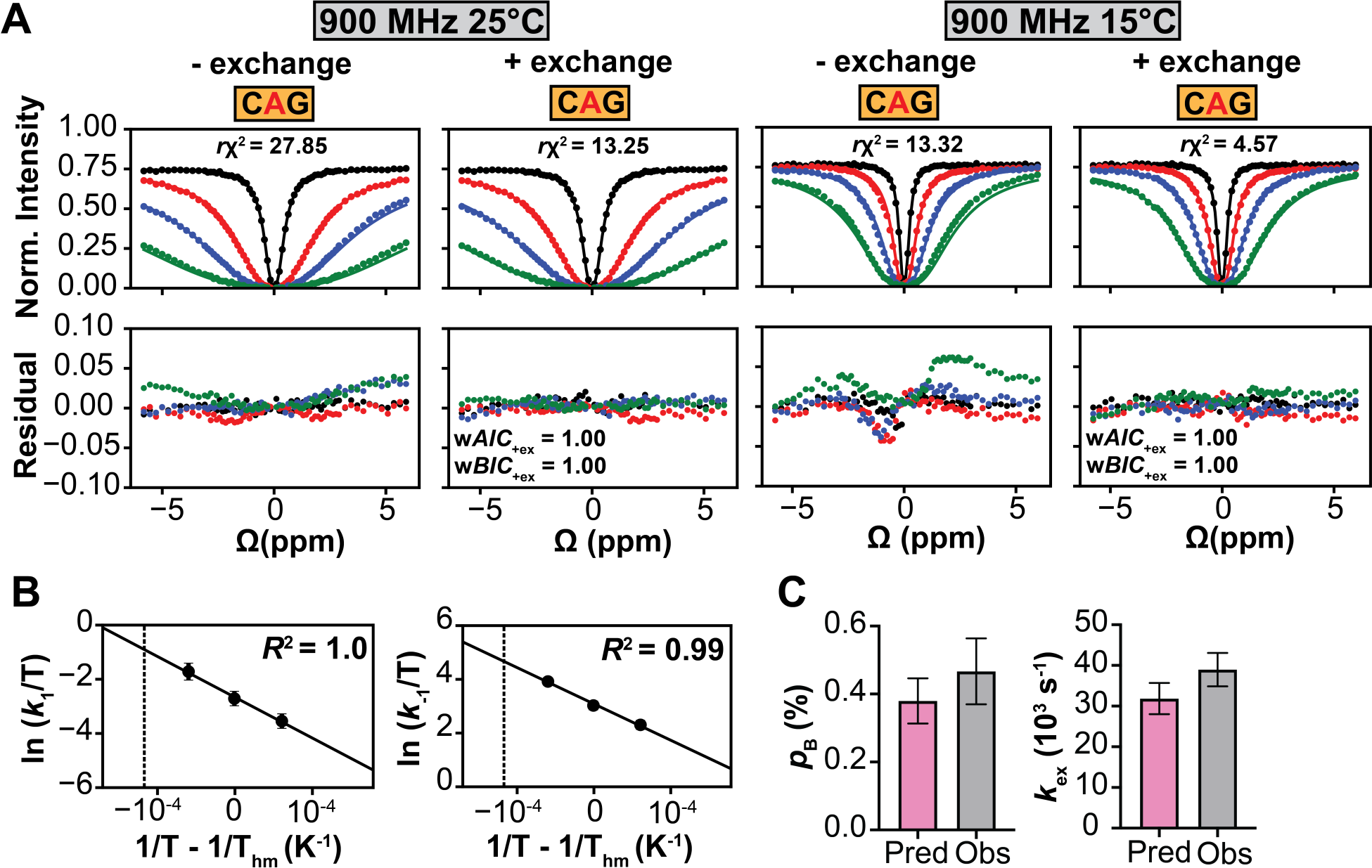
Resolving fast exchange in GC-rich sequences. (A) ^1^H CEST profile for CAG at 900 MHz and T=25°C and T=15°C. The radio frequency powers for T=25°C are color-coded 250 Hz (black), 1000 Hz (red), 2000 Hz (blue) and 4000 Hz (green) and for T= 15°C 100 Hz (black), 250 Hz (red), 500 Hz (blue) and 1000 Hz (green). The error bars for the ^1^H CEST data are obtained from the reference no RF irradiation experiment as described in Methods. The error bars are smaller than the data points. (B) Temperature dependence of forward (*k*_1_) and backward (*k*_-1_) rate constants for the Watson-Crick to Hoogsteen exchange measured for CAG at 900 MHz (Methods). The interpolated rate constants at T=25°C are indicated with a dotted line. The *R*^2^ denotes coefficient of determination. The errors bars are obtained by propagating the uncertainty in the exchange parameters for various temperatures. (C) Comparison of the Hoogsteen population (*p*_B_) and exchange rate (*k*_ex_) for CAG at T=25°C predicted (Pred) based on temperature interpolation (Methods) and those measured (Obs) from ^1^H CEST measurements performed at T=25°C at 900MHz. The uncertainty in the predicted parameters is obtained by propagating the errors in the temperature dependent rate constants *k*_1_ and *k_-_*_1_. The uncertainty in the observed parameters (Obs) represent the fitting error of the ^1^H CEST data calculated as described previously (61,98).

We independently obtained the exchange parameters for CAG, CAC, and TAC by interpolating the kinetic rate constants measured at lower temperatures to room temperature using the modified van’t Hoff equation (61,62) (Figure 5B and S4D and Table S2) and for CAG also by measuring ^13^C *R*_1ρ_ at 15°C (Figure S4A-B). In all cases, the fitted exchange parameters were within 2-fold of those obtained using ^1^H CEST at high field (Figure 5C and S4 and Table S1-3). Thus, for these GC-rich sequence contexts, the exchange contribution was comparatively smaller relative to other sequence contexts at T=25°C and 600 MHz primarily due to at least ∼5-fold faster exchange kinetics and in the case of GAC also due to the low population (∼0.05%) of the A-T Hoogsteen bp.

### Sequence-dependence of A-T Hoogsteen dynamics

Figure 6A summarizes the sequence-dependent population (*p*_B_) and rate of exchange (*k*_ex_) for the central A-T bp. Data is shown as fold difference relative to the AAA reference sequence. Even when eliminating the GAC data point which has comparatively high experimental uncertainty, the *p*_B_ and *k*_ex_ varied at least ∼4-fold and 16-fold respectively which is well within the detection limit of the ^1^H CEST and ^13^C *R*_1ρ_ experiments. We verified the statistical significance of these sequence-specific differences using a cross-fit analysis by looking for a poorer fit to the ^1^H CEST data for a sequence **i** when fixing one of the exchange parameters (*p*_B_ or *k*_ex_) to the best-fit values obtained for another sequence **j** (Figure S5 and Methods).

The A-T Hoogsteen dynamics especially the exchange kinetics did not solely depend on the 5’ neighbor (Figure 6A). For a given dinucleotide step, the population varied up to 2-fold depending on the 3’ neighbor while the exchange rate varied up to 14-fold. Flexible TA and CA dinucleotide steps (86) had the highest Hoogsteen populations, particularly for TAT, and exhibited the most significant kinetic dependence on 3’-neighbor (Figure 6A). In contrast, the stiffer AA and GA steps (86) had the lowest populations, and their exchange kinetics showed a weaker dependence on 3’-neighbor (Figure 6A and 6B).

**Figure 6.**
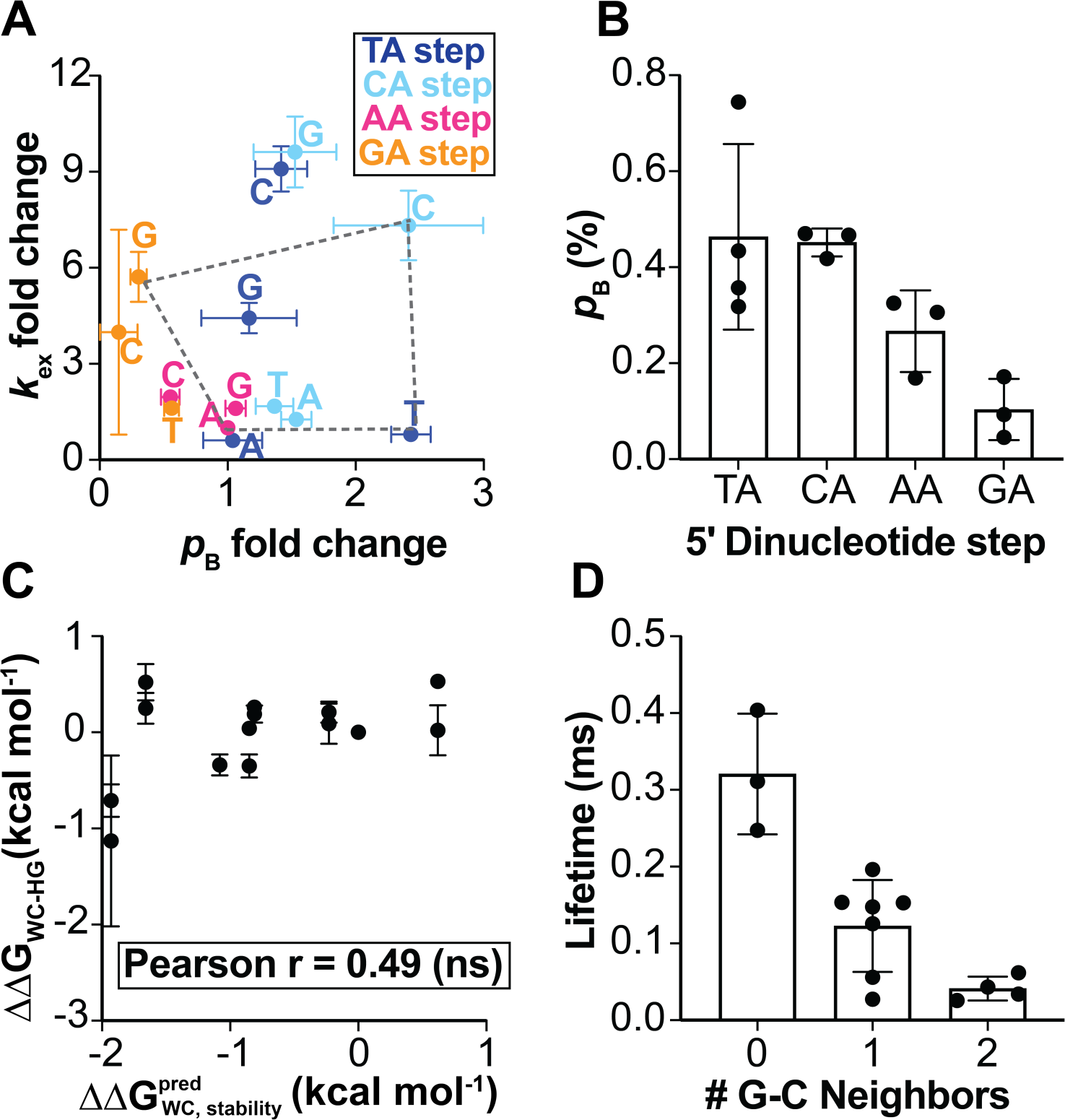
The sequence dependence of A-T Watson-Crick to Hoogsteen dynamics. (A) The population (*p*_B_) and exchange rate (*k*_ex_) measured for the A-T Hoogsteen dynamics in 14 trinucleotide sequence contexts using ^1^H CEST and off-resonance ^13^C *R*_1p_. Data represents the fold change relative to the AAA reference sequence context. Data are color-coded according to the 5’ neighbor (see inset) and the 3’ neighbor is indicated on the plot. The error bars were obtained by propagating the uncertainty in the exchange parameters. The dotted polygon encloses the sequence contexts exhibiting the extremities of the exchange parameters. (B) The average population (*p*_B_) of Hoogsteen bps for the various 5’ dinucleotide steps. The error bars represent the standard deviation of *p*_B_ values for a given dinucleotide step. (C) Correlation plot between the overall stability of the duplexes (normalized relative to AAA sequence context; ΔΔG^pred^ = ΔG^pred^ (seq) – ΔG^pred^ (AAA) predicted using the latest nearest-neighbor parameters (85) and conformational propensity to form Hoogsteen ΔΔG_WC-HG_ = ΔG_WC-HG_ (seq) – ΔG_WC-HG_ (AAA)) relative to AAA sequence context measured in this study using NMR. The error bars for ΔΔG_WC-HG_ are obtained by propagating the uncertainty in the exchange parameters. The statistical significance of the Pearson r is assessed with the two-tailed p-value (0.0781). (D) Average lifetime (1/*k*_-1_) of Hoogsteen bps as a function of number of neighboring G-C bps. The error bars represent the standard deviation of lifetime values for a given dinucleotide step.

On average, the Hoogsteen populations followed the order TA∼CA>AA∼GA (Figure 6B), which is approximately the reverse order of dinucleotide stability GA∼CA>AA>TA (87). Thus, the sequence-specific thermodynamic propensities to form the A-T Hoogsteen bp are driven in part by the sequence-specific stabilities of the A-T Watson-Crick bp, in agreement with prior study employing a more limited set of sequence contexts under a variety of buffer conditions (54,77). Despite these qualitative trends aggregated over the dinucleotide steps, the sequence-specific A-T Hoogsteen propensities for the individual triplets did not quantitatively correlate (Pearson r = 0.49, non-significant) with the sequence-specific thermodynamic stabilities of these triplets predicted using nearest neighbor rules (85,87) (Figure 6C). This difference is to be expected considering that the energetics of the Hoogsteen conformation is influenced by unique sequence-specific interactions involving the *syn* adenine base and differ from the sequence-specific energetics of the Watson-Crick bps. Interestingly, we did observe a modest correlation (Pearson r = 0.77) between the sequence-dependent T-H3 chemical shift difference between the Watson-Crick and Hoogsteen conformational states (Δω_WC-HG_) and Hoogsteen propensities. Sequence contexts with a higher propensity to form Hoogsteen bps had smaller Δω_WC-HG_ values (Figure S6B). This correlation could be leveraged in the future to accelerate assessment of sequence-dependent Hoogsteen energetics.

As noted above, GC-rich sequence contexts exhibited surprisingly fast A-T Hoogsteen exchange kinetics due to a much faster backward rate. Indeed, across all sequence contexts examined, the lifetime of the A-T Hoogsteen bp with two G-C neighbors (CAC, CAG, GAC, GAG) was on average at least ∼3-fold shorter than sequences (AAC, AAG, CAT, GAT, TAC, TAG) with one G-C neighbor; which in turn was at least ∼3-fold shorter than the sequences without G-C neighbors (AAA, TAA, TAT) (Figure 6D). Future studies should investigate the A-T Hoogsteen conformation in GC-rich sequence contexts in which it could have distinct structural characteristics such as weaker h-bonds that could explain the faster backward rate to form the Watson-Crick conformation.

### Dependence on monovalent ion concentration and pH

Prior studies showed that the A-T Hoogsteen dynamics are largely independent of pH over the range 5.3 - 7.6 (67,77,83,88) and salt concentration (77). To examine the impact of varying pH and salt concentration on sequence-specific Hoogsteen dynamics, we repeated the ^1^H CEST experiments (Figure 7A and S2B) when simultaneously increasing the monovalent ion concentration from 25 mM to 100 mM NaCl and the pH from 6.8 to 8 and using a buffer that mimics the conditions typically used in optical melting experiments (77,85).

**Figure 7.**
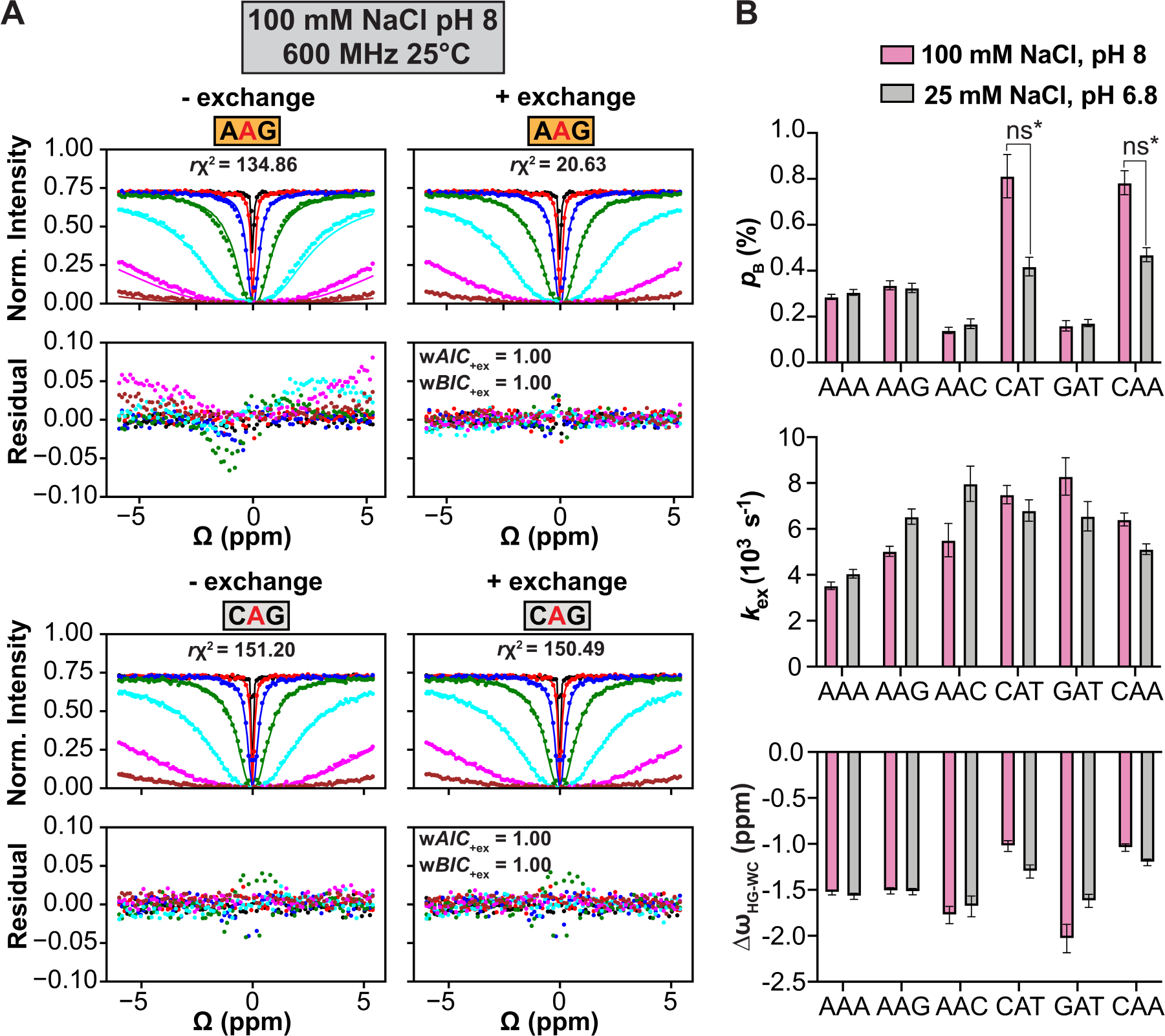
Examining the dependence of sequence-specific A-T Hoogsteen dynamics on monovalent ion concentration and pH. Representative ^1^H CEST profile for AAG (left) and CAG (right) measured at 100 mM NaCl, pH 8 and T=25°C at 600 MHz. The solid lines represent the fit of the data to the Bloch-McConnell equations (94,95) with and without exchange. The bottom panels show the corresponding residuals of the fit. Also shown are the reduced chi-squares (*rχ*^2^), the Akaike (w*AIC*_+ex_) and Bayesian information weights (w*BIC_+ex_*) for the fits with exchange (96,97). The radio frequency powers used in the ^1^H CEST experiment are color-coded 10 Hz (black), 30 Hz (red), 90 Hz (blue), 270 Hz (green), 810 Hz (cyan), 2430 Hz (pink) and 5000 Hz (brown). The error bars for the ^1^H CEST data are obtained from the reference no RF irradiation experiment as described in Methods. The error bars are smaller than the data points. (B) Shown are the population (*p*_B_) exchange rate (*k*_ex_) and chemical shift difference (Δ**ω**) for six sequence contexts where Hoogsteen exchange parameters were extractable for both 25 and 100 mM NaCl datasets from T=25°C measurements. The error bars represent the uncertainty in exchange parameters (61,98). The statistical significance of the *p*_B_ for CAT and CAA (ns*) is assessed based on the cross-fit analysis for the datasets for the two salt conditions. As described in Methods. All ^1^H CEST experiments were performed at 600 MHz.

For six out of eleven trinucleotide sequence contexts on which ^1^H CEST measurements were possible, changing the salt concentration and pH minimally impacted (differences in *p*_b_ and *k*_ex_ <2-fold) the Hoogsteen dynamics (Figure 7B and S2B). For two sequence contexts (CAA and CAT), the differences in *p*B were ∼1.7-1.9 fold, however based on our cross-fit analysis (methods) these differences were not statistically significant. In addition, for the remaining five GC-rich trinucleotide sequence contexts, the fast Hoogsteen exchange kinetics was preserved in 100 mM NaCl and pH 8 (Figure S2B). Therefore, for the eleven sequence contexts examined in this study, the monovalent ions do not appear to preferentially interact with and stabilize the Watson-Crick versus Hoogsteen conformation and the transition state is likely not protonated. These results underscore the utility of the ^1^H CEST experiment in mapping the dependence of DNA dynamics on environmental conditions.

## Discussion

Depending on the trinucleotide sequence context, the A-T Hoogsteen population and exchange rate varied 4-fold and 16-fold, respectively. Therefore, the binding affinities of proteins that employ an A-T Hoogsteen bp to bind DNA can vary 4-fold (∼0.9 kcal/mol at T=25°C) with sequence purely due to the sequence-specific propensities to form the Hoogsteen conformation and potentially sixteen fold for proteins that bind to tandem Hoogsteen bps (63). The kinetic rates of biochemical processes in which forming an A-T Hoogsteen bp is rate-limiting could vary by up to 23-fold (∼1.9 kcal/mol at T=25°C) depending on trinucleotide sequence context due to the inherent differences in Hoogsteen exchange kinetics. These sequence-specific A-T Hoogsteen dynamics could shape the sequence-dependent kinetics of DNA-protein association (64,69) as well as DNA damage (76,89). The Hoogsteen dynamics did not strongly depend on the monovalent ion concentration and pH in agreement with a prior studies employing *R*_1ρ_ to examine the impact of monovalent and pH on A-T Hoogsteen dynamics (67,77,83,88).

The sequence-specific propensities to form A-T Hoogsteen bps differed from the sequence-specific thermodynamic stabilities of A-T Watson-Crick bps (Figure 6C). We also did not observe any statistically significant correlation (Pearson r = 0.40) between the propensity to form the A-T Hoogsteen bps and the predicted sequence-specific Watson-Crick minor groove width (90), another key characteristic of the ground-state (Figure S6C). These results highlight how the propensities to form alternative DNA conformational states with unique stacking interactions can have markedly different dependencies on sequence.

We recently reported the ‘delta-melt’ approach for measuring propensities to form alternative rare conformations such as Hoogsteen bps using optical melting experiments (77). In delta-melt, chemical modifications are used to mimic the rare conformational state, and the conformational propensities are inferred from the difference between the melting energetics of the modified duplex versus its unmodified counterpart. The relative Hoogsteen propensities measured across seven overlapping triplets using delta-melt for which Hoogsteen population could be determined with high confidence here using ^1^H CEST and ^13^C *R*_1ρ_ were within experimental uncertainty (Figure S6D). While delta-melt can potentially be performed in much higher throughput and lower cost, the higher precision of the ^1^H CEST NMR experiment makes it better suited for characterizing small (< 1 kcal/mol) sequence-specific contributions to conformational propensities. The ^1^H CEST experiment also provides kinetic information inaccessible from delta-melt.

Our results revealed that the lifetimes of the A-T Hoogsteen bps are especially short for the thermodynamically stable GC-rich sequence contexts. It could be that the G-C neighbors more substantially stabilize the Watson-Crick ground-state and transition state relative to the Hoogsteen conformation, exhibiting an early transition state unlike that of the A-T rich sequences (54). Another possibility is that in these sequence contexts the Hoogsteen dynamics proceeds via alternative pathways. MD simulations (91) suggest there can be multiple pathways for transitioning from the Watson-Crick to Hoogsteen conformation which differ with regards to whether the purine flips inside the helix or the entire nucleotide first flips outside, followed by purine rotation along the glycosidic bond to generate the *syn* conformation and finally flipping back in the *syn* conformation to form the Hoogsteen (91). Based on the MD simulations, it has been speculated that the transition pathway can be sequence dependent (91). The sequence-dependence of the transition pathway could be explored in the future by coupling ^1^H CEST experiments with chemical modifications and examining the impact on the forward and backward rate constants using the phi-value analysis framework (92) as has been recently done using ^13^C and ^15^N *R*_1p_ in studies of RNA dynamics (93).

Finally, while we have focused on A-T, the ^1^H CEST strategy reported here can immediately be applied to measure the sequence and pH-dependent propensities to form of G(*syn*)-C^+^ and G(*syn*)-A^+^ Hoogsteen bps (47) and to examine how these Hoogsteen dynamics vary with mismatch, damage (78), and epigenetic modifications.

## Materials and Methods

### Sample Preparation

#### DNA oligonucleotides

The 32 unlabeled DNA single strands (Figure 2A) and 2 hairpin oligonucleotides (Figure 4A) were purchased from Integrated DNA Technologies (IDT) with standard desalting purification. An additional seven (CAC, CAG, GAC, GAG, TAA, TAG, TAT) single strands where the central adenine was fully ^13^C and ^15^N labeled were purchased from Yale Keck Oligonucleotide Synthesis Facility with cartridge purification. The sl-TAG sample with the central ^13^C/^15^N labeled adenine was synthesized in-house using the MerMade12 synthesizer with DMT-on protocol and purified using the cartridge method as previously described (77).

#### Buffer preparation

The NMR buffer used in this study was composed of 15 mM sodium phosphate, 25 mM sodium chloride and 0.1mM EDTA at pH 6.8. The high salt buffer used in this study was composed of 10 mM sodium phosphate, 100 mM sodium chloride, 1 mM EDTA and 1 mM triethanolamine (TEOA) in 95:5% H_2_O:D_2_O at pH 8.8.

#### Annealing and buffer exchange

The single stranded DNA oligonucleotides were dissolved in deionized water to yield 1-2 mM single-strand stocks. The single-strands were combined in equimolar ratio using the absorbance at 260 nm and coefficients calculated using ADT Bio Oligo Calculator (https://www.atdbio.com/tools/oligo-calculator) to deduce the oligonucleotide concentration. The duplexes were annealed by heating to 95 °C for 5 min, followed by slow cooling to room temperature for >1 hour. The duplex samples were then buffer exchanged into NMR buffer using Amicon Ultra-15 centrifugal concentrators (3-kDa cutoff, Millipore Sigma) at least 3 times, to a final concentration of 1-1.5 mM. The samples were supplemented with 10% D_2_O (Millipore Sigma). Similarly duplex samples were prepared in the high salt buffer, except supplementation with D_2_O was not performed as the high salt buffer contained 5% D_2_O. The final pH of the high salt (100 mM NaCl) NMR samples measured using a thin pH electrode was ∼8.0.

To prepare the ^13^C/^15^N labeled duplex samples, the ^13^C/^15^N labeled single strands (∼0.5-0.7 mM) similarly combined with their unlabeled complements in equimolar ratio, annealed and buffer exchanged into NMR buffer.

Hairpin samples were annealed in water at low concentrations (< 50 μM) by heating at 95°C for 5 minutes and immediately placing on ice for > 30 minutes. Samples were then buffer exchanged to NMR buffer following the same protocol as duplex samples.

### NMR experiments

#### Resonance assignment

The imino protons of the unlabeled DNA duplexes were assigned using 2D [^1^H-^1^H] NOESY and [^1^H-^15^N] HSQC experiments collected on a 700 MHz Bruker Avance III spectrometer equipped with a triple resonance cryogenic probe at T=25°C. 1D spectra were processed on Topspin3.6.3 (99). 2D NMR spectra were processed using NMRpipe (100) and analyzed using SPARKY (Thomas D. Goddard and Donald G. Kneller, SPARKY 3, University of California, San Francisco).

#### ^1^H CEST experiments

##### Data collection

^1^H CEST experiments were performed on Bruker Avance III 600, 800 and 900 MHz spectrometers as described previously (61). Data were collected at T=25°C, unless stated otherwise using a relaxation delay of 100 ms except for TAC for which the relaxation delay was 80 ms. The RF powers ranged between 30-4000 Hz and the offsets spanned ± 6 ppm (Table S5 and S6). The RF powers and inhomogeneities were calibrated as previously described (61). We note that under the near neutral pH or basic conditions used in this study, the protonated G-C^+^ Hoogsteen bps are exceptionally lowly populated (<0.1%) falling below the detection limits (∼0.1%) of these NMR experiments.

##### Data analysis

The ^1^H CEST profiles were fit to a 2-state model (61). Briefly, the peak intensities as a function of RF power and offset frequency were extracted using NMRPipe (100). The peak intensity for each RF power were normalized by the average peak intensity in a reference experiment with no RF irradiation, performed in triplicate (61). The standard deviation of the peak intensity in the reference experiment was assumed to be the error in the peak intensity in the presence of RF irradiation (61). The ^1^H CEST profiles were generated by plotting the normalized intensity as a function of the offset Ω = *ω_RF_* −*ω_obs_*, in which *ω_RF_* is the angular frequency of the applied RF field and *ω_obs_* is the larmor frequency of the resonance of interest. The normalized intensities were fit to the Bloch-McConnell (B-M) equations (94,95). The quality of the 2-state fits with and without conformational exchange was compared based on the *rχ*^2^ and AIC/BIC metrics (96,97). The uniqueness of the exchange parameters was tested by examining the quality of the fit was examined as a function of varying and fixing each of the three exchange parameters (*p*_B_, *k*_ex_ and Δω). Furthermore, to assess the statistical significance of the differences in exchange parameters obtained for a pair of sequences (i, j), we performed cross-fit analysis in which we compared the quality of the fit obtained by fixing a parameter from the solution of sequence i to the ^1^H CEST data of solution j and *vice versa*. Only when the forced fits for data of sequence i with parameters of sequence j and data of sequence j with parameters of sequence i yielded poorer quality fits than their best fit solution, did we deem the sequences i and j to have statistically different Hoogsteen exchange parameters.

##### Temperature interpolation

For select sequences (CAG, CAC and TAC) the exchange parameters at T=25°C were also interpolated by fitting the forward and backward rate constants (*k*_1_ and *k*_-1_) measured at lower temperature (7-20°C) to the modified van’t Hoff equation (54,61,62):

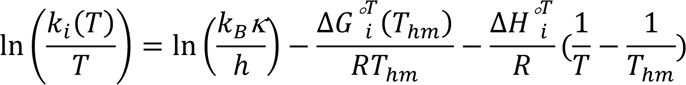

*k*_i_ (i=1, −1) are the forward and backward rate constants, respectively. ΔG°^T^ and ΔH°^T^ are the Gibbs free energy and enthalpy of activation (i=1) or deactivation (i=-1) at a temperature T. *R* is the universal gas constant, k_B_ is the Boltzmann constant, and *κ* is the transmission coefficient (assumed to be 1). *T*_hm_ is the harmonic mean of the temperatures employed. The activation (i=1) and deactivation (i=-1) (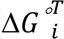) barriers were calculated using:

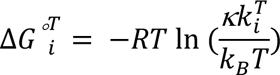

##### ^13^C R1ρ experiments

Off-resonance ^13^C *R*_1ρ_ experiments were performed using a 1D selective excitation scheme, as previously described (57). The spin lock powers (*ω*_1_/2*π*) ranged between 150 – 3000 Hz. The offsets ranged ±3.5 times the spin lock power (Table S7). ^13^C spin-locks were applied for durations optimized for ∼70% decay of the magnetization at the end of the relaxation period. The *R*_1ρ_ value for each spin-lock and offset combination was computed by fitting the decay of the magnetization to a mono-exponential curve as described previously (57,101). The experimental errors in the *R*_1ρ_ values were estimated using a Monte-Carlo approach (57). The exchange parameters were obtained by fitting the *R*_1ρ_ data to Bloch-McConnell (B-M) equations (94,95) and the uncertainty in the exchange parameters represent the standard error of the fit obtained by propagating the error in the *R*_1ρ_ values using a Monte-Carlo approach (57,101).

### Structural and energetic analysis

#### Stacking overlap area calculation

The X3DNA web server (102) was used to generate idealized B-DNA fiber models for all 16 duplexes. X3DNA DSSR was then used to compute the stacking overlap area for the central A-T bp of interest with the 5’ and 3’ neighboring Watson-Crick bps using only the aromatic ring atoms (24).

#### Minor Groove width calculation

The sequence dependent minor groove widths were calculated using the DNAshape web server (90). 5’-GCACXAYTGCCG-3’, in which X and Y were varied to generate all possible 16 trinucleotides, was used as the input sequence. The minor groove width for the 6^th^ residue was denoted as the minor groove width of the trinucleotide of interest.

#### Thermal stability prediction

The thermal stability of the 16 duplexes was computed using the latest set of nearest neighbor parameters (85) using the NUPACK web server (www.nupack.org). The two strands of the duplex at 1 *µ*M concentration were added in a tube for calculation of melting equilibria and the maximum complex size was limited to 2. dna04 parameter set with ‘All stacking’ ensemble setting was used. 0.05 M Na^+^ and 0 M Mg^2+^ ion concentrations were used for the calculations to match the experimental conditions used in NMR experiments. The Gibbs free energy of melting was predicted at T=25°C.

## Acknowledgments

We thank all members of the Al-Hashimi laboratory for their critical input. We thank Duke Magnetic Resonance Spectroscopy Center and New York Structural Biology Center (NYSBC) for NMR resources.

