## Supplementary Information for "Quantitative and systematic NMR measurements of sequence-dependent A-T Hoogsteen dynamics uncovers unique conformational specificity in the DNA double helix"

**A**

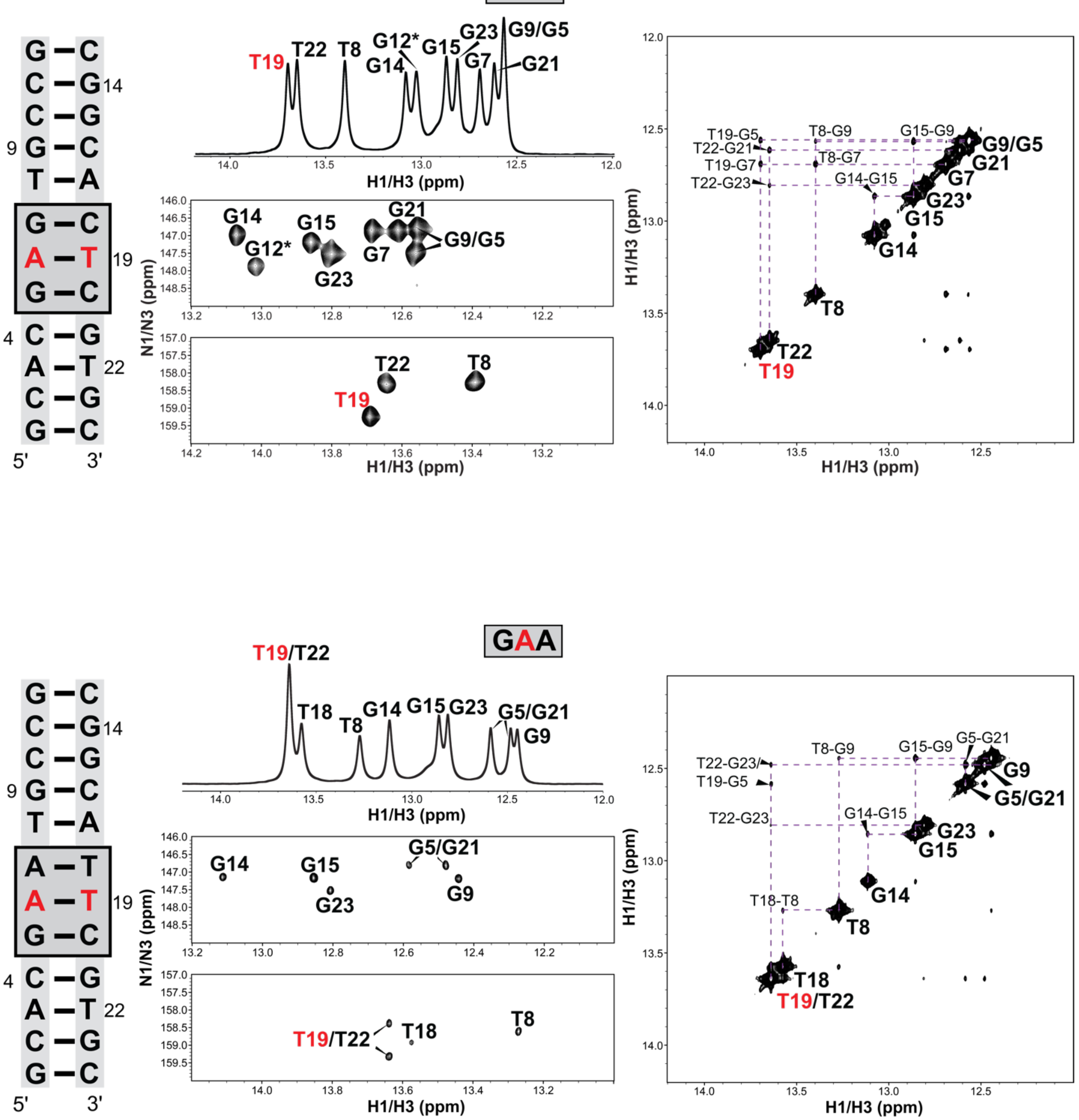

GAG
T19
T22
T8
G12*
G14
G15
G23
G9/G5
G7
G21
H1/H3 (ppm)
N1/N3 (ppm)
H1/H3 (ppm)
T19-G5
T22-G21
T19-G7
T22-G23
T8-G9
T8-G7
G15-G9
G14-G15
5'
3'
GAA
T19/T22
T18
G5/G21
G9
T22-G23/
T19-G5
T18-T8
G5-G21

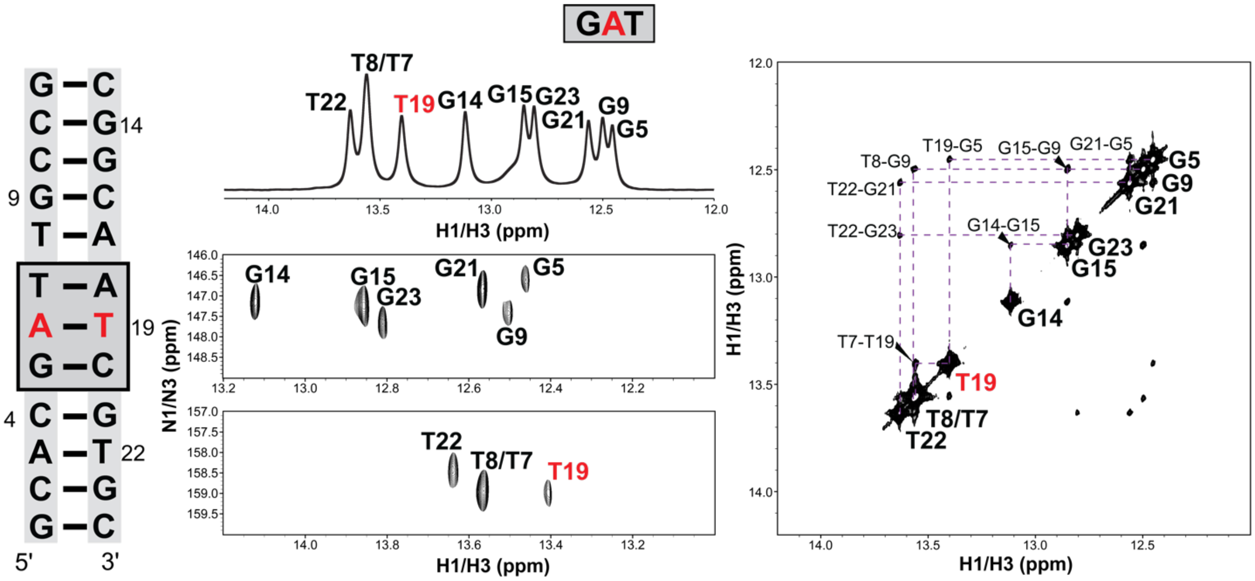
GAT
5'
3'
H1/H3 (ppm)
N1/N3 (ppm)
T8/T7
T22
T19
G14
G15
G23
G21
G9
G5
T7-T19
T8-G9
T19-G5
G15-G9
G21-G5
T22-G21
T22-G23
G14-G15

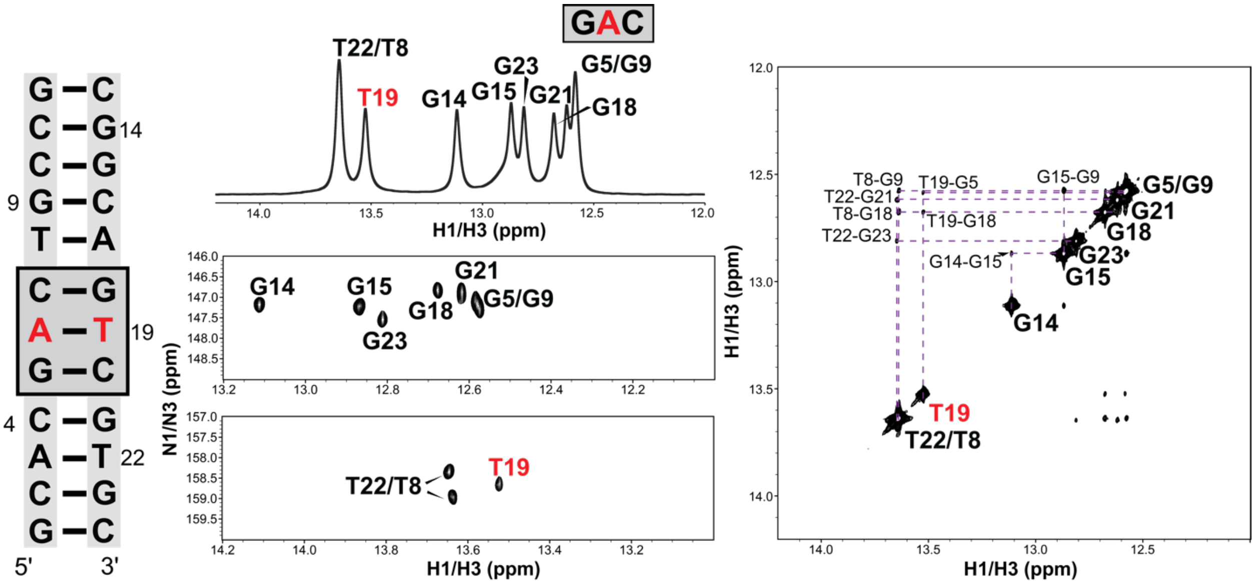
GAC
5'
3'
H1/H3 (ppm)
N1/N3 (ppm)
T22/T8
T19
G14
G15
G23
G21
G5/G9
G18
T8-G9
T19-G5
G15-G9
T22-G21
T8-G18
T19-G18
T22-G23
G14-G15

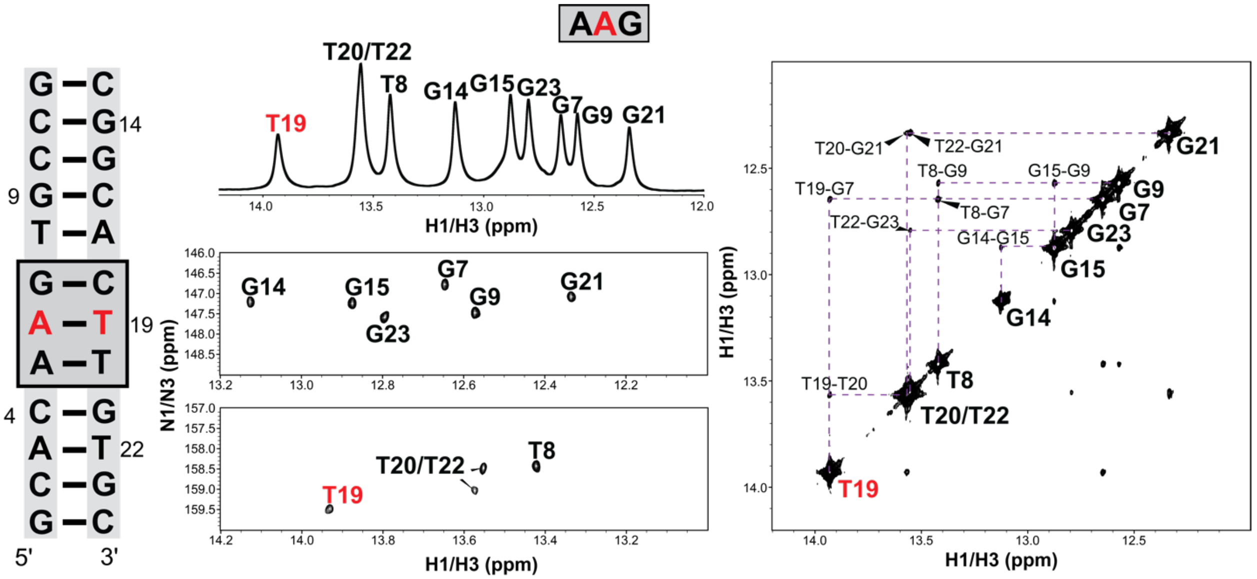
AAG
G C
C G 14
C G
9 G C
T A
G C
A T 19
A T
4 C G
A T 22
C G
G C
5'
3'
T20/T22
T8
G14
G15
G23
G7
G9
G21
T19
H1/H3 (ppm)
N1/N3 (ppm)
T20-G21
T22-G21
T8-G9
G15-G9
T19-G7
T8-G7
T22-G23
G14-G15
T19-T20
H1/H3 (ppm)

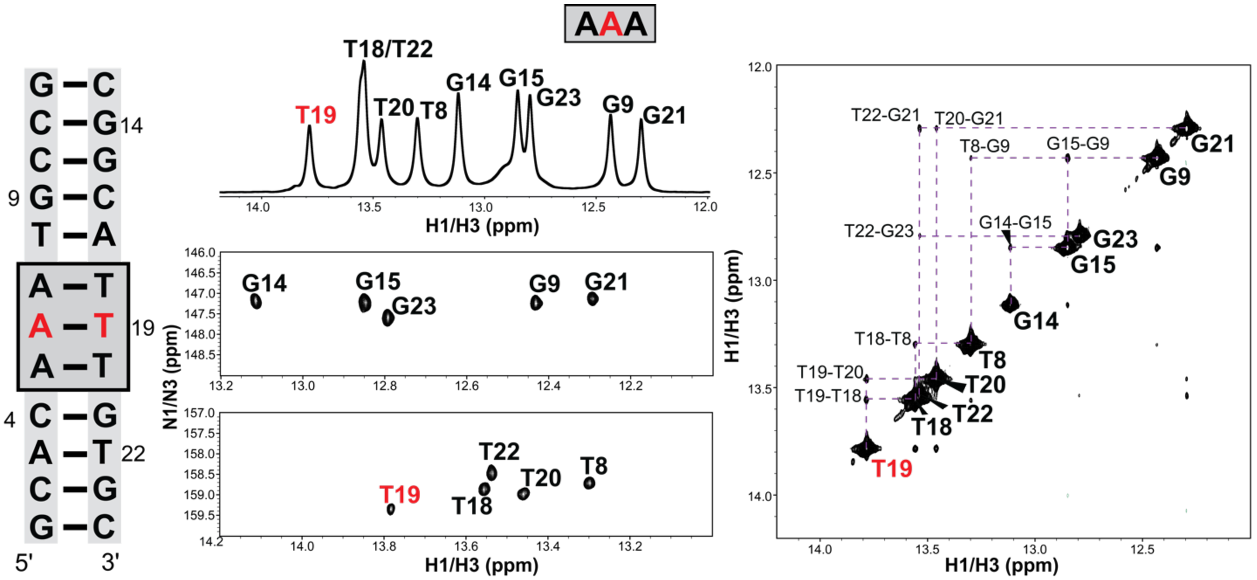
AAA
G C
C G 14
C G
9 G C
T A
A T
A T 19
A T
4 C G
A T 22
C G
G C
5'
3'
T18/T22
T19
T20
T8
G14
G15
G23
G9
G21
H1/H3 (ppm)
N1/N3 (ppm)
T22
T18
T22-G21
T20-G21
T8-G9
G15-G9
G14-G15
T22-G23
T18-T8
T19-T20
T19-T18
H1/H3 (ppm)

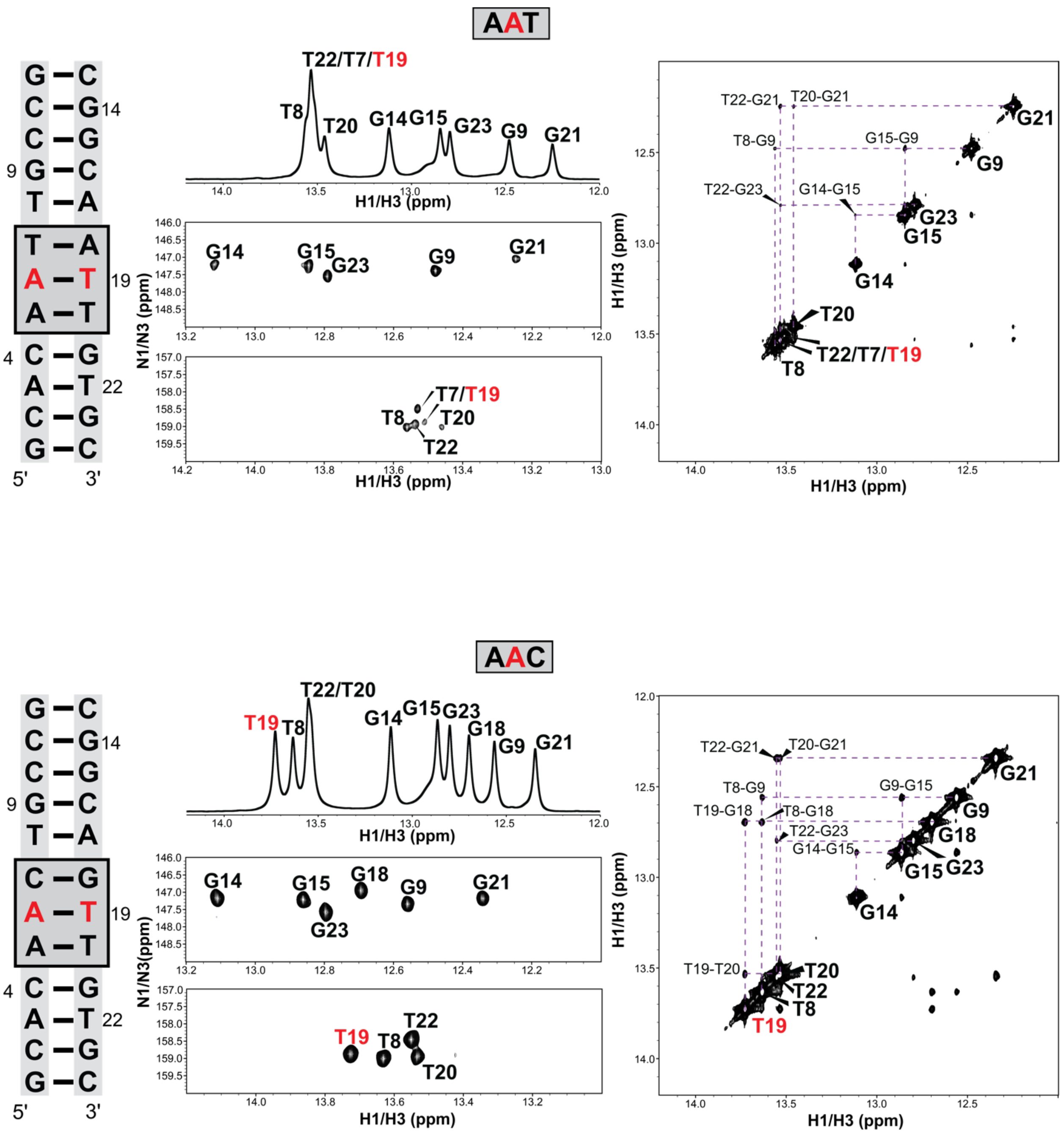

AAT
T22/T7/T19
T8
T20
G14
G15
G23
G9
G21
H1/H3 (ppm)
N1/N3 (ppm)
T7/T19
T22
T22-G21
T20-G21
T8-G9
G15-G9
T22-G23
G14-G15
T22/T7/T19
5'
3'
AAC
T22/T20
T19
G18
T22-G21
T20-G21
T8-G9
G9-G15
T19-G18
T8-G18
T22-G23
G14-G15
T19-T20

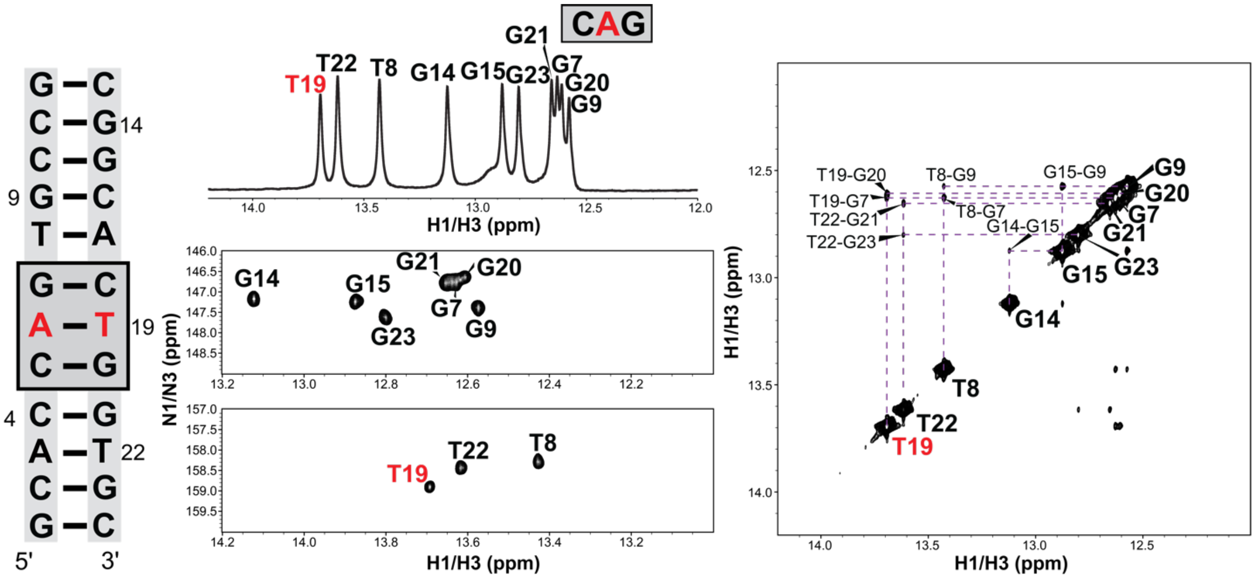
CAG
G–C
C–G 14
C–G
9 G–C
T–A
G–C
A–T 19
C–G
4 C–G
A–T 22
C–G
G–C
5'
3'
T19
T22
T8
G14
G15
G23
G21
G7
G20
G9
H1/H3 (ppm)
N1/N3 (ppm)
H1/H3 (ppm)
T19-G20
T19-G7
T22-G21
T22-G23
T8-G9
T8-G7
G15-G9
G14-G15
H1/H3 (ppm)
H1/H3 (ppm)

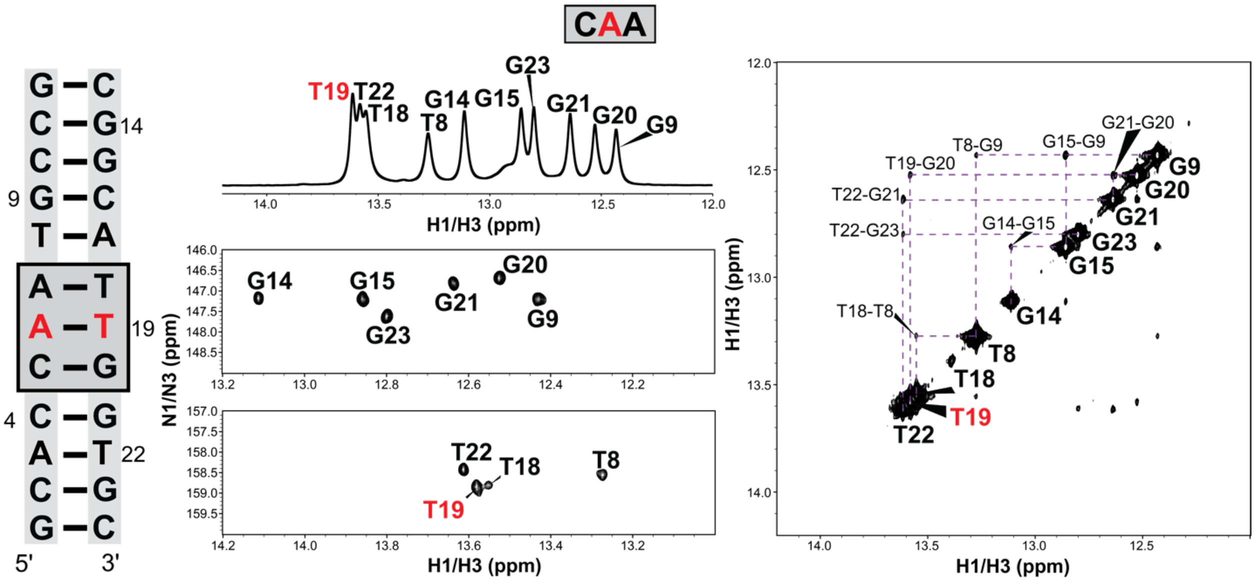
CAA
G–C
C–G 14
C–G
9 G–C
T–A
A–T
A–T 19
C–G
4 C–G
A–T 22
C–G
G–C
5'
3'
T19
T22
T18
T8
G14
G15
G23
G21
G20
G9
H1/H3 (ppm)
N1/N3 (ppm)
H1/H3 (ppm)
T19-G20
T22-G21
T22-G23
T18-T8
T8-G9
G15-G9
G21-G20
G14-G15
H1/H3 (ppm)
H1/H3 (ppm)

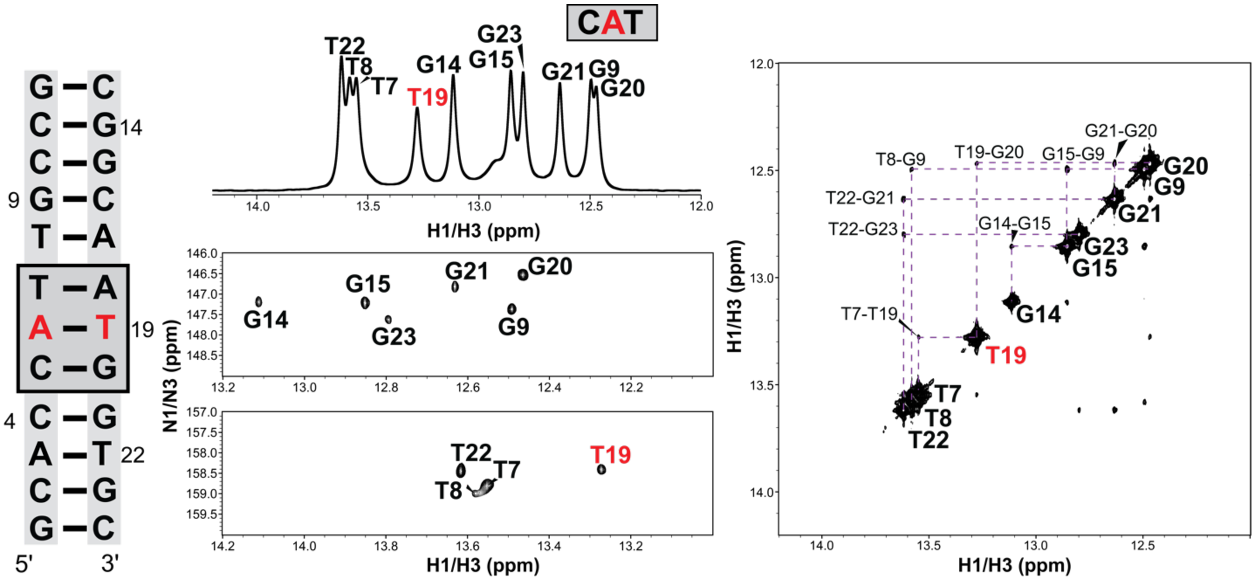
CAT
G – C
C – G 14
C – G
9 G – C
T – A
T – A
A – T 19
C – G
4 C – G
A – T 22
C – G
G – C
5'
3'
T22
T8
T7
T19
G14
G15
G23
G21
G9
G20
H1/H3 (ppm)
N1/N3 (ppm)
T8-G9
T19-G20
G15-G9
G21-G20
T22-G21
T22-G23
G14-G15
T7-T19

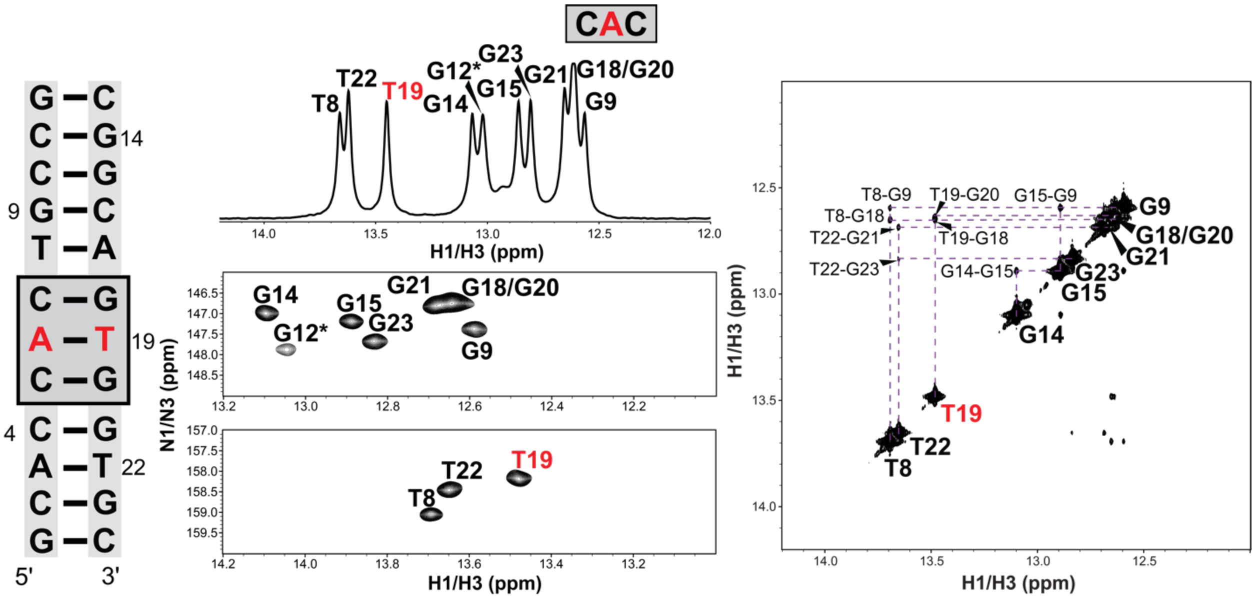
CAC
G – C
C – G 14
C – G
9 G – C
T – A
C – G
A – T 19
C – G
4 C – G
A – T 22
C – G
G – C
5'
3'
T8
T22
T19
G12*
G14
G23
G15
G21
G18/G20
G9
H1/H3 (ppm)
N1/N3 (ppm)
T8-G9
T19-G20
G15-G9
T8-G18
T22-G21
T19-G18
T22-G23
G14-G15

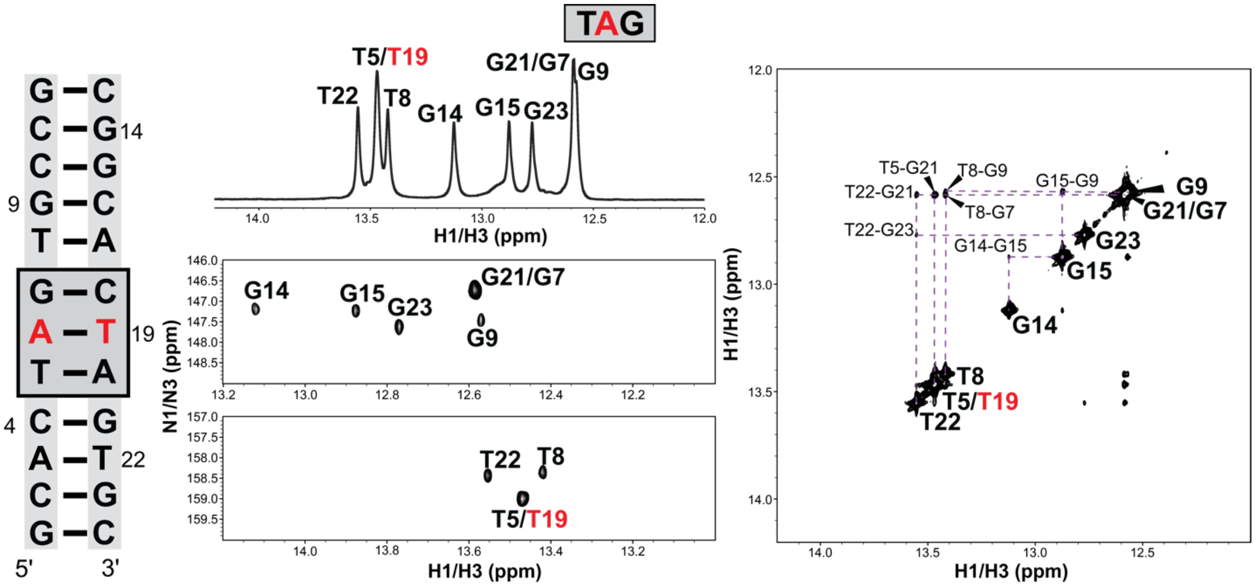
TAG
T5/T19
T22
T8
G14
G15
G23
G21/G7
G9
H1/H3 (ppm)
N1/N3 (ppm)
T5-G21
T8-G9
T22-G21
T8-G7
G15-G9
T22-G23
G14-G15
5'
3'
14
9
19
4
22

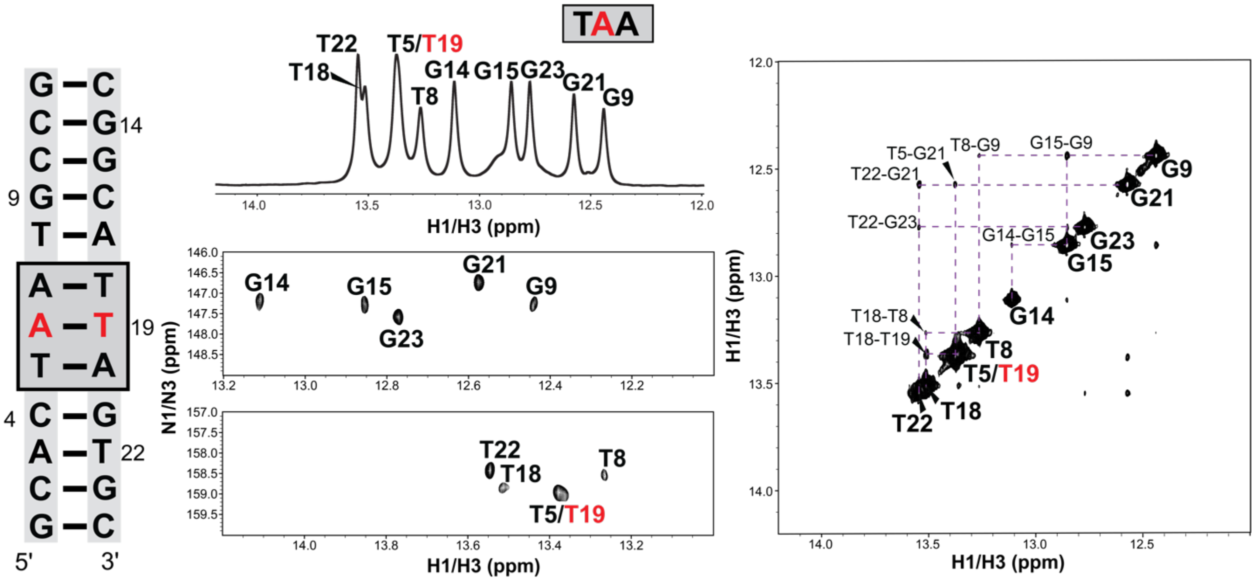
TAA
T22
T18
T5/T19
T8
G14
G15
G23
G21
G9
H1/H3 (ppm)
N1/N3 (ppm)
T5-G21
T8-G9
G15-G9
T22-G21
T22-G23
G14-G15
T18-T8
T18-T19
5'
3'
14
9
19
4
22

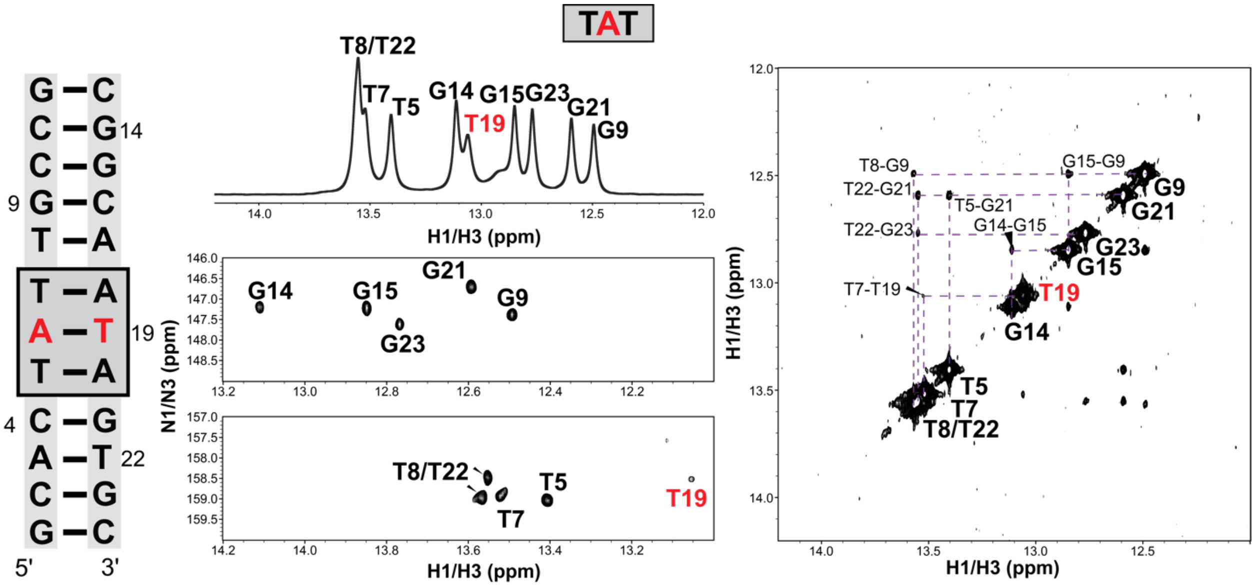
TAT
G C
C G 14
C G
9 G C
T A
T A
A T 19
T A
4 C G
A T 22
C G
G C
5'
3'
T8/T22
T7
T5
G14
T19
G15
G23
G21
G9
H1/H3 (ppm)
N1/N3 (ppm)
T8-G9
G15-G9
T22-G21
T5-G21
T22-G23
G14-G15
T7-T19

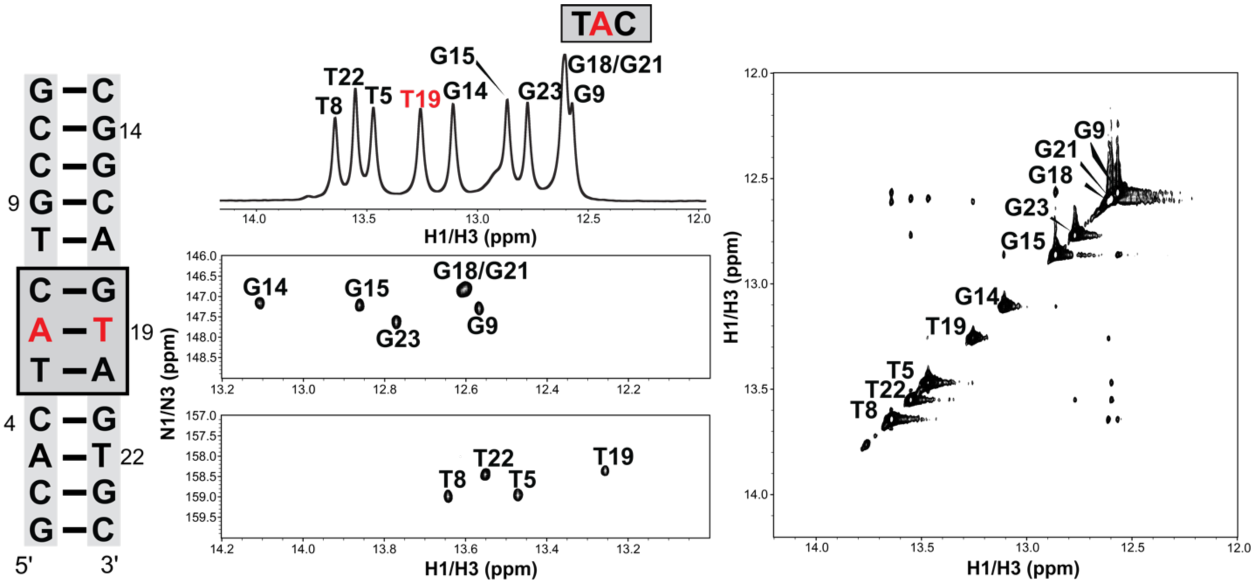
TAC
G C
C G 14
C G
9 G C
T A
C G
A T 19
T A
4 C G
A T 22
C G
G C
5'
3'
T22
T8
T5
T19
G15
G14
G23
G18/G21
G9
G21
G18
H1/H3 (ppm)
N1/N3 (ppm)

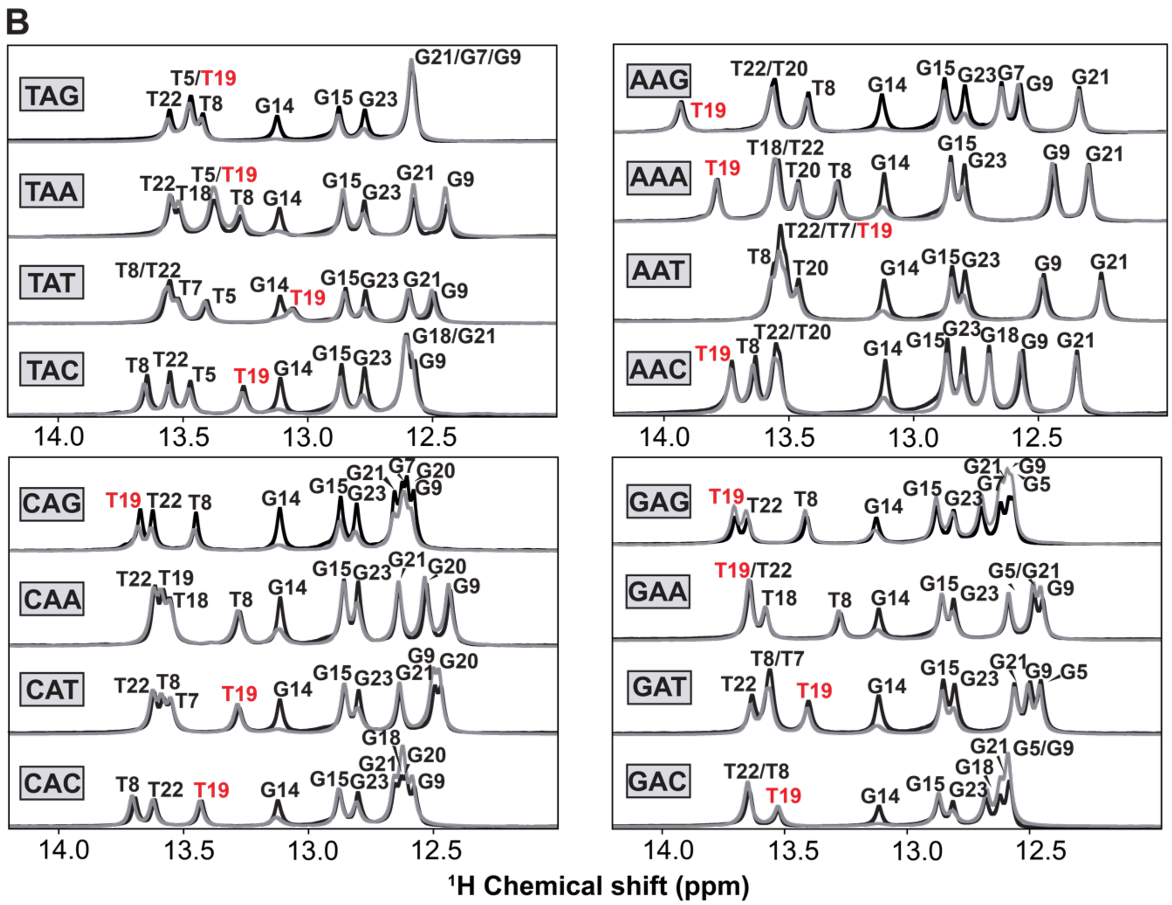


**Figure S1. Assignment of $^{1}$H Imino resonances.** (A) Imino $^{1}$H 1D, 2D$^{1}$H-$^{15}$N HSQC and $^{1}$H-$^{1}$H NOESY spectra used for assignment of imino resonances of the 16 duplexes used in this study. Also shown is the NOESY imino walk used for imino peak assignment. All spectra were collected at 700 MHz frequency at 25 mM NaCl, pH 6.8 and T=25ºC. Duplex constructs were used for all sequences except GAG and CAC, for which hairpin constructs were used. (B) Shown are the $^{1}$H 1D imino spectra of the 16 duplexes used in this study in 25 mM NaCl, pH 6.8 (black) and 100 mM NaCl, pH 8 (grey) at T=25ºC.

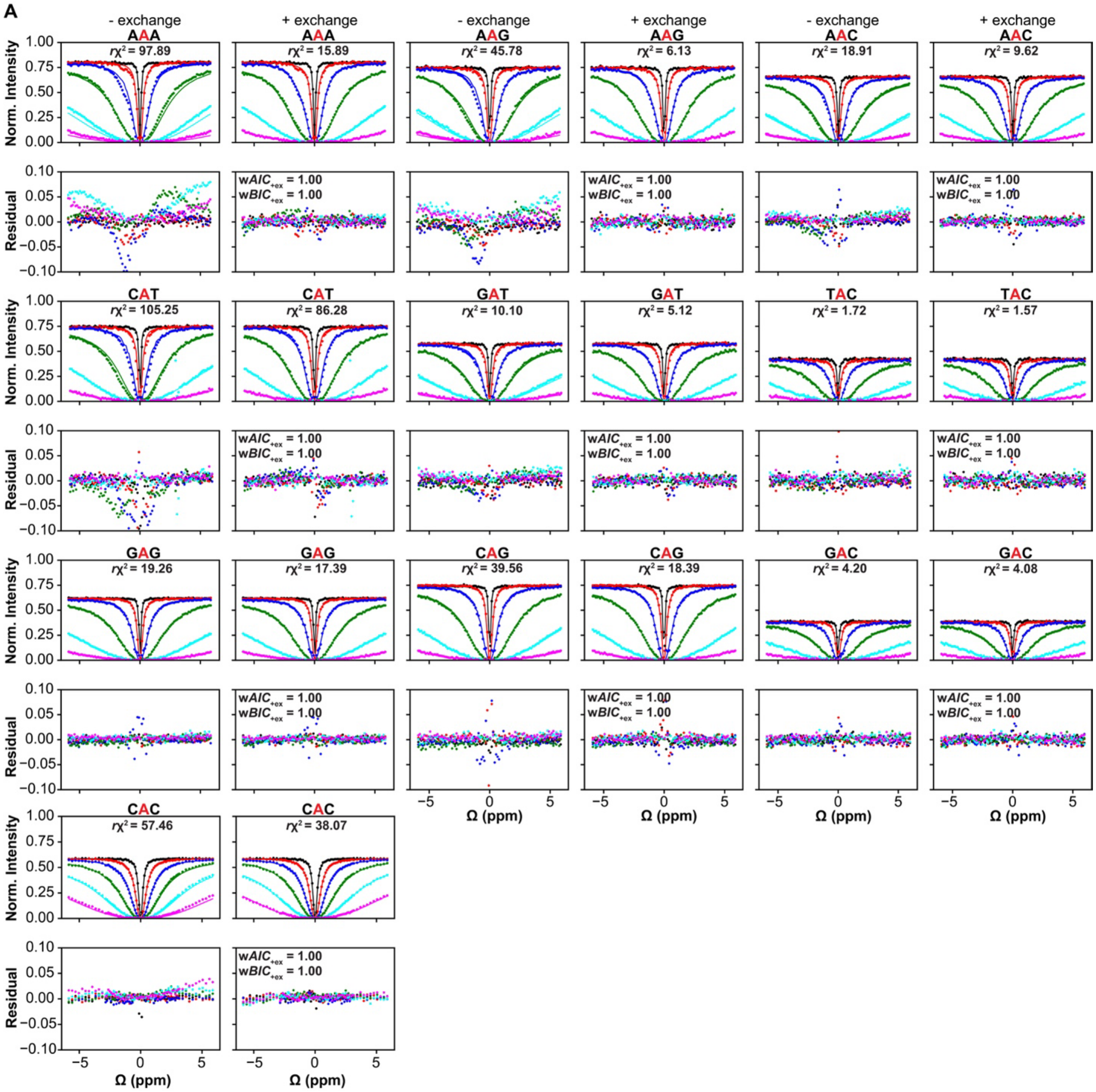
A
- exchange
+ exchange
AAA
AAG
AAC
CAT
GAT
TAC
GAG
CAG
GAC
CAC
rχ² = 97.89
rχ² = 15.89
rχ² = 45.78
rχ² = 6.13
rχ² = 18.91
rχ² = 9.62
rχ² = 105.25
rχ² = 86.28
rχ² = 10.10
rχ² = 5.12
rχ² = 1.72
rχ² = 1.57
rχ² = 19.26
rχ² = 17.39
rχ² = 39.56
rχ² = 18.39
rχ² = 4.20
rχ² = 4.08
rχ² = 57.46
rχ² = 38.07
wAIC+ex = 1.00
wBIC+ex = 1.00
Norm. Intensity
Residual
Ω (ppm)

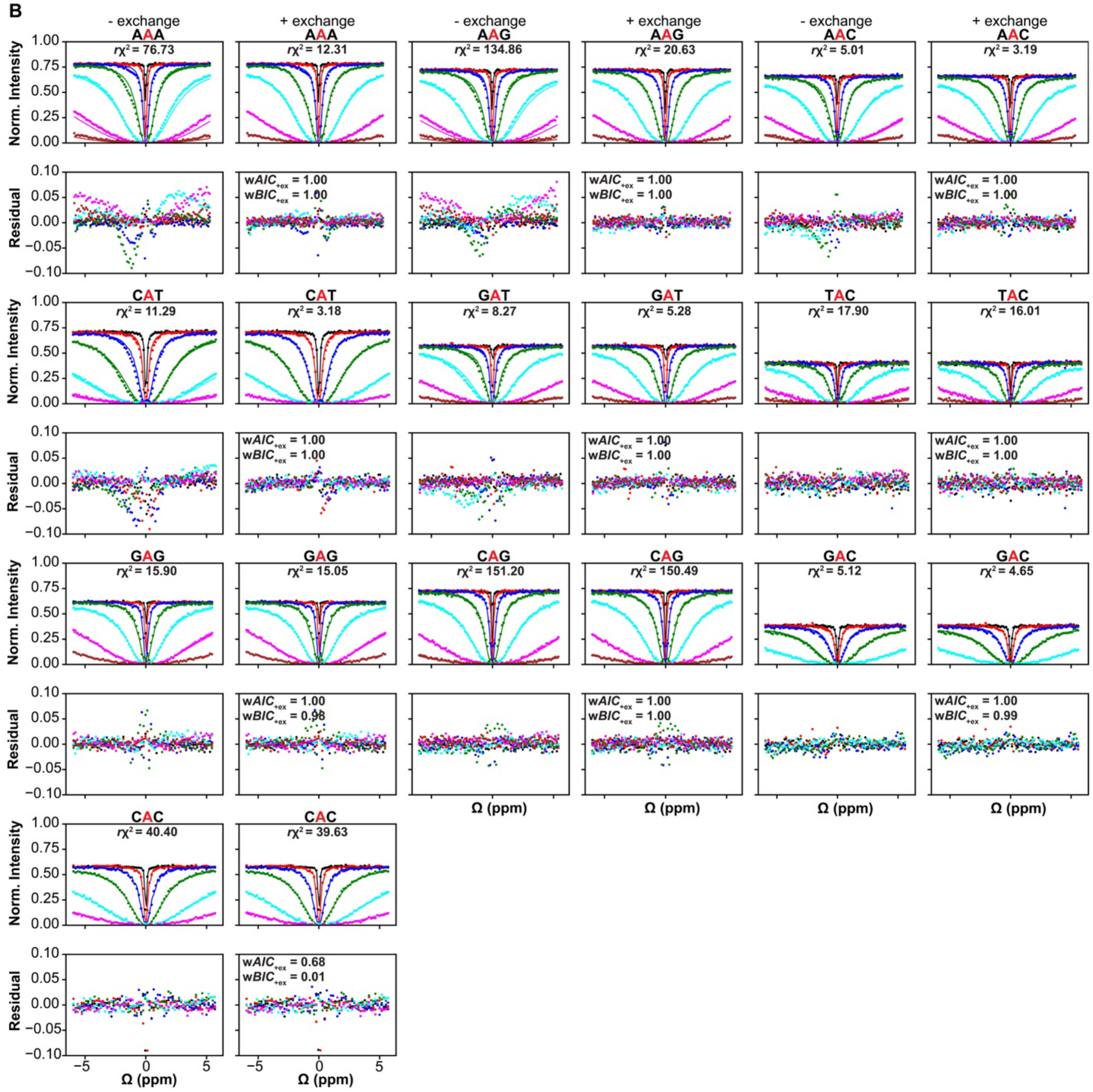


**Figure S2. [1]H CEST profiles for ten trinucleotide sequence contexts.** (A) Shown are the [1]H CEST profiles for ten trinucleotide sequence contexts measured at 25 mM NaCl, pH 6.8 and T=25ºC. The solid lines represent the fit of the data to the Bloch-McConell equation with and without exchange, respectively. The bottom panels show the corresponding residuals of the fit. Also shown are the reduced chi-squares ($r\chi^2$), the Akaike (w$AIC_{+ex}$) and Bayesian information weights (w$BIC_{+ex}$) for the fits with exchange (1,2). The radio frequency powers used in the [1]H

CEST experiments for AAA - GAC are color-coded 30 Hz (black), 90 Hz (red), 270 Hz (blue), 810 Hz (green), 2430 Hz (cyan) and 5000 Hz (pink) and for CAC are color-coded as 100 Hz (black), 250 Hz (red), 500 Hz (blue), 1000 Hz (green), 2000 Hz (cyan) and 4000 Hz (pink). Except for CAC, which was collected at 900 MHz, all other datasets were measured at 600 MHz. (B) Same as (A), except data were collected at 100 mM NaCl, pH 8 and T=25ºC at 600 MHz. The radio frequency powers used in the $^1$H CEST experiments for CAT, TAC and CAC are color-coded 30 Hz (black), 90 Hz (red), 270 Hz (blue), 810 Hz (green), 2430 Hz (cyan) and 5000 Hz (pink) and for all other sequence contexts are color-coded 10 Hz (black), 30 Hz (red), 90 Hz (blue), 270 Hz (green), 810 Hz (cyan), 2430 Hz (pink) and 5000 Hz (brown). The error bars for the $^1$H CEST data are obtained from the reference no RF irradiation experiment as described in Methods. The error bars are smaller than the data points.

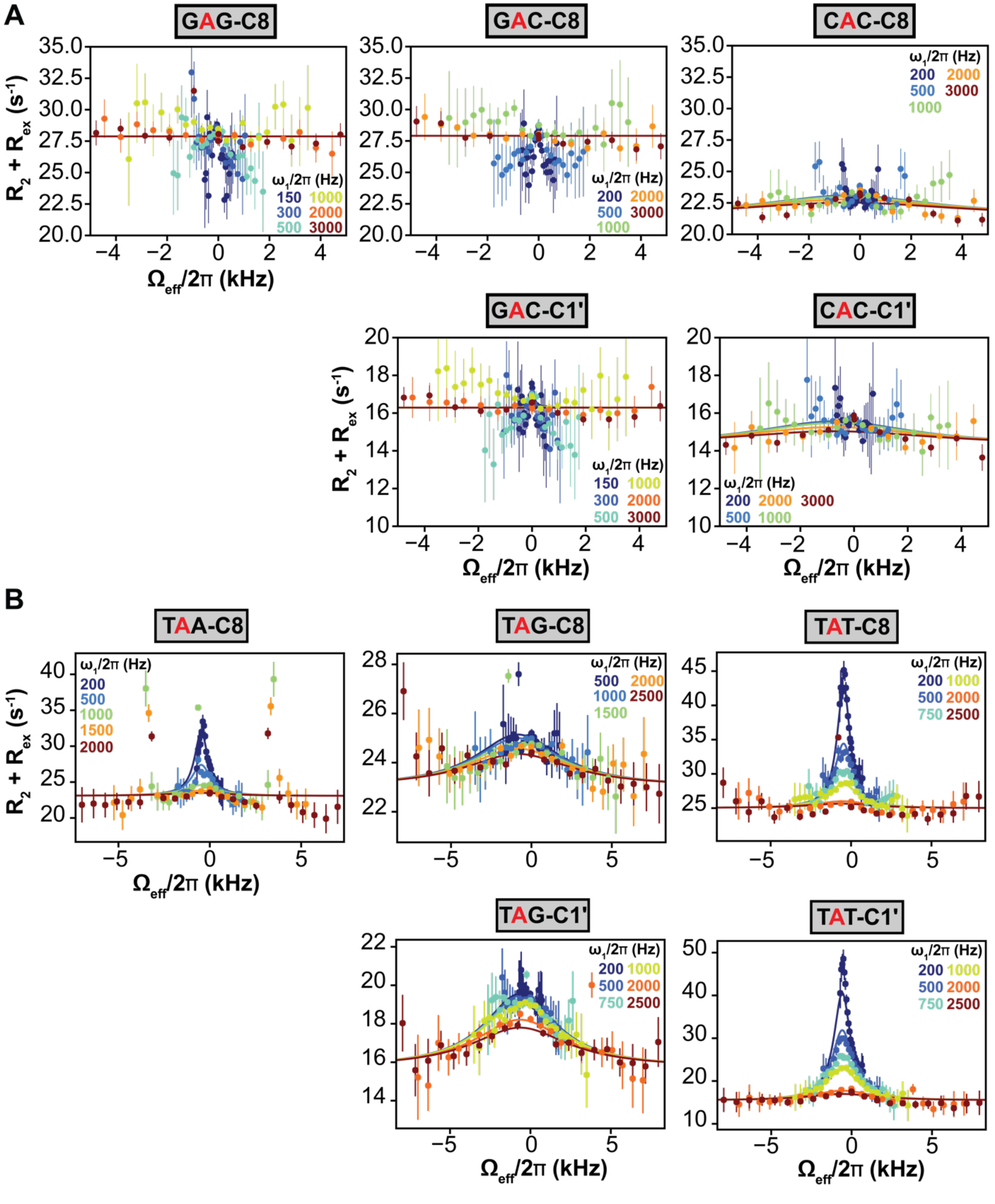


**Figure S3. Off-resonance $^{13}$C $R_{1\rho}$ profiles for G-C rich and TA step trinucleotides.** (A) Shown are the $^{13}$C $R_{1\rho}$ profiles for the central adenine in G-C rich triplets which showed fast exchange in

$^{1}H$ CEST. Measurements were performed at 800 MHz (GAG-C8, GAC-C8 and GAC-C1') or 700 MHz (CAC-C8 and CAC-C1'). (B) Shown are the off-resonance $^{13}C$ $R_{1\rho}$ profiles for the central adenine in TA step sequences where the T-H3 imino peak was overlapped with other residues. Measurements were performed at 600 MHz (TAA-C8) or 700 MHz (TAG-C8, TAG-C1', TAT-C8 and TAT-C1'). All measurements were performed at 25 mM NaCl, pH 6.8 and T=25ºC. The central adenine (highlighted in red) was isotopically labeled with $^{13}C$ and $^{15}N$. The error bars represent the experimental uncertainty in the $R_{1\rho}$ data estimated using a Monte-Carlo scheme as described previously (3).

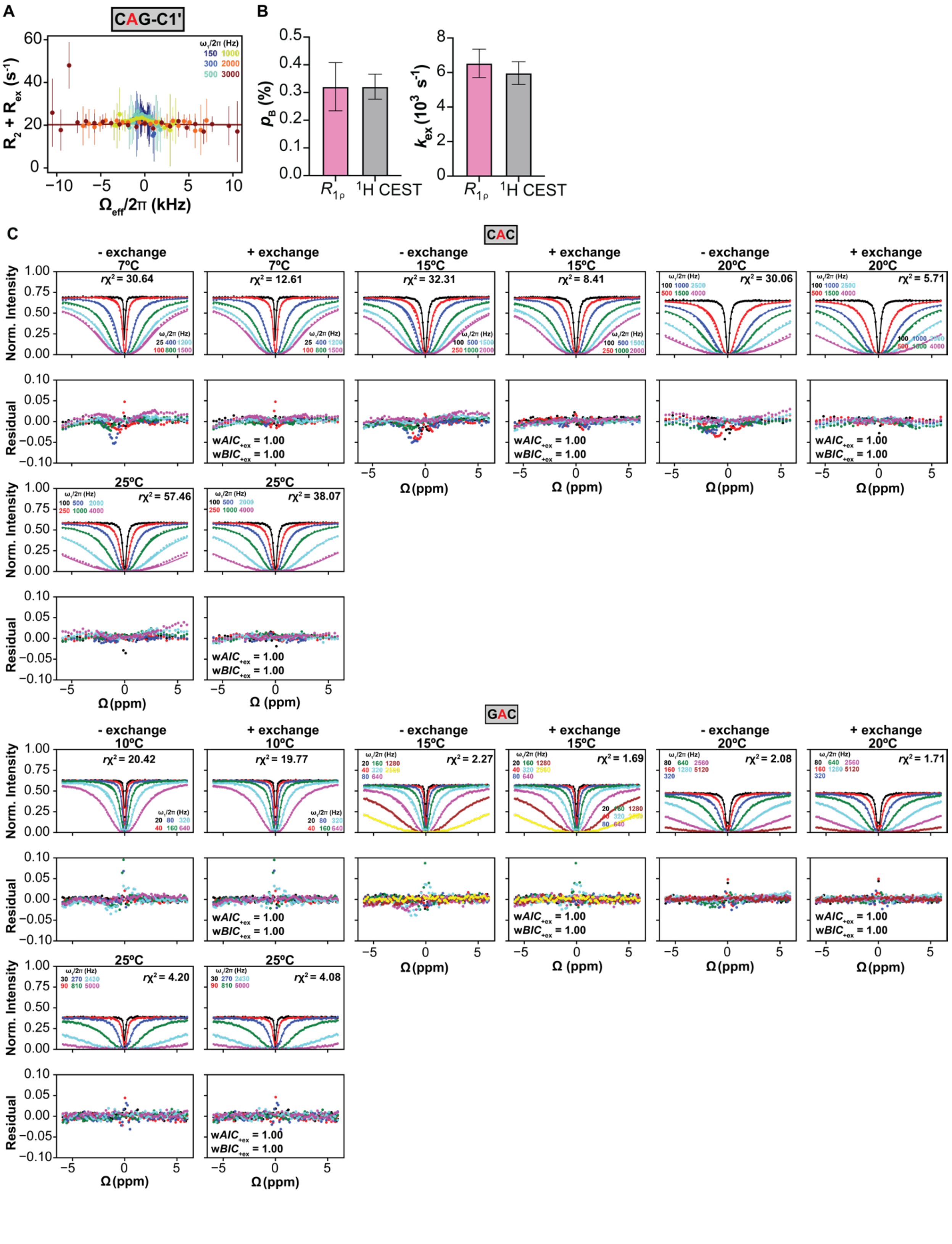
A
CAG-C1'
$R_2 + R_{ex}$ (s$^{-1}$)
$\Omega_{eff}/2\pi$ (kHz)
$\omega_1/2\pi$ (Hz)
150 1000
300 2000
500 3000
B
$p_B$ (%)
$k_{ex}$ ($10^3$ s$^{-1}$)
$R_{1\rho}$
$^1$H CEST
C
CAC
- exchange
+ exchange
7ºC
15ºC
20ºC
25ºC
$r\chi^2$ = 30.64
$r\chi^2$ = 12.61
$r\chi^2$ = 32.31
$r\chi^2$ = 8.41
$r\chi^2$ = 30.06
$r\chi^2$ = 5.71
$r\chi^2$ = 57.46
$r\chi^2$ = 38.07
Norm. Intensity
Residual
w$AIC_{+ex}$ = 1.00
w$BIC_{+ex}$ = 1.00
Ω (ppm)
GAC
10ºC
15ºC
20ºC
25ºC
$r\chi^2$ = 20.42
$r\chi^2$ = 19.77
$r\chi^2$ = 2.27
$r\chi^2$ = 1.69
$r\chi^2$ = 2.08
$r\chi^2$ = 1.71
$r\chi^2$ = 4.20
$r\chi^2$ = 4.08

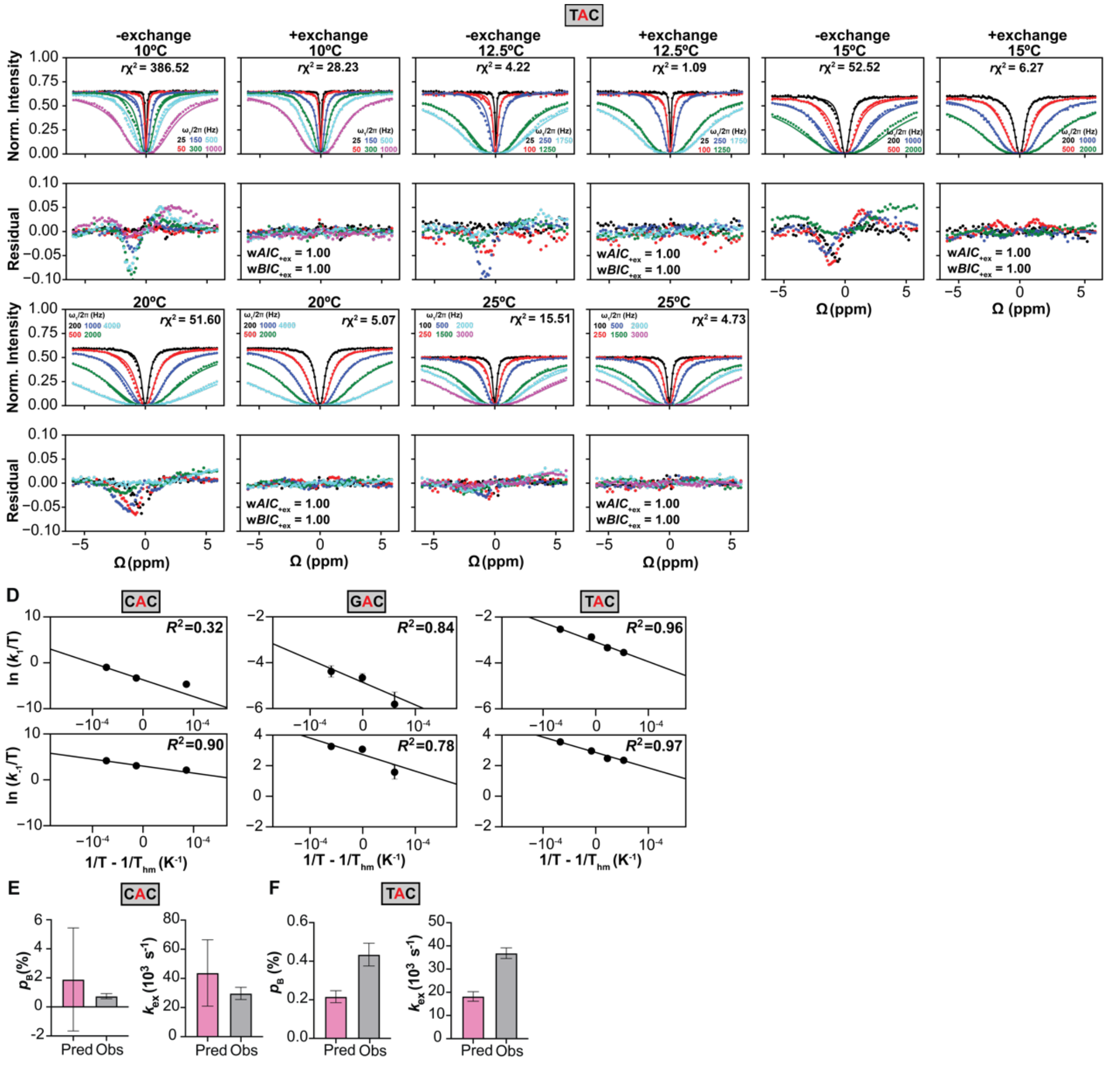

**Figure S4. Resolving fast exchange in GC-rich sequences.** (A) Off-resonance $^{13}C$ $R_{1\rho}$ C1' and C8 profiles for CAG measured at 700 MHz and at 25 mM NaCl, pH 6.8 and T=15ºC. The error bars represent the experimental uncertainty in the $R_{1\rho}$ data estimated using a Monte-Carlo scheme as described previously (3). (B) Comparison of Hoogsteen population ($p_B$) and exchange rate ($k_{ex}$) measured using $R_{1\rho}$ and $^1H$ CEST for CAG sequence context at T=15ºC. The error bars represent the uncertainty in the exchange parameters (Methods) calculated as described previously (3-6). (C) Temperature dependent $^1H$ CEST measurements of Hoogsteen dynamics for CAC at 900 MHz, GAC at 600 MHz, and TAC at 800 MHz. All measurements were performed at 25 mM NaCl, pH 6.8 and at indicated temperatures. The error bars for the $^1$H CEST data are obtained from the reference no RF irradiation experiment as described in Methods. The error bars are smaller than the data points. (D) Temperature dependence of forward ($k_1$) and backward ($k_{-1}$) rate constants for the Watson-Crick to Hoogsteen exchange for CAC, GAC and TAC sequence contexts. The $R^2$ denotes coefficient of determination. The errors bars are obtained by propagating the uncertainty in the exchange parameters for various temperatures. (E, F) Comparison of the Hoogsteen population ($p_B$) and exchange rate ($k_{ex}$) for CAC (E) and TAC (F) at T=25ºC obtained using temperature interpolation (Pred) and those deduced from $^1$H CEST measurements (Obs) performed at T=25ºC. The uncertainty in the predicted parameters (Pred) is obtained by propagating the errors in the temperature dependent rate constants $k_1$ and $k_{-1}$. The uncertainty in the observed parameters (Obs) represent the fitting error of the $^1$H CEST data calculated as described previously (4,5).

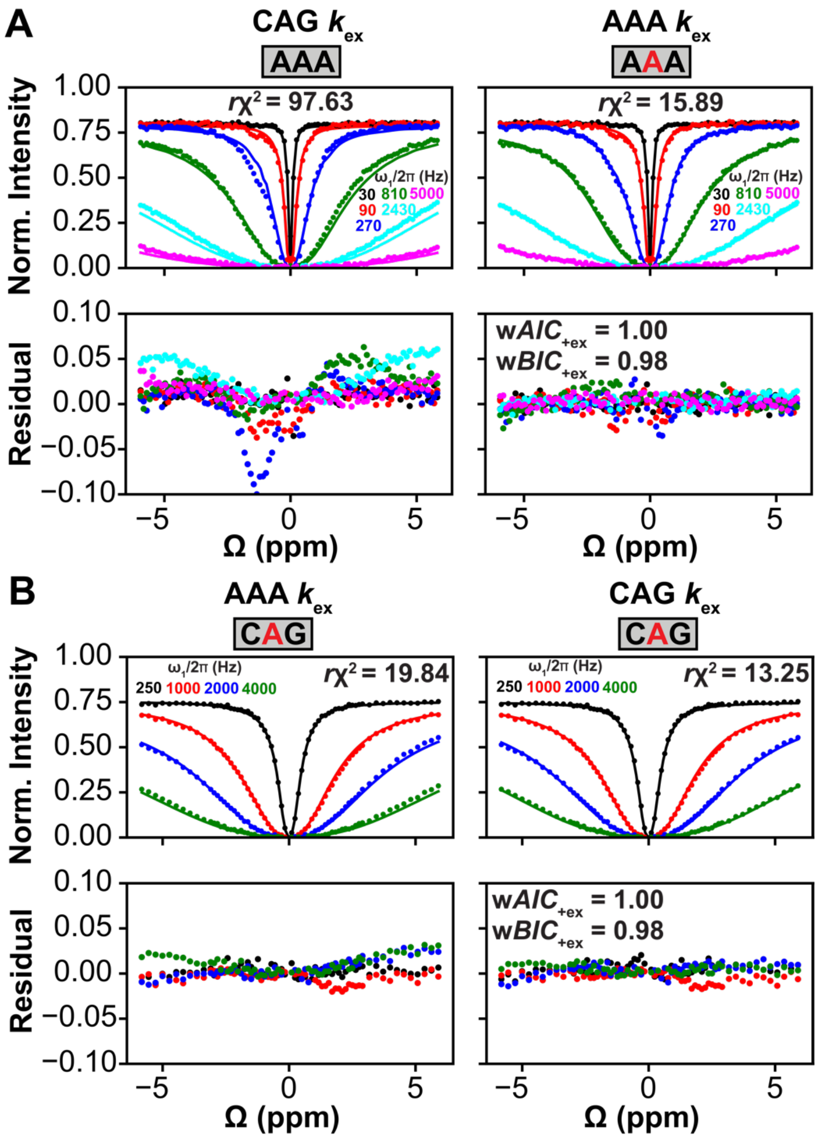


**Figure S5. Representative cross-fit analysis to verify Hoogsteen exchange parameters.** (A) Comparison of fit to the $^1$H CEST data measured for the AAA trinucleotide sequence context when fixing $k_{ex}$ = 39,000 s$^{-1}$ to the value obtained for CAG (left) or to its best-fitted value $k_{ex}$ ~4,000 s$^{-1}$ (right). (B) As in (A) but when fitting the $^1$H CEST data measured for CAG trinucleotide sequence and fixing $k_{ex}$ ~4,000 s$^{-1}$ to the value obtained for AAA trinucleotide sequence context (left) or to its best-fitted value $k_{ex}$ ~39,000 s$^{-1}$ (right). The radio frequency powers used in the $^1$H CEST experiments are color-coded. The error bars for the $^1$H CEST data are obtained from the reference no RF irradiation experiment as described in Methods. The error bars are smaller than the data points.

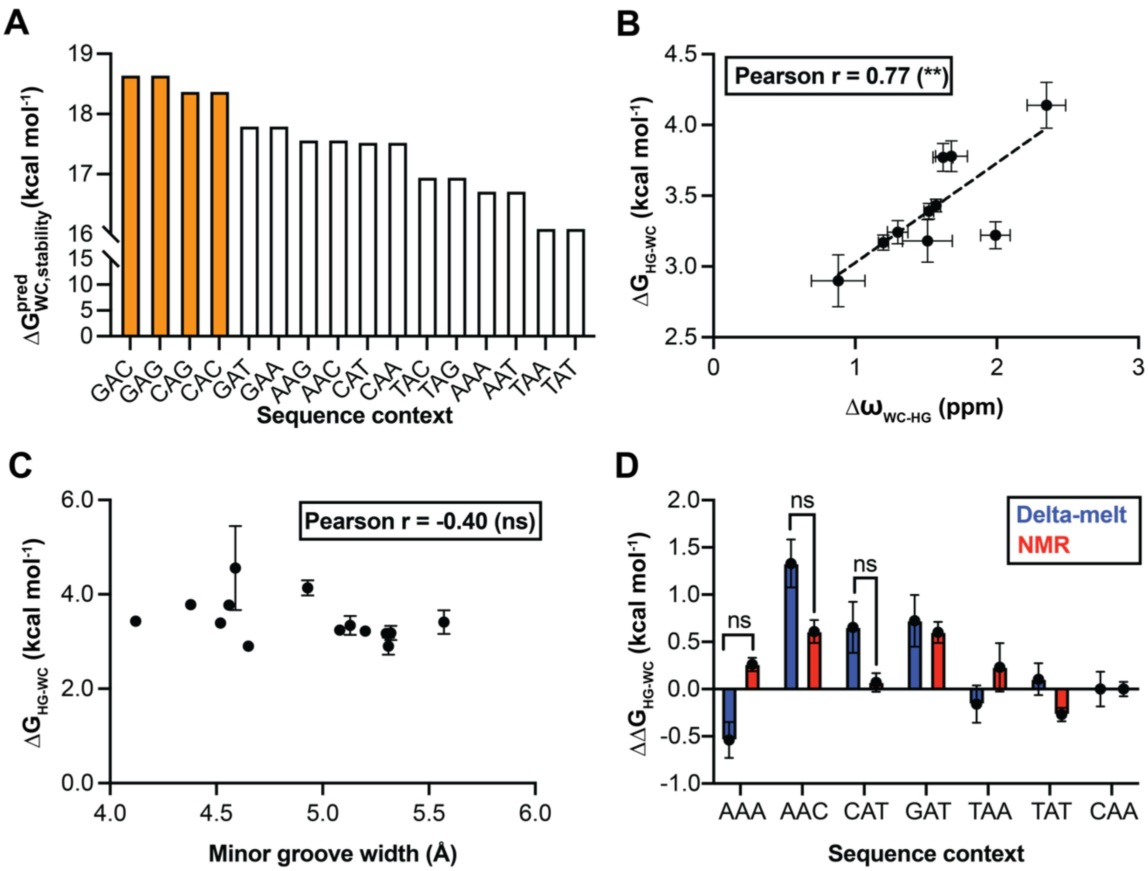


**Figure S6. Relationship between propensity to form Hoogsteen and biophysical properties.** (A) Overall stability of the duplexes predicted using the latest nearest-neighbor parameters (7). The most stable GC-rich sequence contexts exhibiting fast Hoogsteen exchange are colored in orange. (B) Correlation between T-H3 chemical shift difference between the Watson-Crick and Hoogsteen conformational states ($\Delta\omega_{WC\text{-}HG}$) and Hoogsteen propensities ($\Delta G_{WC\text{-}HG}$). Shown is the Pearson correlation coefficient (r) and its statistical significance assessed by the two-tailed p-value (0.0086). The error bar for $\Delta\omega_{WC\text{-}HG}$ represents the fitting error of the $^{1}$H CEST data calculated as described previously (4,5). The error bar for ($\Delta G_{WC\text{-}HG}$) is obtained by propagating the error in the exchange parameters. (C) Correlation between the minor groove width of the duplexes at the center of the trinucleotide calculated using DNAshape (8) and conformational propensity to form Hoogsteen ($\Delta G_{WC\text{-}HG}$) measured in this study using NMR. Shown is the Pearson correlation coefficient (r) and its statistical significance assessed by the two-tailed p-value (0.158). The error in $\Delta G_{WC\text{-}HG}$ is obtained by propagating the error in the exchange parameters. (D) The relative Hoogsteen conformational propensities for 7 overlapping sequences measured in this study using NMR and delta-melt in a prior study (9) calculated with respect to CAA sequence context. The statistical significance (ns) is assessed based on the (~ $\pm$ 0.83 kcal mol$^{-1}$) 95% confidence interval of the relative Hoogsteen conformational penalties predicted by delta-melt.

### Supplementary tables

**Table S1.** Parameters for Watson-Crick – Hoogsteen exchange at 25ºC obtained from $^{1}$H CEST measurements performed at 25 mM NaCl, pH 6.8 and T=25ºC

| Sequence | *p*B (%) | $k_{ex}$ ($10^3$ $s^{-1}$) | Δω (ppm) | $R_1$ ($s^{-1}$) | $R_2$ ($s^{-1}$) | $^{1}$H MHz |
|---|---|---|---|---|---|---|
| **AAA** | 0.31 ± 0.01 | 4.05 ± 0.19 | -1.57 ± 0.03 | 2.24 ± 0.01 | 15.05 ± 0.21 | 600 |
| **AAG** | 0.33 ± 0.02 | 6.54 ± 0.33 | -1.52 ± 0.03 | 2.91 ± 0.01 | 15.32 ± 0.18 | 600 |
| **AAC** | 0.17 ± 0.02 | 7.97 ± 0.77 | -1.68 ± 0.11 | 4.23 ± 0.01 | 18.12 ± 0.23 | 600 |
| **CAT** | 0.42 ± 0.04 | 6.80 ± 0.46 | -1.30 ± 0.07 | 2.93 ± 0.01 | 14.61 ± 0.14 | 600 |
| **GAT** | 0.17 ± 0.02 | 6.55 ± 0.64 | -1.62 ± 0.07 | 5.53 ± 0.01 | 17.97 ± 0.20 | 600 |
| **CAA** | 0.47 ± 0.03 | 5.12 ± 0.23 | -1.20 ± 0.04 | 2.42 ± 0.01 | 14.74 ± 0.16 | 600 |
| **TAC** | 0.43 ± 0.06 | 36.85 ± 2.31 | -1.99 ± 0.10 | 8.51 ± 0.01 | 13.46 ± 0.64 | 800 |
| **GAG** | 0.09 ± 0.02 | 23.17 ± 2.97 | -2.35 ± 0.14 | 4.65 ± 0.01 | 20.73 ± 0.62 | 900 |
| **CAG** | 0.47 ± 0.10 | 38.98 ± 4.10 | -1.51 ± 0.18 | 2.91 ± 0.02 | 14.08 ± 0.79 | 900 |
| **CAC** | 0.74 ± 0.18 | 29.70 ± 4.21 | -0.88 ± 0.19 | 5.35 ± 0.01 | 18.12 ± 0.61 | 900 |

**Table S2.** Parameters for Watson-Crick – Hoogsteen exchange at 25 mM NaCl, pH 6.8 and T=25ºC interpolated from temperature dependent $^{1}H$ CEST measurements.

| **Sequence** | ***p*B (%)** | **$k_{ex}$ ($10^3$ $s^{-1}$)** | **$^{1}H$ MHz** |
|---|---|---|---|
| **GAC** | 0.05 ± 0.04 | 16.16 ± 12.95 | 600 |
| **TAC** | 0.22 ± 0.03 | 18.22 ± 2.05 | 800 |
| **CAG** | 0.38 ± 0.07 | 31.85 ± 3.81 | 900 |
| **CAC** | 1.89 ± 3.54 | 43.74 ± 22.76 | 900 |

**Table S3.** Parameters for Watson-Crick – Hoogsteen exchange at 100 mM NaCl, pH 8 and T=25ºC obtained from $^{1}$H CEST measurements.

| Sequence | *p*B (%) | $k_{ex}$ ($10^3$ $s^{-1}$) | Δω (ppm) | $R_1$ ($s^{-1}$) | $R_2$ ($s^{-1}$) | $^{1}$H MHz |
|---|---|---|---|---|---|---|
| **AAA** | 0.29 ± 0.01 | 3.53 ± 0.16 | -1.53 ± 0.03 | 2.49 ± 0.01 | 18.47 ± 0.19 | 600 |
| **AAG** | 0.34 ± 0.02 | 5.03 ± 0.22 | -1.51 ± 0.03 | 3.30 ± 0.01 | 19.75 ± 0.24 | 600 |
| **AAC** | 0.14 ± 0.01 | 5.51 ± 0.72 | -1.77 ± 0.09 | 4.16 ± 0.01 | 19.50 ± 0.19 | 600 |
| **CAT** | 0.81 ± 0.09 | 7.50 ± 0.40 | -1.02 ± 0.06 | 3.44 ± 0.01 | 17.45 ± 0.17 | 600 |
| **GAT** | 0.16 ± 0.02 | 8.29 ± 0.82 | -2.03 ± 0.15 | 5.63 ± 0.01 | 20.20 ± 0.18 | 600 |
| **CAA** | 0.78 ± 0.05 | 6.41 ± 0.29 | -1.04 ± 0.04 | 2.84 ± 0.01 | 16.8 ± 0.27 | 600 |

**Table S4.** Parameters for Watson-Crick – Hoogsteen exchange at 25 mM NaCl, pH 6.8 and at indicated temperatures obtained from $^{13}$C $R_{1\rho}$ measurements performed at and field strengths.

| Sequence | Nucleus | *p*B (%) | $k_{ex}$ ($10^3$ $s^{-1}$) | Δω (ppm) | $R_1$ ($s^{-1}$) | $R_2$ ($s^{-1}$) | $^1$H MHz |
|---|---|---|---|---|---|---|---|
| **TAA, 25ºC** | C8 | 0.32 ± 0.07 | 2.49 ± 0.64 | 3.1 ± 0.5 | 2.18 ± 0.11 | 23.08 ± 0.34 | 600 |
| **AAG, 25ºC** | C8 | 0.32 ± 0.03 | 5.57 ± 0.44 | 2.68 ± 0.19 | 1.87 ± 0.08 | 27.50 ± 0.28 | 800 |
| | C1' | | | 3.71 ± 0.23 | 1.19 ± 0.06 | 15.78 ± 0.22 | 800 |
| **TAG, 25ºC** | C8 | 0.36 ± 0.11 | 17.94 ± 1.75 | 2.90 ± 0.40 | 1.67 ± 0.02 | 23.13 ± 0.22 | 700 |
| | C1' | | | 4.06 ± 0.55 | 1.53 ± 0.02 | 15.57 ± 0.35 | 700 |
| **TAT, 25ºC** | C8 | 0.74 ± 0.04 | 3.24 ± 0.15 | 2.58 ± 0.09 | 2.03 ± 0.06 | 24.63 ± 0.19 | 700 |
| | C1' | | | 3.36 ± 0.12 | 1.34 ± 0.08 | 16.0 ± 0.23 | 700 |
| **sl-TAG, 25ºC** | C8 | 0.23 ± 0.21 | 15.80 ± 5.50 | 1.0 ± 0.77 | 2.02 ± 0.04 | 21.88 ± 0.29 | 800 |
| | C1' | | | 3.98 ± 1.40 | 1.49 ± 0.06 | 13.22 ± 1.09 | 800 |
| **CAG, 15ºC** | C8 | 0.15 ± 0.02 | 5.98 ± 0.44 | 3.79 ± 0.24 | 1.94 ± 0.02 | 31.20 ± 0.09 | 700 |
| | C1' | | | 4.51 ± 0.40 | 1.79 ± 0.07 | 20.49 ± 0.29 | 700 |

**Table S5.** RF powers and offsets used in [1]H CEST experiments performed at 25 mM NaCl, pH 6.8 and T=25ºC with indicated relaxation durations at the indicated fields.

| Sample | [RF field power] [offset frequencies] |
|---|---|
| | [ω/2π (Hz)] [Ω/2π (Hz)] |
| AAA, 25ºC<br>$T_{EX}$ = 100 ms<br>600 MHz | [30], [-3487, -3415, -3342, -3269, -3196, -3124, -3051, -2978, -2906, -2833, -2760, -2688, -2615, -2542, -2470, -2397, -2324, -2252, -2179, -2106, -2034, -1961, -1888, -1815, -1743, -1670, -1597, -1525, -1452, -1379, -1307, -1234, -1161, -1089, -1016, -943, -871, -798, -725, -652, -580, -507, -434, -362, -289, -216, -144, -71, 2, 74, 147, 220, 292, 365, 438, 510, 583, 656, 729, 801, 874, 947, 1019, 1092, 1165, 1237, 1310, 1383, 1455, 1528, 1601, 1673, 1746, 1819, 1892, 1964, 2037, 2110, 2182, 2255, 2328, 2400, 2473, 2546, 2618, 2691, 2764, 2836, 2909, 2982, 3055, 3127, 3200, 3273, 3345, 3418, 3491, 3563]<br>[90], [-3487, -3415, -3342, -3269, -3196, -3124, -3051, -2978, -2906, -2833, -2760, -2688, -2615, -2542, -2470, -2397, -2324, -2252, -2179, -2106, -2033, -1961, -1888, -1815, -1743, -1670, -1597, -1525, -1452, -1379, -1307, -1234, -1161, -1089, -1016, -943, -871, -798, -725, -652, -580, -507, -434, -362, -289, -216, -144, -71, 2, 74, 147, 220, 292, 365, 438, 511, 583, 656, 729, 801, 874, 947, 1019, 1092, 1165, 1237, 1310, 1383, 1455, 1528, 1601, 1674, 1746, 1819, 1892, 1964, 2037, 2110, 2182, 2255, 2328, 2400, 2473, 2546, 2618, 2691, 2764, 2836, 2909, 2982, 3055, 3127, 3200, 3273, 3345, 3418, 3491, 3563]<br>[270], [-3487, -3414, -3342, -3269, -3196, -3124, -3051, -2978, -2906, -2833, -2760, -2688, -2615, -2542, -2470, -2397, -2324, -2251, -2179, -2106, -2033, -1961, -1888, -1815, -1743, -1670, -1597, -1525, -1452, -1379, -1307, -1234, -1161, -1088, -1016, -943, -870, -798, -725, -652, -580, -507, -434, -362, -289, -216, -144, 74, 220, 293, 365, 438, 511, 583, 656, 729, 801, 874, 947, 1019, 1092, 1165, 1237, 1310, 1383, 1456, 1528, 1601, 1674, 1746, 1819, 1892, 1964, 2037, 2110, 2182, 2255, 2328, 2400, 2473, 2546, 2619, 2691, 2764, 2837, 2909, 2982, 3055, 3127, 3200, 3273, 3345, 3418, 3491, 3563]<br>[810], [-3487, -3414, -3342, -3269, -3196, -3124, -3051, -2978, -2906, -2833, -2760, -2688, -2615, -2542, -2470, -2397, -2324, -2252, -2179, -2106, -2033, -1961, -1888, -1815, -1743, -1670, -1597, -1525, -1452, -1379, -1307, -1234, -1161, -1089, -1016, -943, -870, -798, -725, -652, -580, -507, -434, -362, -289, -216, -144, 2, 220, 293, 365, 438, 511, 583, 656, 729, 801, 874, 947, 1019, 1092, 1165, 1237, 1310, 1383, 1455, 1528, 1601, 1674, 1746, 1819, 1892, 1964, 2037, 2110, 2182, 2255, 2328, 2400, 2473, 2546, 2618, 2691, 2764, 2837, 2909, 2982, 3055, 3127, 3200, 3273, 3345, 3418, 3491, 3563]<br>[2430], [-3487, -3414, -3342, -3269, -3196, -3124, -3051, -2978, -2906, -2833, -2760, -2688, -2615, -2542, -2470, -2397, -2324, -2252, -2179, -2106, -2033, -1961, -1888, -1815, -1743, -1670, -1597, -1525, -1452, -1379, -1307, -1234, -1161, -1089, -1016, -943, -870, -798, -725, -652, -580, -507, -362, -289, -144, 365, 511, 583, 656, 729, 801, 874, 947, 1019, 1092, 1165, 1237, 1310, 1383, 1455, 1528, 1601, 1674, 1746, 1819, 1892, 1964, 2037, 2110, 2182, 2255, 2328, |

| | |
|---|---|
| | 2400, 2473, 2546, 2618, 2691, 2764, 2837, 2909, 2982, 3055, 3127, 3200, 3273, 3345, 3418, 3491, 3563]<br>[5000], [-3487, -3415, -3342, -3269, -3196, -3124, -3051, -2978, -2906, -2833, -2760, -2688, -2615, -2542, -2470, -2397, -2324, -2252, -2179, -2106, -2033, -1961, -1888, -1815, -1743, -1670, -1597, -1525, -1452, -1379, -1307, -1234, -1161, -1089, -1016, -943, -871, -798, -725, -652, -580, -507, -362, -289, -144, -71, 74, 147, 220, 292, 365, 511, 583, 656, 874, 947, 1019, 1092, 1165, 1237, 1310, 1383, 1455, 1528, 1601, 1673, 1746, 1819, 1892, 1964, 2037, 2110, 2182, 2255, 2328, 2400, 2473, 2546, 2618, 2691, 2764, 2836, 2909, 2982, 3055, 3127, 3200, 3273, 3345, 3418, 3491, 3563] |
| AAG, 25ºC<br>$T_{EX}$ = 100 ms<br>600 MHz | [30], [-3461, -3388, -3315, -3243, -3170, -3097, -3025, -2952, -2879, -2807, -2734, -2661, -2589, -2516, -2443, -2370, -2298, -2225, -2152, -2080, -2007, -1934, -1862, -1789, -1716, -1644, -1571, -1498, -1426, -1353, -1280, -1207, -1135, -1062, -989, -917, -844, -771, -699, -626, -553, -481, -408, -335, -263, -190, -117, -44, 28, 101, 174, 246, 319, 392, 464, 537, 610, 682, 755, 828, 900, 973, 1046, 1118, 1191, 1264, 1337, 1409, 1482, 1555, 1627, 1700, 1773, 1845, 1918, 1991, 2063, 2136, 2209, 2281, 2354, 2427, 2500, 2572, 2645, 2718, 2790, 2863, 2936, 3008, 3081, 3154, 3226, 3299, 3372, 3444, 3517, 3590]<br>[90], [-3461, -3388, -3315, -3243, -3170, -3097, -3025, -2952, -2879, -2807, -2734, -2661, -2588, -2516, -2443, -2370, -2298, -2225, -2152, -2080, -2007, -1934, -1862, -1789, -1716, -1644, -1571, -1498, -1425, -1353, -1280, -1207, -1135, -1062, -989, -917, -844, -771, -699, -626, -553, -481, -408, -335, -263, -190, -117, -44, 101, 174, 246, 319, 392, 464, 537, 610, 682, 755, 828, 900, 973, 1046, 1119, 1191, 1264, 1337, 1409, 1482, 1555, 1627, 1700, 1773, 1845, 1918, 1991, 2063, 2136, 2209, 2282, 2354, 2427, 2500, 2572, 2645, 2718, 2790, 2863, 2936, 3008, 3081, 3154, 3226, 3299, 3372, 3444, 3517, 3590]<br>[270], [-3461, -3388, -3315, -3243, -3170, -3097, -3025, -2952, -2879, -2807, -2734, -2661, -2588, -2516, -2443, -2370, -2298, -2225, -2152, -2080, -2007, -1934, -1862, -1789, -1716, -1644, -1571, -1498, -1425, -1353, -1280, -1207, -1135, -1062, -989, -917, -844, -771, -699, -626, -553, -481, -408, -335, -262, -190, 101, 174, 246, 319, 392, 464, 537, 610, 682, 755, 828, 900, 973, 1046, 1119, 1191, 1264, 1337, 1409, 1482, 1555, 1627, 1700, 1773, 1845, 1918, 1991, 2063, 2136, 2209, 2282, 2354, 2427, 2500, 2572, 2645, 2718, 2790, 2863, 2936, 3008, 3081, 3154, 3226, 3299, 3372, 3444, 3517, 3590]<br>[810], [-3461, -3388, -3315, -3243, -3170, -3097, -3025, -2952, -2879, -2807, -2734, -2661, -2588, -2516, -2443, -2370, -2298, -2225, -2152, -2080, -2007, -1934, -1862, -1789, -1716, -1644, -1571, -1498, -1425, -1353, -1280, -1207, -1135, -1062, -989, -917, -844, -771, -699, -626, -553, -481, -408, -335, -190, -44, 101, 246, 319, 392, 464, 537, 610, 682, 755, 828, 900, 973, 1046, 1119, 1191, 1264, 1337, 1409, 1482, 1555, 1627, 1700, 1773, 1845, 1918, 1991, 2063, 2136, 2209, 2281, 2354, 2427, 2500, 2572, 2645, 2718, 2790, 2863, 2936, 3008, 3081, 3154, 3226, 3299, 3372, 3444, 3517, 3590]<br>[2430], [-3461, -3388, -3315, -3243, -3170, -3097, -3025, -2952, -2879, -2806, -2734, -2661, -2588, -2516, -2443, -2370, -2298, -2225, -2152, -2080, -2007, - |

| | |
|---|---|
| | 1934, -1862, -1789, -1716, -1644, -1571, -1498, -1425, -1353, -1280, -1207, -1135, -1062, -989, -917, -844, -771, -699, -626, -553, -335, -262, -190, -117, -44, 28, 392, 464, 610, 682, 755, 828, 901, 973, 1046, 1119, 1191, 1264, 1337, 1409, 1482, 1555, 1627, 1700, 1773, 1845, 1918, 1991, 2063, 2136, 2209, 2282, 2354, 2427, 2500, 2572, 2645, 2718, 2790, 2863, 2936, 3008, 3081, 3154, 3226, 3299, 3372, 3445, 3517, 3590]<br>[5000], [-3461, -3388, -3315, -3243, -3170, -3097, -3025, -2952, -2879, -2807, -2734, -2661, -2588, -2516, -2443, -2370, -2298, -2225, -2152, -2080, -2007, -1934, -1862, -1789, -1716, -1644, -1571, -1498, -1425, -1353, -1280, -1207, -1135, -1062, -989, -917, -844, -771, -699, -553, -481, -408, -335, -117, 28, 174, 464, 537, 610, 682, 828, 900, 973, 1046, 1119, 1191, 1264, 1337, 1409, 1482, 1555, 1627, 1700, 1773, 1845, 1918, 1991, 2063, 2136, 2209, 2282, 2354, 2427, 2500, 2572, 2645, 2718, 2790, 2863, 2936, 3008, 3081, 3154, 3226, 3299, 3372, 3444, 3517, 3590] |
| AAC, 25ºC<br>$T_{EX}$ = 100 ms<br>600 MHz | [30], [-3535, -3462, -3390, -3317, -3244, -3172, -3099, -3026, -2954, -2881, -2808, -2736, -2663, -2590, -2517, -2445, -2372, -2299, -2227, -2154, -2081, -2009, -1936, -1863, -1791, -1718, -1645, -1573, -1500, -1427, -1355, -1282, -1209, -1136, -1064, -991, -918, -846, -773, -700, -628, -555, -482, -410, -337, -264, -192, -119, -46, 27, 99, 172, 245, 317, 390, 463, 535, 608, 681, 753, 826, 899, 971, 1044, 1117, 1190, 1262, 1335, 1408, 1480, 1553, 1626, 1698, 1771, 1844, 1916, 1989, 2062, 2134, 2207, 2280, 2352, 2425, 2498, 2571, 2643, 2716, 2789, 2861, 2934, 3007, 3079, 3152, 3225, 3297, 3370, 3443, 3515, 3588]<br>[90], [-3535, -3463, -3390, -3317, -3245, -3172, -3099, -3026, -2954, -2881, -2808, -2736, -2663, -2590, -2518, -2445, -2372, -2300, -2227, -2154, -2082, -2009, -1936, -1863, -1791, -1718, -1645, -1573, -1500, -1427, -1355, -1282, -1209, -1137, -1064, -991, -919, -846, -773, -701, -628, -555, -482, -410, -337, -264, -192, -119, -46, 99, 172, 244, 317, 390, 462, 535, 608, 681, 753, 826, 899, 971, 1044, 1117, 1189, 1262, 1335, 1407, 1480, 1553, 1625, 1698, 1771, 1844, 1916, 1989, 2062, 2134, 2207, 2280, 2352, 2425, 2498, 2570, 2643, 2716, 2788, 2861, 2934, 3006, 3079, 3152, 3225, 3297, 3370, 3443, 3515, 3588]<br>[270], [-3535, -3463, -3390, -3317, -3244, -3172, -3099, -3026, -2954, -2881, -2808, -2736, -2663, -2590, -2518, -2445, -2372, -2300, -2227, -2154, -2081, -2009, -1936, -1863, -1791, -1718, -1645, -1573, -1500, -1427, -1355, -1282, -1209, -1137, -1064, -991, -919, -846, -773, -700, -628, -555, -482, -410, -337, -264, -192, -119, -46, 26, 172, 244, 317, 390, 463, 535, 608, 681, 753, 826, 899, 971, 1044, 1117, 1189, 1262, 1335, 1407, 1480, 1553, 1626, 1698, 1771, 1844, 1916, 1989, 2062, 2134, 2207, 2280, 2352, 2425, 2498, 2570, 2643, 2716, 2788, 2861, 2934, 3007, 3079, 3152, 3225, 3297, 3370, 3443, 3515, 3588]<br>[810], [-3535, -3463, -3390, -3317, -3244, -3172, -3099, -3026, -2954, -2881, -2808, -2736, -2663, -2590, -2518, -2445, -2372, -2300, -2227, -2154, -2082, -2009, -1936, -1863, -1791, -1718, -1645, -1573, -1500, -1427, -1355, -1282, -1209, -1137, -1064, -991, -919, -846, -773, -700, -628, -555, -482, -410, -337, -264, -119, 99, 172, 244, 317, 390, 463, 535, 608, 681, 753, 826, 899, 971, 1044, |

| | |
|---|---|
| | 1117, 1189, 1262, 1335, 1407, 1480, 1553, 1625, 1698, 1771, 1844, 1916, 1989, 2062, 2134, 2207, 2280, 2352, 2425, 2498, 2570, 2643, 2716, 2788, 2861, 2934, 3007, 3079, 3152, 3225, 3297, 3370, 3443, 3515, 3588]<br>[2430], [-3535, -3463, -3390, -3317, -3245, -3172, -3099, -3027, -2954, -2881, -2808, -2736, -2663, -2590, -2518, -2445, -2372, -2300, -2227, -2154, -2082, -2009, -1936, -1864, -1791, -1718, -1645, -1573, -1500, -1427, -1355, -1282, -1209, -1137, -1064, -991, -919, -846, -773, -701, -628, -555, -483, -410, -337, -192, -46, 99, 244, 317, 390, 462, 535, 608, 680, 753, 826, 899, 971, 1044, 1117, 1189, 1262, 1335, 1407, 1480, 1553, 1625, 1698, 1771, 1843, 1916, 1989, 2061, 2134, 2207, 2280, 2352, 2425, 2498, 2570, 2643, 2716, 2788, 2861, 2934, 3006, 3079, 3152, 3224, 3297, 3370, 3443, 3515, 3588]<br>[5000], [-3535, -3462, -3390, -3317, -3244, -3172, -3099, -3026, -2954, -2881, -2808, -2736, -2663, -2590, -2518, -2445, -2372, -2299, -2227, -2154, -2081, -2009, -1936, -1863, -1791, -1718, -1645, -1573, -1500, -1427, -1355, -1282, -1209, -1137, -1064, -991, -918, -846, -773, -700, -628, -555, -264, -119, 317, 390, 535, 753, 826, 899, 971, 1044, 1117, 1189, 1262, 1335, 1407, 1480, 1553, 1626, 1698, 1771, 1844, 1916, 1989, 2062, 2134, 2207, 2280, 2352, 2425, 2498, 2570, 2643, 2716, 2789, 2861, 2934, 3007, 3079, 3152, 3225, 3297, 3370, 3443, 3515, 3588] |
| CAT, 25ºC<br>$T_{EX}$ = 100 ms<br>600 MHz | [30], [-3584, -3512, -3439, -3366, -3294, -3221, -3148, -3076, -3003, -2930, -2858, -2785, -2712, -2639, -2567, -2494, -2421, -2349, -2276, -2203, -2131, -2058, -1985, -1913, -1840, -1767, -1695, -1622, -1549, -1476, -1404, -1331, -1258, -1186, -1113, -1040, -968, -895, -822, -750, -677, -604, -532, -459, -386, -314, -241, -168, -95, -23, 50, 123, 195, 268, 341, 413, 486, 559, 631, 704, 777, 849, 922, 995, 1068, 1140, 1213, 1286, 1358, 1431, 1504, 1576, 1649, 1722, 1794, 1867, 1940, 2012, 2085, 2158, 2231, 2303, 2376, 2449, 2521, 2594, 2667, 2739, 2812, 2885, 2957, 3030, 3103, 3175, 3248, 3321, 3393, 3466]<br>[90], [-3584, -3512, -3439, -3366, -3294, -3221, -3148, -3076, -3003, -2930, -2858, -2785, -2712, -2639, -2567, -2494, -2421, -2349, -2276, -2203, -2131, -2058, -1985, -1913, -1840, -1767, -1695, -1622, -1549, -1476, -1404, -1331, -1258, -1186, -1113, -1040, -968, -895, -822, -750, -677, -604, -532, -459, -386, -314, -241, -168, -95, -23, 50, 123, 195, 268, 341, 413, 486, 559, 631, 704, 777, 849, 922, 995, 1068, 1140, 1213, 1286, 1358, 1431, 1504, 1576, 1649, 1722, 1794, 1867, 1940, 2012, 2085, 2158, 2230, 2303, 2376, 2449, 2521, 2594, 2667, 2739, 2812, 2885, 2957, 3030, 3103, 3175, 3248, 3321, 3393, 3466]<br>[270], [-3584, -3512, -3439, -3366, -3294, -3221, -3148, -3075, -3003, -2930, -2857, -2785, -2712, -2639, -2567, -2494, -2421, -2349, -2276, -2203, -2131, -2058, -1985, -1913, -1840, -1767, -1694, -1622, -1549, -1476, -1404, -1331, -1258, -1186, -1113, -1040, -968, -895, -822, -750, -677, -604, -531, -459, -386, -313, -241, -168, -95, 50, 123, 195, 268, 341, 413, 486, 559, 632, 704, 777, 850, 922, 995, 1068, 1140, 1213, 1286, 1358, 1431, 1504, 1576, 1649, 1722, 1794, 1867, 1940, 2013, 2085, 2158, 2231, 2303, 2376, 2449, 2521, 2594, 2667, 2739, 2812, 2885, 2957, 3030, 3103, 3176, 3248, 3321, 3394, 3466] |

| | |
|---|---|
| | [810], [-3584, -3512, -3439, -3366, -3294, -3221, -3148, -3075, -3003, -2930, -2857, -2785, -2712, -2639, -2567, -2494, -2421, -2349, -2276, -2203, -2131, -2058, -1985, -1913, -1840, -1767, -1694, -1622, -1549, -1476, -1404, -1331, -1258, -1186, -1113, -1040, -968, -895, -822, -750, -677, -604, -531, -459, -386, -313, -241, -168, -95, 268, 341, 413, 486, 559, 631, 704, 777, 850, 922, 995, 1068, 1140, 1213, 1286, 1358, 1431, 1504, 1576, 1649, 1722, 1794, 1867, 1940, 2013, 2085, 2158, 2231, 2303, 2376, 2449, 2521, 2594, 2667, 2739, 2812, 2885, 2957, 3030, 3103, 3176, 3248, 3321, 3394, 3466]<br>[2430], [-3584, -3512, -3439, -3366, -3294, -3221, -3148, -3076, -3003, -2930, -2857, -2785, -2712, -2639, -2567, -2494, -2421, -2349, -2276, -2203, -2131, -2058, -1985, -1913, -1622, -1549, -1476, -1404, -1331, -1258, -1186, -1113, -1040, -968, -895, -822, -750, -677, -604, -531, -386, -313, -168, 50, 123, 195, 413, 486, 559, 631, 704, 777, 850, 922, 995, 1068, 1140, 1213, 1286, 1358, 1431, 1504, 1576, 1649, 1722, 1794, 1867, 1940, 2013, 2085, 2158, 2231, 2303, 2376, 2449, 2521, 2594, 2667, 2739, 2812, 2885, 2957, 3030, 3103, 3175, 3248, 3321, 3394, 3466]<br>[5000], [-3584, -3512, -3439, -3366, -3294, -3221, -3148, -3076, -3003, -2930, -2857, -2785, -2712, -2639, -2567, -2494, -2421, -2349, -2276, -2203, -2131, -2058, -1985, -1913, -1840, -1767, -1695, -1622, -1549, -1476, -1404, -1331, -1258, -1186, -1113, -1040, -968, -895, -822, -750, -677, -604, -532, -241, -168, -23, 195, 268, 486, 704, 849, 922, 995, 1068, 1140, 1213, 1286, 1358, 1431, 1504, 1576, 1649, 1722, 1794, 1867, 1940, 2012, 2085, 2158, 2231, 2303, 2376, 2449, 2521, 2594, 2667, 2739, 2812, 2885, 2957, 3030, 3103, 3175, 3248, 3321, 3394, 3466] |
| GAT, 25ºC<br>$T_{EX}$ = 100 ms<br>600 MHz | [30], [-3542, -3469, -3396, -3324, -3251, -3178, -3106, -3033, -2960, -2888, -2815, -2742, -2670, -2597, -2524, -2452, -2379, -2306, -2233, -2161, -2088, -2015, -1943, -1870, -1797, -1725, -1652, -1579, -1507, -1434, -1361, -1289, -1216, -1143, -1070, -998, -925, -852, -780, -707, -634, -562, -489, -416, -344, -271, -198, -126, -53, 92, 165, 238, 311, 383, 456, 529, 601, 674, 747, 819, 892, 965, 1037, 1110, 1183, 1255, 1328, 1401, 1474, 1546, 1619, 1692, 1764, 1837, 1910, 1982, 2055, 2128, 2200, 2273, 2346, 2418, 2491, 2564, 2636, 2709, 2782, 2855, 2927, 3000, 3073, 3145, 3218, 3291, 3363, 3436, 3509]<br>[90], [-3542, -3469, -3396, -3324, -3251, -3178, -3106, -3033, -2960, -2888, -2815, -2742, -2670, -2597, -2524, -2452, -2379, -2306, -2233, -2161, -2088, -2015, -1943, -1870, -1797, -1725, -1652, -1579, -1507, -1434, -1361, -1289, -1216, -1143, -1071, -998, -925, -852, -780, -707, -634, -562, -489, -416, -344, -271, -198, -126, 20, 92, 165, 238, 311, 383, 456, 529, 601, 674, 747, 819, 892, 965, 1037, 1110, 1183, 1255, 1328, 1401, 1474, 1546, 1619, 1692, 1764, 1837, 1910, 1982, 2055, 2128, 2200, 2273, 2346, 2418, 2491, 2564, 2636, 2709, 2782, 2855, 2927, 3000, 3073, 3145, 3218, 3291, 3363, 3436, 3509]<br>[270], [-3542, -3469, -3396, -3324, -3251, -3178, -3106, -3033, -2960, -2888, -2815, -2742, -2670, -2597, -2524, -2452, -2379, -2306, -2233, -2161, -2088, -2015, -1943, -1870, -1797, -1725, -1652, -1579, -1507, -1434, -1361, -1289, -1216, -1143, -1070, -998, -925, -852, -780, -707, -634, -562, -489, -416, -344, - |

| | |
|---|---|
| | 271, -198, -126, 20, 92, 165, 238, 311, 383, 456, 529, 601, 674, 747, 819, 892, 965, 1037, 1110, 1183, 1255, 1328, 1401, 1474, 1546, 1619, 1692, 1764, 1837, 1910, 1982, 2055, 2128, 2200, 2273, 2346, 2418, 2491, 2564, 2637, 2709, 2782, 2855, 2927, 3000, 3073, 3145, 3218, 3291, 3363, 3436, 3509]<br>[810], [-3542, -3469, -3396, -3324, -3251, -3178, -3106, -3033, -2960, -2888, -2815, -2742, -2670, -2597, -2524, -2452, -2379, -2306, -2233, -2161, -2088, -2015, -1943, -1870, -1797, -1725, -1652, -1579, -1507, -1434, -1361, -1289, -1216, -1143, -1071, -998, -925, -852, -780, -707, -634, -562, -489, -416, -344, -271, -198, -126, 20, 92, 165, 238, 311, 383, 456, 529, 601, 674, 747, 819, 892, 965, 1037, 1110, 1183, 1255, 1328, 1401, 1474, 1546, 1619, 1692, 1764, 1837, 1910, 1982, 2055, 2128, 2200, 2273, 2346, 2418, 2491, 2564, 2636, 2709, 2782, 2855, 2927, 3000, 3073, 3145, 3218, 3291, 3363, 3436, 3509]<br>[2430], [-3542, -3469, -3396, -3324, -3251, -3178, -3106, -3033, -2960, -2888, -2815, -2742, -2670, -2597, -2524, -2452, -2379, -2306, -2234, -2161, -2088, -2015, -1943, -1870, -1797, -1725, -1652, -1579, -1507, -1434, -1361, -1289, -1216, -1143, -1071, -998, -925, -852, -780, -707, -634, -562, -344, -271, -126, -53, 20, 311, 456, 529, 601, 674, 747, 819, 892, 965, 1037, 1110, 1183, 1255, 1328, 1401, 1473, 1546, 1619, 1692, 1764, 1837, 1910, 1982, 2055, 2128, 2200, 2273, 2346, 2418, 2491, 2564, 2636, 2709, 2782, 2855, 2927, 3000, 3073, 3145, 3218, 3291, 3363, 3436, 3509]<br>[5000], [-3542, -3469, -3396, -3324, -3251, -3178, -3106, -3033, -2960, -2888, -2815, -2742, -2670, -2597, -2524, -2452, -2379, -2306, -2233, -2161, -2088, -2015, -1943, -1870, -1797, -1725, -1652, -1579, -1507, -1434, -1361, -1289, -1216, -1143, -1070, -998, -925, -852, -707, -562, -489, -416, -344, 311, 383, 456, 529, 601, 819, 892, 965, 1037, 1110, 1183, 1255, 1328, 1401, 1474, 1546, 1619, 1692, 1764, 1837, 1910, 1982, 2055, 2128, 2200, 2273, 2346, 2418, 2491, 2564, 2636, 2709, 2782, 2855, 2927, 3000, 3073, 3145, 3218, 3291, 3363, 3436, 3509] |
| TAC, 25ºC<br>$T_{EX}$ = 100 ms<br>600 MHz | [30], [-3586, -3513, -3441, -3368, -3295, -3222, -3150, -3077, -3004, -2932, -2859, -2786, -2714, -2641, -2568, -2496, -2423, -2350, -2278, -2205, -2132, -2060, -1987, -1914, -1841, -1769, -1696, -1623, -1551, -1478, -1405, -1333, -1260, -1187, -1115, -1042, -969, -897, -824, -751, -678, -606, -533, -460, -388, -315, -242, -170, -97, -24, 48, 121, 194, 266, 339, 412, 485, 557, 630, 703, 775, 848, 921, 993, 1066, 1139, 1211, 1284, 1357, 1429, 1502, 1575, 1647, 1720, 1793, 1866, 1938, 2011, 2084, 2156, 2229, 2302, 2374, 2447, 2520, 2592, 2665, 2738, 2810, 2883, 2956, 3029, 3101, 3174, 3247, 3319, 3392]<br>[90], [-3586, -3513, -3441, -3368, -3295, -3222, -3150, -3077, -3004, -2932, -2859, -2786, -2714, -2641, -2568, -2496, -2423, -2350, -2278, -2205, -2132, -2060, -1987, -1914, -1841, -1769, -1696, -1623, -1551, -1478, -1405, -1333, -1260, -1187, -1115, -1042, -969, -897, -824, -751, -678, -606, -533, -460, -388, -315, -242, -170, -97, -24, 48, 121, 194, 266, 339, 412, 485, 557, 630, 703, 775, 848, 921, 993, 1066, 1139, 1211, 1284, 1357, 1429, 1502, 1575, 1647, 1720, 1793, 1866, 1938, 2011, 2084, 2156, 2229, 2302, 2374, 2447, 2520, 2592, 2665, 2738, 2810, 2883, 2956, 3029, 3101, 3174, 3247, 3319, 3392] |

| | |
|---|---|
| | [270], [-3586, -3513, -3441, -3368, -3295, -3222, -3150, -3077, -3004, -2932, -2859, -2786, -2714, -2641, -2568, -2496, -2423, -2350, -2278, -2205, -2132, -2059, -1987, -1914, -1841, -1769, -1696, -1623, -1551, -1478, -1405, -1333, -1260, -1187, -1115, -1042, -969, -896, -824, -751, -678, -606, -533, -460, -388, -315, -242, -170, -97, 48, 121, 194, 266, 339, 412, 485, 557, 630, 703, 775, 848, 921, 993, 1066, 1139, 1211, 1284, 1357, 1429, 1502, 1575, 1648, 1720, 1793, 1866, 1938, 2011, 2084, 2156, 2229, 2302, 2374, 2447, 2520, 2592, 2665, 2738, 2810, 2883, 2956, 3029, 3101, 3174, 3247, 3319, 3392]<br>[810], [-3586, -3513, -3441, -3368, -3295, -3222, -3150, -3077, -3004, -2932, -2859, -2786, -2714, -2641, -2568, -2496, -2423, -2350, -2278, -2205, -2132, -2059, -1987, -1914, -1841, -1769, -1696, -1623, -1551, -1478, -1405, -1333, -1260, -1187, -1115, -1042, -969, -896, -824, -751, -678, -606, -533, -460, -388, -315, -242, -97, 194, 266, 339, 412, 485, 557, 630, 703, 775, 848, 921, 993, 1066, 1139, 1211, 1284, 1357, 1429, 1502, 1575, 1648, 1720, 1793, 1866, 1938, 2011, 2084, 2156, 2229, 2302, 2374, 2447, 2520, 2592, 2665, 2738, 2810, 2883, 2956, 3029, 3101, 3174, 3247, 3319, 3392]<br>[2430], [-3586, -3513, -3441, -3368, -3295, -3222, -3150, -3077, -3004, -2932, -2859, -2786, -2714, -2641, -2568, -2496, -2423, -2350, -2278, -2205, -2132, -2059, -1987, -1914, -1841, -1769, -1696, -1623, -1551, -1478, -1405, -1333, -1260, -1187, -1115, -1042, -969, -896, -824, -751, -678, -460, -388, -315, -242, -170, -97, 48, 121, 194, 485, 557, 630, 703, 775, 848, 921, 993, 1066, 1139, 1211, 1284, 1357, 1429, 1502, 1575, 1648, 1720, 1793, 1866, 1938, 2011, 2084, 2156, 2229, 2302, 2374, 2447, 2520, 2592, 2665, 2738, 2811, 2883, 2956, 3029, 3101, 3174, 3247, 3319, 3392]<br>[5000], [-3586, -3513, -3441, -3368, -3295, -3222, -3150, -3077, -3004, -2932, -2859, -2786, -2714, -2641, -2568, -2496, -2423, -2350, -2278, -2205, -2132, -2060, -1987, -1914, -1841, -1769, -1696, -1623, -1551, -1478, -1405, -1333, -1260, -1187, -1115, -1042, -969, -606, -533, -388, -97, 48, 339, 484, 703, 848, 921, 993, 1066, 1139, 1211, 1284, 1357, 1429, 1502, 1575, 1647, 1720, 1793, 1866, 1938, 2011, 2084, 2156, 2229, 2302, 2374, 2447, 2520, 2592, 2665, 2738, 2810, 2883, 2956, 3029, 3101, 3174, 3247, 3319, 3392] |
| TAC, 25ºC<br>$T_{EX}$ = 80 ms<br>800 MHz | [100], [-4674, -4503, -4331, -4160, -3988, -3817, -3645, -3474, -3302, -3131, -2959, -2788, -2616, -2445, -2273, -2175, -2077, -1979, -1881, -1783, -1685, -1587, -1489, -1391, -1293, -1195, -1097, -999, -901, -803, -705, -607, -509, -411, -313, -215, -117, -19, 79, 177, 275, 373, 471, 569, 667, 765, 863, 961, 1059, 1157, 1255, 1353, 1451, 1549, 1647, 1745, 1843, 1941, 2039, 2137, 2235, 2333, 2431, 2529, 2701, 2872, 3044, 3215, 3387, 3558, 3730, 3901, 4073, 4244, 4416, 4587, 4759]<br>[250], [-4674, -4502, -4331, -4159, -3988, -3816, -3645, -3473, -3302, -3130, -2959, -2787, -2616, -2444, -2273, -2175, -2077, -1979, -1881, -1783, -1685, -1587, -1489, -1391, -1293, -1195, -1097, -999, -901, -803, -705, -607, -509, -411, -313, -215, -117, -19, 79, 177, 275, 373, 471, 569, 667, 765, 863, 961, 1059, 1157, 1255, 1353, 1451, 1549, 1647, 1745, 1843, 1941, 2039, 2137, 2235, 2333, |

| | |
|---|---|
| | 2431, 2529, 2701, 2872, 3044, 3215, 3387, 3558, 3730, 3901, 4073, 4244, 4416, 4587, 4759]<br>[500], [-4674, -4502, -4331, -4159, -3988, -3816, -3645, -3473, -3302, -3130, -2959, -2787, -2616, -2444, -2273, -2175, -2077, -1979, -1881, -1783, -1685, -1587, -1489, -1391, -1293, -1195, -1097, -999, -901, -803, -705, -607, -509, -411, -313, -215, -117, -19, 79, 177, 275, 373, 471, 569, 667, 765, 863, 961, 1059, 1157, 1255, 1353, 1451, 1549, 1647, 1745, 1843, 1941, 2039, 2137, 2235, 2333, 2431, 2529, 2701, 2872, 3044, 3215, 3387, 3558, 3730, 3901, 4073, 4244, 4416, 4587, 4759]<br>[1500], [-4674, -4503, -4331, -4160, -3988, -3817, -3645, -3474, -3302, -3131, -2959, -2788, -2616, -2445, -2273, -2175, -2077, -1979, -1881, -1783, -1685, -1587, -1489, -1391, -1293, -1195, -1097, -999, -901, -803, -705, -607, -509, -411, -313, -215, -117, -19, 275, 373, 471, 569, 667, 765, 863, 961, 1059, 1157, 1255, 1353, 1451, 1549, 1647, 1745, 1843, 1941, 2039, 2137, 2235, 2333, 2431, 2529, 2701, 2872, 3044, 3215, 3387, 3558, 3730, 3901, 4073, 4244, 4416, 4587, 4759]<br>[2000], [-4674, -4503, -4331, -4160, -3988, -3817, -3645, -3474, -3302, -3130, -2959, -2787, -2616, -2444, -2273, -2175, -2077, -1979, -1881, -1783, -1685, -1587, -1489, -1391, -1293, -1195, -1097, -999, -901, -803, -705, -607, -509, -411, -313, 177, 275, 373, 471, 569, 667, 765, 863, 961, 1059, 1157, 1255, 1353, 1451, 1549, 1647, 1745, 1843, 1941, 2039, 2137, 2235, 2333, 2431, 2529, 2701, 2872, 3044, 3215, 3387, 3558, 3730, 3901, 4073, 4244, 4416, 4587, 4759]<br>[3000], [-4674, -4503, -4331, -4160, -3988, -3817, -3645, -3474, -3302, -3131, -2959, -2788, -2616, -2445, -2273, -2175, -2077, -1979, -1881, -1783, -1685, -1587, -1489, -1391, -1293, -1195, -1097, -999, -901, -803, -705, -607, -509, -411, -117, 79, 177, 275, 373, 471, 569, 667, 765, 863, 961, 1059, 1157, 1255, 1353, 1451, 1549, 1647, 1745, 1843, 1941, 2039, 2137, 2235, 2333, 2431, 2529, 2701, 2872, 3044, 3215, 3387, 3558, 3730, 3901, 4073, 4244, 4416, 4587, 4759] |
| TAC, 20ºC<br>$T_{EX}$ = 80 ms<br>800 MHz | [200], [-4695, -4524, -4352, -4181, -4009, -3838, -3666, -3495, -3323, -3152, -2980, -2809, -2637, -2466, -2294, -2196, -2098, -2000, -1902, -1804, -1706, -1608, -1510, -1412, -1314, -1216, -1118, -1020, -922, -824, -726, -628, -530, -432, -334, -236, -138, 58, 156, 254, 352, 450, 548, 646, 744, 842, 940, 1038, 1136, 1234, 1332, 1430, 1528, 1626, 1724, 1822, 1920, 2018, 2116, 2214, 2312, 2410, 2508, 2680, 2851, 3023, 3194, 3366, 3537, 3709, 3880, 4052, 4223, 4395, 4566, 4738]<br>[500], [-4695, -4524, -4352, -4181, -4009, -3838, -3666, -3495, -3323, -3152, -2980, -2809, -2637, -2466, -2294, -2196, -2098, -2000, -1902, -1804, -1706, -1608, -1510, -1412, -1314, -1216, -1118, -1020, -922, -824, -726, -628, -530, -432, -334, -236, -138, 156, 254, 352, 450, 548, 646, 744, 842, 940, 1038, 1136, 1234, 1332, 1430, 1528, 1626, 1724, 1822, 1920, 2018, 2116, 2214, 2312, 2410, 2508, 2680, 2851, 3023, 3194, 3366, 3537, 3709, 3880, 4052, 4223, 4395, 4566, 4738] |

| | |
|---|---|
| | [1000], [-4695, -4524, -4352, -4181, -4009, -3838, -3666, -3495, -3323, -3152, -2980, -2809, -2637, -2466, -2294, -2196, -2098, -2000, -1902, -1804, -1706, -1608, -1510, -1412, -1314, -1216, -1118, -1020, -922, -824, -726, -628, -530, -432, -334, -236, 58, 156, 254, 352, 450, 548, 646, 744, 842, 940, 1038, 1136, 1234, 1332, 1430, 1528, 1626, 1724, 1822, 1920, 2018, 2116, 2214, 2312, 2410, 2508, 2680, 2851, 3023, 3194, 3366, 3537, 3709, 3880, 4052, 4223, 4395, 4566, 4738]<br>[2000], [-4695, -4524, -4352, -4181, -4009, -3838, -3666, -3494, -3323, -3151, -2980, -2808, -2637, -2465, -2294, -2196, -2098, -2000, -1902, -1804, -1706, -1608, -1510, -1412, -1314, -1216, -1118, -1020, -922, -824, -726, -628, -530, -432, -334, -138, 352, 450, 548, 646, 744, 842, 940, 1038, 1136, 1234, 1332, 1430, 1528, 1626, 1724, 1822, 1920, 2018, 2116, 2214, 2312, 2410, 2508, 2680, 2851, 3023, 3194, 3366, 3537, 3709, 3880, 4052, 4223, 4395, 4566, 4738]<br>[4000], [-4695, -4524, -4352, -4181, -4009, -3838, -3666, -3495, -3323, -3152, -2980, -2809, -2637, -2466, -2294, -2196, -2098, -2000, -1902, -1804, -1706, -1608, -1510, -1412, -1314, -1216, -1118, -1020, -922, -824, -726, -628, -530, -432, -334, -236, -138, -40, 254, 352, 450, 548, 744, 842, 940, 1038, 1136, 1234, 1332, 1430, 1528, 1626, 1724, 1822, 1920, 2018, 2116, 2214, 2312, 2410, 2508, 2680, 2851, 3023, 3194, 3366, 3537, 3709, 3880, 4052, 4223, 4395, 4566, 4738] |
| TAC, 15ºC<br>$T_{EX}$ = 100 ms<br>800 MHz | [200], [-4678, -4506, -4335, -4163, -3992, -3820, -3649, -3477, -3306, -3134, -2963, -2791, -2620, -2448, -2277, -2179, -2081, -1983, -1885, -1787, -1689, -1591, -1493, -1395, -1297, -1199, -1101, -1003, -905, -807, -709, -611, -513, -415, -317, -219, -121, -23, 75, 173, 271, 369, 467, 565, 663, 761, 859, 957, 1055, 1153, 1251, 1349, 1447, 1545, 1643, 1741, 1839, 1937, 2035, 2133, 2231, 2329, 2427, 2525, 2697, 2868, 3040, 3211, 3383, 3554, 3726, 3897, 4069, 4240, 4412, 4583, 4755]<br>[500], [-4678, -4506, -4335, -4163, -3992, -3820, -3649, -3477, -3306, -3134, -2963, -2791, -2620, -2448, -2277, -2179, -2081, -1983, -1885, -1787, -1689, -1591, -1493, -1395, -1297, -1199, -1101, -1003, -905, -807, -709, -611, -513, -415, -317, -219, -121, -23, 173, 271, 369, 467, 565, 663, 761, 859, 957, 1055, 1153, 1251, 1349, 1447, 1545, 1643, 1741, 1839, 1937, 2035, 2133, 2231, 2329, 2427, 2525, 2697, 2868, 3040, 3211, 3383, 3554, 3726, 3897, 4069, 4240, 4412, 4583, 4755]<br>[1000], [-4678, -4506, -4335, -4163, -3992, -3820, -3649, -3477, -3306, -3134, -2963, -2791, -2620, -2448, -2277, -2179, -2081, -1983, -1885, -1787, -1689, -1591, -1493, -1395, -1297, -1199, -1101, -1003, -905, -807, -709, -611, -513, -415, -317, -219, -121, -23, 173, 271, 369, 467, 565, 663, 761, 859, 957, 1055, 1153, 1251, 1349, 1447, 1545, 1643, 1741, 1839, 1937, 2035, 2133, 2231, 2329, 2427, 2525, 2697, 2868, 3040, 3211, 3383, 3554, 3726, 3897, 4069, 4240, 4412, 4583, 4755]<br>[2000], [-4678, -4506, -4335, -4163, -3992, -3820, -3649, -3477, -3306, -3134, -2963, -2791, -2620, -2448, -2277, -2179, -2081, -1983, -1885, -1787, -1689, -1591, -1493, -1395, -1297, -1199, -1101, -1003, -905, -807, -709, -611, -513, - |

| | |
|---|---|
| | 415, -317, 565, 663, 761, 859, 957, 1055, 1153, 1251, 1349, 1447, 1545, 1643, 1741, 1839, 1937, 2035, 2133, 2231, 2329, 2427, 2525, 2697, 2868, 3040, 3211, 3383, 3554, 3726, 3897, 4069, 4240, 4412, 4583, 4755] |
| TAC, 12.5ºC $T_{EX}$ = 100 ms<br>800 MHz | [25], [-4688, -4516, -4345, -4173, -4002, -3830, -3659, -3487, -3316, -3144, -2973, -2801, -2630, -2458, -2287, -2189, -2091, -1993, -1895, -1797, -1699, -1601, -1503, -1405, -1307, -1209, -1111, -1013, -915, -817, -719, -621, -523, -425, -327, -229, -131, -33, 65, 163, 261, 359, 457, 555, 653, 751, 849, 947, 1045, 1143, 1241, 1339, 1437, 1535, 1633, 1731, 1829, 1927, 2025, 2123, 2221, 2319, 2417, 2515, 2687, 2858, 3030, 3201, 3373, 3544, 3716, 3887, 4059, 4230, 4402, 4573, 4745]<br>[100], [-4688, -4516, -4345, -4173, -4002, -3830, -3659, -3487, -3316, -3144, -2973, -2801, -2630, -2458, -2287, -2189, -2091, -1993, -1895, -1797, -1699, -1601, -1503, -1405, -1307, -1209, -1111, -1013, -915, -817, -719, -621, -523, -425, -327, -229, -131, -33, 65, 163, 261, 359, 457, 555, 653, 751, 849, 947, 1045, 1143, 1241, 1339, 1437, 1535, 1633, 1731, 1829, 1927, 2025, 2123, 2221, 2319, 2417, 2515, 2687, 2858, 3030, 3201, 3373, 3544, 3716, 3887, 4059, 4230, 4402, 4573, 4745]<br>[250], [-4688, -4516, -4345, -4173, -4002, -3830, -3659, -3487, -3316, -3144, -2973, -2801, -2630, -2458, -2287, -2189, -2091, -1993, -1895, -1797, -1699, -1601, -1503, -1405, -1307, -1209, -1111, -1013, -915, -817, -719, -621, -523, -425, -327, -229, -131, -33, 65, 163, 261, 359, 457, 555, 653, 751, 849, 947, 1045, 1143, 1241, 1339, 1437, 1535, 1633, 1731, 1829, 1927, 2025, 2123, 2221, 2319, 2417, 2515, 2687, 2858, 3030, 3201, 3373, 3544, 3716, 3888, 4059, 4231, 4402, 4574, 4745]<br>[1250], [-4688, -4516, -4345, -4173, -4002, -3830, -3659, -3487, -3316, -3144, -2973, -2801, -2630, -2458, -2287, -2189, -2091, -1993, -1895, -1797, -1699, -1601, -1503, -1405, -1307, -1209, -1111, -1013, -915, -817, -719, -621, -523, -425, -327, 261, 359, 457, 555, 653, 751, 849, 947, 1045, 1143, 1241, 1339, 1437, 1535, 1633, 1731, 1829, 1927, 2025, 2123, 2221, 2319, 2417, 2515, 2687, 2858, 3030, 3201, 3373, 3544, 3716, 3887, 4059, 4230, 4402, 4573, 4745]<br>[1750], [-4688, -4516, -4345, -4173, -4002, -3830, -3659, -3487, -3316, -3144, -2973, -2801, -2630, -2458, -2287, -2189, -2091, -1993, -1895, -1797, -1699, -1601, -1503, -1405, -1307, -1209, -1111, -1013, -915, -817, -719, -621, -523, -425, -327, -131, 65, 261, 359, 457, 555, 653, 751, 849, 947, 1045, 1143, 1241, 1339, 1437, 1535, 1633, 1731, 1829, 1927, 2025, 2123, 2221, 2319, 2417, 2515, 2687, 2858, 3030, 3202, 3373, 3545, 3716, 3888, 4059, 4231, 4402, 4574, 4745] |
| TAC, 10 ºC $T_{EX}$ = 100 ms<br>800 MHz | [25], [-4660, -4489, -4317, -4146, -3974, -3803, -3631, -3460, -3288, -3117, -2945, -2774, -2602, -2431, -2259, -2161, -2063, -1965, -1867, -1769, -1671, -1573, -1475, -1377, -1279, -1181, -1083, -985, -887, -789, -691, -593, -495, -397, -299, -201, -103, -5, 93, 191, 289, 387, 485, 583, 681, 779, 877, 975, 1073, 1171, 1269, 1367, 1465, 1563, 1661, 1759, 1857, 1955, 2053, 2151, 2249, 2347, |

| | |
|---|---|
| | 2445, 2543, 2714, 2886, 3057, 3229, 3400, 3572, 3743, 3915, 4086, 4258, 4429, 4601, 4772]<br>[50], [-4660, -4489, -4317, -4146, -3974, -3803, -3631, -3460, -3288, -3117, -2945, -2774, -2602, -2431, -2259, -2161, -2063, -1965, -1867, -1769, -1671, -1573, -1475, -1377, -1279, -1181, -1083, -985, -887, -789, -691, -593, -495, -397, -299, -201, -103, 93, 191, 289, 387, 485, 583, 681, 779, 877, 975, 1073, 1171, 1269, 1367, 1465, 1563, 1661, 1759, 1857, 1955, 2053, 2151, 2249, 2347, 2445, 2543, 2715, 2886, 3058, 3229, 3401, 3572, 3744, 3915, 4087, 4258, 4430, 4601, 4773]<br>[150], [-4660, -4489, -4317, -4146, -3974, -3803, -3631, -3460, -3288, -3117, -2945, -2774, -2602, -2431, -2259, -2161, -2063, -1965, -1867, -1769, -1671, -1573, -1475, -1377, -1279, -1181, -1083, -985, -887, -789, -691, -593, -495, -397, -299, -201, -103, 93, 191, 289, 387, 485, 583, 681, 779, 877, 975, 1073, 1171, 1269, 1367, 1465, 1563, 1661, 1759, 1857, 1955, 2053, 2151, 2249, 2347, 2445, 2543, 2714, 2886, 3057, 3229, 3400, 3572, 3743, 3915, 4086, 4258, 4429, 4601, 4772]<br>[300], [-4660, -4489, -4317, -4146, -3974, -3803, -3631, -3460, -3288, -3117, -2945, -2774, -2602, -2431, -2259, -2161, -2063, -1965, -1867, -1769, -1671, -1573, -1475, -1377, -1279, -1181, -1083, -985, -887, -789, -691, -593, -495, -397, -299, -201, -103, -5, 93, 191, 289, 387, 485, 583, 681, 779, 877, 975, 1073, 1171, 1269, 1367, 1465, 1563, 1661, 1759, 1857, 1955, 2053, 2151, 2249, 2347, 2445, 2543, 2714, 2886, 3057, 3229, 3401, 3572, 3744, 3915, 4087, 4258, 4430, 4601, 4773]<br>[500], [-4660, -4489, -4317, -4146, -3974, -3803, -3631, -3460, -3288, -3117, -2945, -2774, -2602, -2431, -2259, -2161, -2063, -1965, -1867, -1769, -1671, -1573, -1475, -1377, -1279, -1181, -1083, -985, -887, -789, -691, -593, -495, -397, -299, -201, -103, 93, 191, 289, 387, 485, 583, 681, 779, 877, 975, 1073, 1171, 1269, 1367, 1465, 1563, 1661, 1759, 1857, 1955, 2053, 2151, 2249, 2347, 2445, 2543, 2715, 2886, 3058, 3229, 3401, 3572, 3744, 3915, 4087, 4258, 4430, 4601, 4773]<br>[1000], [-4660, -4489, -4317, -4146, -3974, -3803, -3631, -3460, -3288, -3117, -2945, -2774, -2602, -2431, -2259, -2161, -2063, -1965, -1867, -1769, -1671, -1573, -1475, -1377, -1279, -1181, -1083, -985, -887, -789, -691, -593, -495, -397, -299, -201, 93, 191, 289, 387, 485, 583, 681, 779, 877, 975, 1073, 1171, 1269, 1367, 1465, 1563, 1661, 1759, 1857, 1955, 2053, 2151, 2249, 2347, 2445, 2543, 2714, 2886, 3057, 3229, 3400, 3572, 3743, 3915, 4086, 4258, 4429, 4601, 4772] |
| GAG, 25ºC<br>$T_{EX}$ = 100 ms<br>700 MHz | [30], [-3509, -3437, -3364, -3291, -3219, -3146, -3073, -3001, -2928, -2855, -2782, -2710, -2637, -2564, -2492, -2419, -2346, -2274, -2201, -2128, -2056, -1983, -1910, -1838, -1765, -1692, -1619, -1547, -1474, -1401, -1329, -1256, -1183, -1111, -1038, -965, -893, -820, -747, -675, -602, -529, -457, -384, -311, -238, -166, -93, 52, 125, 198, 270, 343, 416, 488, 561, 634, 706, 779, 852, 925, 997, 1070, 1143, 1215, 1288, 1361, 1433, 1506, 1579, 1651, 1724, 1797, 1869, |

| | |
|---|---|
| | 1942, 2015, 2087, 2160, 2233, 2306, 2378, 2451, 2524, 2596, 2669, 2742, 2814, 2887, 2960, 3032, 3105, 3178, 3250, 3323, 3396, 3469, 3541]<br>[90], [-3509, -3437, -3364, -3291, -3219, -3146, -3073, -3001, -2928, -2855, -2783, -2710, -2637, -2564, -2492, -2419, -2346, -2274, -2201, -2128, -2056, -1983, -1910, -1838, -1765, -1692, -1620, -1547, -1474, -1402, -1329, -1256, -1183, -1111, -1038, -965, -893, -820, -747, -675, -602, -529, -457, -384, -311, -239, -166, -93, -20, 125, 198, 270, 343, 416, 488, 561, 634, 706, 779, 852, 924, 997, 1070, 1143, 1215, 1288, 1361, 1433, 1506, 1579, 1651, 1724, 1797, 1869, 1942, 2015, 2087, 2160, 2233, 2305, 2378, 2451, 2524, 2596, 2669, 2742, 2814, 2887, 2960, 3032, 3105, 3178, 3250, 3323, 3396, 3468, 3541]<br>[270], [-3509, -3437, -3364, -3291, -3219, -3146, -3073, -3001, -2928, -2855, -2783, -2710, -2637, -2564, -2492, -2419, -2346, -2274, -2201, -2128, -2056, -1983, -1910, -1838, -1765, -1692, -1620, -1547, -1474, -1401, -1329, -1256, -1183, -1111, -1038, -965, -893, -820, -747, -675, -602, -529, -457, -384, -311, -239, -166, -93, 52, 125, 198, 270, 343, 416, 488, 561, 634, 706, 779, 852, 924, 997, 1070, 1143, 1215, 1288, 1361, 1433, 1506, 1579, 1651, 1724, 1797, 1869, 1942, 2015, 2087, 2160, 2233, 2306, 2378, 2451, 2524, 2596, 2669, 2742, 2814, 2887, 2960, 3032, 3105, 3178, 3250, 3323, 3396, 3468, 3541]<br>[810], [-3509, -3437, -3364, -3291, -3219, -3146, -3073, -3001, -2928, -2855, -2782, -2710, -2637, -2564, -2492, -2419, -2346, -2274, -2201, -2128, -2056, -1983, -1910, -1838, -1765, -1692, -1619, -1547, -1474, -1401, -1329, -1256, -1183, -1111, -1038, -965, -893, -820, -747, -675, -602, -529, -457, -384, -311, -238, -166, -93, 125, 198, 270, 343, 416, 488, 561, 634, 706, 779, 852, 925, 997, 1070, 1143, 1215, 1288, 1361, 1433, 1506, 1579, 1651, 1724, 1797, 1869, 1942, 2015, 2087, 2160, 2233, 2306, 2378, 2451, 2524, 2596, 2669, 2742, 2814, 2887, 2960, 3032, 3105, 3178, 3250, 3323, 3396, 3469, 3541]<br>[2430], [-3509, -3437, -3364, -3291, -3219, -3146, -3073, -3001, -2928, -2855, -2783, -2710, -2637, -2564, -2492, -2419, -2346, -2274, -2201, -2128, -2056, -1983, -1910, -1838, -1765, -1692, -1620, -1547, -1474, -1402, -1329, -1256, -1183, -1111, -1038, -965, -893, -820, -747, -675, -602, -529, -457, -384, -311, -166, 125, 198, 343, 416, 488, 634, 706, 779, 852, 924, 997, 1070, 1142, 1215, 1288, 1361, 1433, 1506, 1579, 1651, 1724, 1797, 1869, 1942, 2015, 2087, 2160, 2233, 2305, 2378, 2451, 2524, 2596, 2669, 2742, 2814, 2887, 2960, 3032, 3105, 3178, 3250, 3323, 3396, 3468, 3541]<br>[5000], [-3509, -3437, -3364, -3291, -3219, -3146, -3073, -3001, -2928, -2855, -2782, -2710, -2637, -2564, -2492, -2419, -2346, -2274, -2201, -2128, -2056, -1983, -1910, -1838, -1765, -1692, -1620, -1547, -1474, -1401, -1329, -1256, -1183, -1111, -965, -893, -820, -747, -675, -602, -529, -457, -384, -238, -20, 52, 125, 198, 270, 343, 561, 634, 779, 852, 924, 997, 1070, 1143, 1215, 1288, 1361, 1433, 1506, 1579, 1651, 1724, 1797, 1869, 1942, 2015, 2087, 2160, 2233, 2306, 2378, 2451, 2524, 2596, 2669, 2742, 2814, 2887, 2960, 3032, 3105, 3178, 3250, 3323, 3396, 3469, 3541] |
| CAG, 25ºC | [30], [-3519, -3446, -3374, -3301, -3228, -3156, -3083, -3010, -2938, -2865, -2792, -2719, -2647, -2574, -2501, -2429, -2356, -2283, -2211, -2138, -2065, - |

| | |
|---|---|
| $T_{EX}$ = 100 ms<br>600 MHz | 1993, -1920, -1847, -1775, -1702, -1629, -1557, -1484, -1411, -1338, -1266, -1193, -1120, -1048, -975, -902, -830, -757, -684, -612, -539, -466, -394, -321, -248, -175, -103, -30, 43, 115, 188, 261, 333, 406, 479, 551, 624, 697, 769, 842, 915, 988, 1060, 1133, 1206, 1278, 1351, 1424, 1496, 1569, 1642, 1714, 1787, 1860, 1932, 2005, 2078, 2150, 2223, 2296, 2369, 2441, 2514, 2587, 2659, 2732, 2805, 2877, 2950, 3023, 3095, 3168, 3241, 3313, 3386, 3459, 3532]<br>[90], [-3519, -3446, -3374, -3301, -3228, -3155, -3083, -3010, -2937, -2865, -2792, -2719, -2647, -2574, -2501, -2429, -2356, -2283, -2211, -2138, -2065, -1993, -1920, -1847, -1774, -1702, -1629, -1556, -1484, -1411, -1338, -1266, -1193, -1120, -1048, -975, -902, -830, -757, -684, -611, -539, -466, -393, -321, -248, -175, -103, -30, 43, 115, 188, 261, 333, 406, 479, 552, 624, 697, 770, 842, 915, 988, 1060, 1133, 1206, 1278, 1351, 1424, 1496, 1569, 1642, 1714, 1787, 1860, 1933, 2005, 2078, 2151, 2223, 2296, 2369, 2441, 2514, 2587, 2659, 2732, 2805, 2877, 2950, 3023, 3096, 3168, 3241, 3314, 3386, 3459, 3532]<br>[270], [-3519, -3446, -3374, -3301, -3228, -3156, -3083, -3010, -2938, -2865, -2792, -2719, -2647, -2574, -2501, -2429, -2356, -2283, -2211, -2138, -2065, -1993, -1920, -1847, -1775, -1702, -1629, -1556, -1484, -1411, -1338, -1266, -1193, -1120, -1048, -975, -902, -830, -757, -684, -612, -539, -466, -393, -321, -248, -175, -103, 188, 261, 333, 406, 479, 551, 624, 697, 769, 842, 915, 988, 1060, 1133, 1206, 1278, 1351, 1424, 1496, 1569, 1642, 1714, 1787, 1860, 1932, 2005, 2078, 2151, 2223, 2296, 2369, 2441, 2514, 2587, 2659, 2732, 2805, 2877, 2950, 3023, 3095, 3168, 3241, 3313, 3386, 3459, 3532]<br>[810], [-3519, -3446, -3374, -3301, -3228, -3156, -3083, -3010, -2937, -2865, -2792, -2719, -2647, -2574, -2501, -2429, -2356, -2283, -2211, -2138, -2065, -1993, -1920, -1847, -1775, -1702, -1629, -1556, -1484, -1411, -1338, -1266, -1193, -1120, -1048, -975, -902, -830, -757, -684, -612, -539, -466, -393, -321, -248, -30, 43, 188, 261, 333, 406, 479, 551, 624, 697, 770, 842, 915, 988, 1060, 1133, 1206, 1278, 1351, 1424, 1496, 1569, 1642, 1714, 1787, 1860, 1932, 2005, 2078, 2151, 2223, 2296, 2369, 2441, 2514, 2587, 2659, 2732, 2805, 2877, 2950, 3023, 3095, 3168, 3241, 3314, 3386, 3459, 3532]<br>[2430], [-3519, -3446, -3374, -3301, -3228, -3156, -3083, -3010, -2938, -2865, -2792, -2720, -2647, -2574, -2501, -2429, -2356, -2283, -2211, -2138, -2065, -1993, -1920, -1847, -1775, -1702, -1629, -1557, -1484, -1411, -1338, -1266, -1193, -1120, -1048, -975, -902, -830, -757, -684, -612, -539, -466, -394, -321, -176, -30, 115, 261, 406, 479, 551, 624, 697, 769, 842, 915, 987, 1060, 1133, 1206, 1278, 1351, 1424, 1496, 1569, 1642, 1714, 1787, 1860, 1932, 2005, 2078, 2150, 2223, 2296, 2368, 2441, 2514, 2587, 2659, 2732, 2805, 2877, 2950, 3023, 3095, 3168, 3241, 3313, 3386, 3459, 3531]<br>[5000], [-3519, -3446, -3374, -3301, -3228, -3156, -3083, -3010, -2938, -2865, -2792, -2720, -2647, -2574, -2501, -2429, -2356, -2283, -2211, -2138, -2065, -1993, -1920, -1847, -1775, -1702, -1629, -1557, -1484, -1411, -1338, -1266, -1193, -1120, -1048, -975, -902, -830, -757, -684, -612, -539, -176, -30, 115, 261, 551, 624, 697, 769, 842, 915, 987, 1060, 1133, 1206, 1278, 1351, 1424, 1496, 1569, 1642, 1714, 1787, 1860, 1932, 2005, 2078, 2150, 2223, 2296, 2369, |

| | |
|---|---|
| | 2441, 2514, 2587, 2659, 2732, 2805, 2877, 2950, 3023, 3095, 3168, 3241, 3313, 3386, 3459, 3531] |
| CAG, 25ºC<br>$T_{EX}$ = 100 ms<br>900 MHz | [250], [-5312, -5067, -4821, -4576, -4331, -4085, -3840, -3594, -3349, -3103, -2858, -2612, -2492, -2372, -2252, -2132, -2012, -1892, -1772, -1652, -1532, -1412, -1292, -1172, -1052, -932, -812, -692, -572, -452, -332, -212, -92, 148, 268, 388, 508, 628, 748, 868, 988, 1108, 1228, 1348, 1468, 1588, 1708, 1828, 1948, 2068, 2188, 2308, 2428, 2548, 2668, 2788, 3033, 3279, 3524, 3770, 4015, 4261, 4506, 4752, 4997, 5243]<br>[1000], [-5313, -5067, -4822, -4576, -4331, -4085, -3840, -3594, -3349, -3103, -2858, -2612, -2492, -2372, -2252, -2132, -2012, -1892, -1772, -1652, -1532, -1412, -1292, -1172, -1052, -932, -812, -692, -572, -452, -332, -212, 148, 268, 388, 508, 628, 748, 868, 988, 1108, 1228, 1348, 1468, 1588, 1708, 1828, 1948, 2068, 2188, 2308, 2428, 2548, 2668, 2788, 3033, 3279, 3524, 3770, 4015, 4261, 4506, 4751, 4997, 5242]<br>[2000], [-5313, -5067, -4822, -4576, -4331, -4085, -3840, -3594, -3349, -3103, -2858, -2613, -2493, -2373, -2253, -2133, -2013, -1893, -1773, -1652, -1532, -1412, -1292, -1172, -1052, -932, -812, -692, -572, -452, -212, 268, 508, 628, 748, 868, 988, 1108, 1228, 1348, 1468, 1588, 1708, 1828, 1948, 2068, 2188, 2308, 2428, 2548, 2668, 2788, 3033, 3279, 3524, 3770, 4015, 4260, 4506, 4751, 4997, 5242]<br>[4000], [-5312, -5067, -4821, -4576, -4330, -4085, -3840, -3594, -3349, -3103, -2858, -2612, -2492, -2372, -2252, -2132, -2012, -1892, -1772, -1652, -1532, -1412, -1292, -1172, -1052, -932, -812, -692, -332, -92, 268, 388, 508, 628, 748, 868, 988, 1108, 1228, 1348, 1468, 1588, 1708, 1828, 1948, 2068, 2188, 2308, 2428, 2548, 2668, 2788, 3033, 3279, 3524, 3770, 4015, 4261, 4506, 4752, 4997, 5243] |
| CAG, 20ºC<br>$T_{EX}$ = 100 ms<br>900 MHz | [100], [-5339, -5093, -4848, -4602, -4357, -4111, -3866, -3620, -3375, -3129, -2884, -2639, -2519, -2399, -2279, -2159, -2039, -1919, -1799, -1679, -1559, -1438, -1318, -1198, -1078, -958, -838, -718, -598, -478, -358, -238, -118, 2, 122, 242, 362, 482, 602, 722, 842, 962, 1082, 1202, 1322, 1442, 1562, 1682, 1802, 1922, 2042, 2162, 2282, 2402, 2522, 2642, 2762, 3007, 3253, 3498, 3743, 3989, 4234, 4480, 4725, 4971, 5216]<br>[500], [-5339, -5093, -4848, -4603, -4357, -4112, -3866, -3621, -3375, -3130, -2884, -2639, -2519, -2399, -2279, -2159, -2039, -1919, -1799, -1679, -1559, -1439, -1319, -1199, -1079, -959, -839, -719, -599, -479, -359, -239, 121, 241, 361, 481, 601, 721, 841, 961, 1081, 1201, 1321, 1441, 1561, 1681, 1801, 1921, 2041, 2161, 2281, 2401, 2521, 2641, 2761, 3007, 3252, 3498, 3743, 3989, 4234, 4480, 4725, 4971, 5216]<br>[1000], [-5339, -5093, -4848, -4602, -4357, -4111, -3866, -3620, -3375, -3130, -2884, -2639, -2519, -2399, -2279, -2159, -2039, -1919, -1799, -1679, -1559, -1439, -1319, -1199, -1079, -959, -839, -719, -599, -479, -358, -238, 2, 122, 242, 362, 482, 602, 722, 842, 962, 1082, 1202, 1322, 1442, 1562, 1682, 1802, 1922, 2042, 2162, 2282, 2402, 2522, 2642, 2762, 3007, 3253, 3498, 3743, 3989, 4234, 4480, 4725, 4971, 5216] |

| | |
|---|---|
| | [2000], [-5339, -5093, -4848, -4602, -4357, -4112, -3866, -3621, -3375, -3130, -2884, -2639, -2519, -2399, -2279, -2159, -2039, -1919, -1799, -1679, -1559, -1439, -1319, -1199, -1079, -959, -839, -719, -599, -479, -359, 241, 361, 481, 601, 721, 841, 961, 1081, 1201, 1321, 1441, 1561, 1681, 1801, 1921, 2041, 2161, 2281, 2401, 2521, 2641, 2761, 3007, 3252, 3498, 3743, 3989, 4234, 4480, 4725, 4971, 5216] |
| CAG, 15ºC<br>$T_{EX}$ = 100 ms<br>900 MHz | [100], [-5366, -5120, -4875, -4629, -4384, -4138, -3893, -3647, -3402, -3156, -2911, -2665, -2545, -2425, -2305, -2185, -2065, -1945, -1825, -1705, -1585, -1465, -1345, -1225, -1105, -985, -865, -745, -625, -505, -385, -265, -145, 95, 215, 335, 455, 575, 695, 815, 935, 1055, 1175, 1295, 1415, 1535, 1655, 1775, 1895, 2015, 2135, 2255, 2375, 2495, 2615, 2735, 2980, 3226, 3471, 3717, 3962, 4208, 4453, 4698, 4944, 5189]<br>[250], [-5366, -5120, -4875, -4629, -4384, -4138, -3893, -3648, -3402, -3157, -2911, -2666, -2546, -2426, -2306, -2186, -2066, -1946, -1826, -1706, -1586, -1466, -1346, -1226, -1106, -986, -866, -746, -626, -506, -386, -266, -146, -26, 94, 214, 334, 454, 574, 694, 814, 934, 1054, 1174, 1294, 1414, 1534, 1654, 1774, 1894, 2014, 2134, 2255, 2375, 2495, 2615, 2735, 2980, 3225, 3471, 3716, 3962, 4207, 4453, 4698, 4944, 5189]<br>[500], [-5366, -5121, -4875, -4630, -4384, -4139, -3893, -3648, -3402, -3157, -2911, -2666, -2546, -2426, -2306, -2186, -2066, -1946, -1826, -1706, -1586, -1466, -1346, -1226, -1106, -986, -866, -746, -626, -506, -386, -266, -146, -26, 94, 214, 334, 454, 574, 694, 814, 934, 1054, 1174, 1294, 1414, 1534, 1654, 1774, 1894, 2014, 2134, 2254, 2374, 2494, 2614, 2734, 2980, 3225, 3471, 3716, 3962, 4207, 4452, 4698, 4943, 5189]<br>[1000], [-5366, -5120, -4875, -4629, -4384, -4138, -3893, -3647, -3402, -3156, -2911, -2666, -2546, -2426, -2306, -2186, -2066, -1946, -1825, -1705, -1585, -1465, -1345, -1225, -1105, -985, -865, -745, -625, -505, -385, -265, -25, 95, 335, 455, 575, 695, 815, 935, 1055, 1175, 1295, 1415, 1535, 1655, 1775, 1895, 2015, 2135, 2255, 2375, 2495, 2615, 2735, 2980, 3226, 3471, 3717, 3962, 4207, 4453, 4698, 4944, 5189] |
| CAG, 10ºC<br>$T_{EX}$ = 100 ms<br>900 MHz | [50], [-5303, -5057, -4812, -4566, -4321, -4075, -3830, -3584, -3339, -3093, -2848, -2603, -2483, -2363, -2243, -2123, -2003, -1882, -1762, -1642, -1522, -1402, -1282, -1162, -1042, -922, -802, -682, -562, -442, -322, -202, -82, 38, 158, 278, 398, 518, 638, 758, 878, 998, 1118, 1238, 1358, 1478, 1598, 1718, 1838, 1958, 2078, 2198, 2318, 2438, 2558, 2678, 2798, 3043, 3289, 3534, 3780, 4025, 4270, 4516, 4761, 5007, 5252]<br>[200], [-5302, -5057, -4811, -4566, -4320, -4075, -3829, -3584, -3339, -3093, -2848, -2602, -2482, -2362, -2242, -2122, -2002, -1882, -1762, -1642, -1522, -1402, -1282, -1162, -1042, -922, -802, -682, -562, -442, -322, -202, -82, 38, 158, 278, 398, 518, 638, 758, 878, 998, 1118, 1238, 1358, 1478, 1598, 1718, 1838, 1958, 2078, 2198, 2318, 2438, 2558, 2678, 2798, 3043, 3289, 3534, 3780, 4025, 4271, 4516, 4762, 5007, 5253]<br>[500], [-5303, -5057, -4812, -4566, -4321, -4075, -3830, -3584, -3339, -3093, -2848, -2602, -2482, -2362, -2242, -2122, -2002, -1882, -1762, -1642, -1522, - |

| | |
|---|---|
| | 1402, -1282, -1162, -1042, -922, -802, -682, -562, -442, -322, -202, -82, 158, 278, 398, 518, 638, 758, 878, 998, 1118, 1238, 1358, 1478, 1598, 1718, 1838, 1958, 2078, 2198, 2318, 2438, 2558, 2678, 2798, 3043, 3289, 3534, 3780, 4025, 4270, 4516, 4761, 5007, 5252]<br>[1000], [-5302, -5057, -4811, -4566, -4320, -4075, -3829, -3584, -3338, -3093, -2847, -2602, -2482, -2362, -2242, -2122, -2002, -1882, -1762, -1642, -1522, -1402, -1282, -1162, -1042, -922, -802, -682, -562, -442, -322, -202, 38, 158, 278, 518, 638, 758, 878, 998, 1118, 1238, 1358, 1478, 1598, 1718, 1838, 1958, 2078, 2198, 2318, 2438, 2558, 2678, 2798, 3044, 3289, 3535, 3780, 4026, 4271, 4516, 4762, 5007, 5253] |
| GAC, 25ºC<br>$T_{EX}$ = 100 ms<br>600 MHz | [30], [-3577, -3504, -3431, -3359, -3286, -3213, -3141, -3068, -2995, -2923, -2850, -2777, -2705, -2632, -2559, -2486, -2414, -2341, -2268, -2196, -2123, -2050, -1978, -1905, -1832, -1760, -1687, -1614, -1542, -1469, -1396, -1323, -1251, -1178, -1105, -1033, -960, -887, -815, -742, -669, -597, -524, -451, -379, -306, -233, -161, -88, -15, 58, 130, 203, 276, 348, 421, 494, 566, 639, 712, 784, 857, 930, 1002, 1075, 1148, 1221, 1293, 1366, 1439, 1511, 1584, 1657, 1729, 1802, 1875, 1947, 2020, 2093, 2165, 2238, 2311, 2383, 2456, 2529, 2602, 2674, 2747, 2820, 2892, 2965, 3038, 3110, 3183, 3256, 3328, 3401, 3474, 3546]<br>[90], [-3577, -3504, -3431, -3359, -3286, -3213, -3141, -3068, -2995, -2923, -2850, -2777, -2704, -2632, -2559, -2486, -2414, -2341, -2268, -2196, -2123, -2050, -1978, -1905, -1832, -1760, -1687, -1614, -1542, -1469, -1396, -1323, -1251, -1178, -1105, -1033, -960, -887, -815, -742, -669, -597, -524, -451, -379, -306, -233, -160, -88, -15, 130, 203, 276, 348, 421, 494, 566, 639, 712, 784, 857, 930, 1002, 1075, 1148, 1221, 1293, 1366, 1439, 1511, 1584, 1657, 1729, 1802, 1875, 1947, 2020, 2093, 2165, 2238, 2311, 2384, 2456, 2529, 2602, 2674, 2747, 2820, 2892, 2965, 3038, 3110, 3183, 3256, 3328, 3401, 3474, 3547]<br>[270], [-3577, -3504, -3431, -3359, -3286, -3213, -3141, -3068, -2995, -2923, -2850, -2777, -2705, -2632, -2559, -2487, -2414, -2341, -2268, -2196, -2123, -2050, -1978, -1905, -1832, -1760, -1687, -1614, -1542, -1469, -1396, -1324, -1251, -1178, -1105, -1033, -960, -887, -815, -742, -669, -597, -524, -451, -379, -306, -233, -161, -88, 57, 130, 203, 276, 348, 421, 494, 566, 639, 712, 784, 857, 930, 1002, 1075, 1148, 1220, 1293, 1366, 1439, 1511, 1584, 1657, 1729, 1802, 1875, 1947, 2020, 2093, 2165, 2238, 2311, 2383, 2456, 2529, 2602, 2674, 2747, 2820, 2892, 2965, 3038, 3110, 3183, 3256, 3328, 3401, 3474, 3546]<br>[810], [-3577, -3504, -3431, -3359, -3286, -3213, -3141, -3068, -2995, -2923, -2850, -2777, -2705, -2632, -2559, -2486, -2414, -2341, -2268, -2196, -2123, -2050, -1978, -1905, -1832, -1760, -1687, -1614, -1542, -1469, -1396, -1324, -1251, -1178, -1105, -1033, -960, -887, -815, -742, -669, -597, -524, -451, -379, -306, -233, -161, -88, -15, 58, 203, 276, 348, 421, 494, 566, 639, 712, 784, 857, 930, 1002, 1075, 1148, 1221, 1293, 1366, 1439, 1511, 1584, 1657, 1729, 1802, 1875, 1947, 2020, 2093, 2165, 2238, 2311, 2383, 2456, 2529, 2602, 2674, 2747, 2820, 2892, 2965, 3038, 3110, 3183, 3256, 3328, 3401, 3474, 3546]<br>[2430], [-3577, -3504, -3431, -3359, -3286, -3213, -3141, -3068, -2995, -2923, -2850, -2777, -2704, -2632, -2559, -2486, -2414, -2341, -2268, -2196, -2123, - |

| | |
|---|---|
| | 2050, -1978, -1905, -1832, -1760, -1687, -1614, -1542, -1469, -1396, -1323, -1251, -1178, -1105, -1033, -960, -887, -815, -742, -669, -597, -524, -451, -379, -306, -160, 58, 276, 348, 494, 566, 639, 712, 784, 857, 930, 1002, 1075, 1148, 1221, 1293, 1366, 1439, 1511, 1584, 1657, 1729, 1802, 1875, 1947, 2020, 2093, 2165, 2238, 2311, 2384, 2456, 2529, 2602, 2674, 2747, 2820, 2892, 2965, 3038, 3110, 3183, 3256, 3328, 3401, 3474, 3547]<br>[5000], [-3577, -3504, -3431, -3359, -3286, -3213, -3141, -3068, -2995, -2923, -2850, -2777, -2705, -2632, -2559, -2486, -2414, -2341, -2268, -2196, -2123, -2050, -1978, -1905, -1832, -1760, -1687, -1614, -1542, -1469, -1396, -1323, -1251, -1178, -1033, -960, -887, -597, -451, -379, -88, -15, 58, 203, 348, 566, 712, 857, 930, 1002, 1075, 1221, 1293, 1366, 1439, 1584, 1657, 1729, 1802, 1875, 1947, 2020, 2093, 2165, 2238, 2311, 2383, 2456, 2529, 2602, 2674, 2747, 2820, 2892, 2965, 3038, 3110, 3183, 3256, 3328, 3401, 3474, 3546] |
| GAC, 20ºC<br>$T_{EX}$ = 100 ms<br>600 MHz | [80], [-3573, -3500, -3427, -3355, -3282, -3209, -3137, -3064, -2991, -2919, -2846, -2773, -2700, -2628, -2555, -2482, -2410, -2337, -2264, -2192, -2119, -2046, -1974, -1901, -1828, -1756, -1683, -1610, -1538, -1465, -1392, -1319, -1247, -1174, -1101, -1029, -956, -883, -811, -738, -665, -593, -520, -447, -375, -302, -229, -156, -84, -11, 62, 134, 207, 280, 352, 425, 498, 570, 643, 716, 788, 861, 934, 1007, 1079, 1152, 1225, 1297, 1370, 1443, 1515, 1588, 1661, 1733, 1806, 1879, 1951, 2024, 2097, 2169, 2242, 2315, 2388, 2460, 2533, 2606, 2678, 2751, 2824, 2896, 2969, 3042, 3114, 3187, 3260, 3332, 3405, 3478, 3551]<br>[160], [-3573, -3500, -3427, -3355, -3282, -3209, -3137, -3064, -2991, -2919, -2846, -2773, -2701, -2628, -2555, -2482, -2410, -2337, -2264, -2192, -2119, -2046, -1974, -1901, -1828, -1756, -1683, -1610, -1538, -1465, -1392, -1319, -1247, -1174, -1101, -1029, -956, -883, -811, -738, -665, -593, -520, -447, -375, -302, -229, -157, -84, -11, 62, 134, 207, 280, 352, 425, 498, 570, 643, 716, 788, 861, 934, 1006, 1079, 1152, 1225, 1297, 1370, 1443, 1515, 1588, 1661, 1733, 1806, 1879, 1951, 2024, 2097, 2169, 2242, 2315, 2388, 2460, 2533, 2606, 2678, 2751, 2824, 2896, 2969, 3042, 3114, 3187, 3260, 3332, 3405, 3478, 3550]<br>[320], [-3573, -3500, -3427, -3355, -3282, -3209, -3137, -3064, -2991, -2919, -2846, -2773, -2701, -2628, -2555, -2482, -2410, -2337, -2264, -2192, -2119, -2046, -1974, -1901, -1828, -1756, -1683, -1610, -1538, -1465, -1392, -1319, -1247, -1174, -1101, -1029, -956, -883, -811, -738, -665, -593, -520, -447, -375, -302, -229, -156, -84, -11, 62, 134, 207, 280, 352, 425, 498, 570, 643, 716, 788, 861, 934, 1006, 1079, 1152, 1225, 1297, 1370, 1443, 1515, 1588, 1661, 1733, 1806, 1879, 1951, 2024, 2097, 2169, 2242, 2315, 2388, 2460, 2533, 2606, 2678, 2751, 2824, 2896, 2969, 3042, 3114, 3187, 3260, 3332, 3405, 3478, 3550]<br>[640], [-3573, -3500, -3427, -3355, -3282, -3209, -3136, -3064, -2991, -2918, -2846, -2773, -2700, -2628, -2555, -2482, -2410, -2337, -2264, -2192, -2119, -2046, -1974, -1901, -1828, -1755, -1683, -1610, -1537, -1465, -1392, -1319, -1247, -1174, -1101, -1029, -956, -883, -811, -738, -665, -592, -520, -447, -374, -302, -229, -156, -84, -11, 207, 280, 352, 425, 498, 571, 643, 716, 789, 861, 934, 1007, 1079, 1152, 1225, 1297, 1370, 1443, 1515, 1588, 1661, 1733, 1806, |

| | |
|---|---|
| | 1879, 1952, 2024, 2097, 2170, 2242, 2315, 2388, 2460, 2533, 2606, 2678, 2751, 2824, 2896, 2969, 3042, 3115, 3187, 3260, 3333, 3405, 3478, 3551]<br>[1280], [-3573, -3500, -3427, -3355, -3282, -3209, -3137, -3064, -2991, -2919, -2846, -2773, -2700, -2628, -2555, -2482, -2410, -2337, -2264, -2192, -2119, -2046, -1974, -1901, -1828, -1756, -1683, -1610, -1537, -1465, -1392, -1319, -1247, -1174, -1101, -1029, -956, -883, -811, -738, -665, -593, -520, -447, -375, -302, -229, -156, -84, -11, 134, 207, 352, 425, 498, 570, 643, 716, 788, 861, 934, 1007, 1079, 1152, 1225, 1297, 1370, 1443, 1515, 1588, 1661, 1733, 1806, 1879, 1951, 2024, 2097, 2169, 2242, 2315, 2388, 2460, 2533, 2606, 2678, 2751, 2824, 2896, 2969, 3042, 3114, 3187, 3260, 3332, 3405, 3478, 3551]<br>[2560], [-3573, -3500, -3427, -3355, -3282, -3209, -3137, -3064, -2991, -2919, -2846, -2773, -2701, -2628, -2555, -2482, -2410, -2337, -2264, -2192, -2119, -2046, -1974, -1901, -1828, -1756, -1683, -1610, -1538, -1465, -1392, -1320, -1247, -1174, -1101, -1029, -956, -883, -811, -738, -665, -593, -520, -447, -302, -157, -11, 134, 280, 352, 498, 570, 643, 716, 788, 861, 934, 1006, 1079, 1152, 1224, 1297, 1370, 1443, 1515, 1588, 1661, 1733, 1806, 1879, 1951, 2024, 2097, 2169, 2242, 2315, 2387, 2460, 2533, 2606, 2678, 2751, 2824, 2896, 2969, 3042, 3114, 3187, 3260, 3332, 3405, 3478, 3550]<br>[5120], [-3573, -3500, -3427, -3355, -3282, -3209, -3137, -3064, -2991, -2919, -2846, -2773, -2700, -2628, -2555, -2482, -2410, -2337, -2264, -2192, -2119, -2046, -1974, -1901, -1828, -1756, -1683, -1610, -1537, -1465, -1392, -1319, -1247, -1174, -1101, -1029, -883, -811, -738, -593, -520, -447, -375, -229, -156, 280, 352, 425, 498, 570, 716, 788, 861, 934, 1007, 1079, 1152, 1225, 1297, 1370, 1443, 1515, 1588, 1661, 1733, 1806, 1879, 1951, 2024, 2097, 2170, 2242, 2315, 2388, 2460, 2533, 2606, 2678, 2751, 2824, 2896, 2969, 3042, 3114, 3187, 3260, 3332, 3405, 3478, 3551] |
| GAC, 15ºC<br>$T_{EX}$ = 100 ms<br>600 MHz | [20], [-3590, -3518, -3445, -3372, -3300, -3227, -3154, -3082, -3009, -2936, -2864, -2791, -2718, -2645, -2573, -2500, -2427, -2355, -2282, -2209, -2137, -2064, -1991, -1919, -1846, -1773, -1701, -1628, -1555, -1482, -1410, -1337, -1264, -1192, -1119, -1046, -974, -901, -828, -756, -683, -610, -538, -465, -392, -319, -247, -174, -101, -29, 44, 117, 189, 262, 335, 407, 480, 553, 625, 698, 771, 843, 916, 989, 1062, 1134, 1207, 1280, 1352, 1425, 1498, 1570, 1643, 1716, 1788, 1861, 1934, 2006, 2079, 2152, 2225, 2297, 2370, 2443, 2515, 2588, 2661, 2733, 2806, 2879, 2951, 3024, 3097, 3169, 3242, 3315, 3387, 3460, 3533]<br>[40], [-3590, -3518, -3445, -3372, -3300, -3227, -3154, -3082, -3009, -2936, -2863, -2791, -2718, -2645, -2573, -2500, -2427, -2355, -2282, -2209, -2137, -2064, -1991, -1919, -1846, -1773, -1700, -1628, -1555, -1482, -1410, -1337, -1264, -1192, -1119, -1046, -974, -901, -828, -756, -683, -610, -538, -465, -392, -319, -247, -174, -101, -29, 44, 117, 189, 262, 335, 407, 480, 553, 625, 698, 771, 844, 916, 989, 1062, 1134, 1207, 1280, 1352, 1425, 1498, 1570, 1643, 1716, 1788, 1861, 1934, 2007, 2079, 2152, 2225, 2297, 2370, 2443, 2515, 2588, 2661, 2733, 2806, 2879, 2951, 3024, 3097, 3169, 3242, 3315, 3388, 3460, 3533] |

| | |
|---|---|
| | [80], [-3590, -3518, -3445, -3372, -3300, -3227, -3154, -3081, -3009, -2936, -2863, -2791, -2718, -2645, -2573, -2500, -2427, -2355, -2282, -2209, -2137, -2064, -1991, -1918, -1846, -1773, -1700, -1628, -1555, -1482, -1410, -1337, -1264, -1192, -1119, -1046, -974, -901, -828, -756, -683, -610, -537, -465, -392, -319, -247, -174, -101, 44, 117, 189, 262, 335, 407, 480, 553, 626, 698, 771, 844, 916, 989, 1062, 1134, 1207, 1280, 1352, 1425, 1498, 1570, 1643, 1716, 1789, 1861, 1934, 2007, 2079, 2152, 2225, 2297, 2370, 2443, 2515, 2588, 2661, 2733, 2806, 2879, 2951, 3024, 3097, 3170, 3242, 3315, 3388, 3460, 3533]<br>[160], [-3590, -3518, -3445, -3372, -3300, -3227, -3154, -3081, -3009, -2936, -2863, -2791, -2718, -2645, -2573, -2500, -2427, -2355, -2282, -2209, -2137, -2064, -1991, -1919, -1846, -1773, -1700, -1628, -1555, -1482, -1410, -1337, -1264, -1192, -1119, -1046, -974, -901, -828, -756, -683, -610, -537, -465, -392, -319, -247, -174, -101, 44, 117, 189, 262, 335, 407, 480, 553, 625, 698, 771, 844, 916, 989, 1062, 1134, 1207, 1280, 1352, 1425, 1498, 1570, 1643, 1716, 1788, 1861, 1934, 2007, 2079, 2152, 2225, 2297, 2370, 2443, 2515, 2588, 2661, 2733, 2806, 2879, 2951, 3024, 3097, 3170, 3242, 3315, 3388, 3460, 3533]<br>[320], [-3590, -3518, -3445, -3372, -3300, -3227, -3154, -3082, -3009, -2936, -2863, -2791, -2718, -2645, -2573, -2500, -2427, -2355, -2282, -2209, -2137, -2064, -1991, -1919, -1846, -1773, -1701, -1628, -1555, -1482, -1410, -1337, -1264, -1192, -1119, -1046, -974, -901, -828, -756, -683, -610, -538, -465, -392, -319, -247, -174, -29, 44, 117, 189, 262, 335, 407, 480, 553, 625, 698, 771, 844, 916, 989, 1062, 1134, 1207, 1280, 1352, 1425, 1498, 1570, 1643, 1716, 1788, 1861, 1934, 2006, 2079, 2152, 2225, 2297, 2370, 2443, 2515, 2588, 2661, 2733, 2806, 2879, 2951, 3024, 3097, 3169, 3242, 3315, 3388, 3460, 3533]<br>[640], [-3590, -3518, -3445, -3372, -3300, -3227, -3154, -3081, -3009, -2936, -2863, -2791, -2718, -2645, -2573, -2500, -2427, -2355, -2282, -2209, -2137, -2064, -1991, -1918, -1846, -1773, -1700, -1628, -1555, -1482, -1410, -1337, -1264, -1192, -1119, -1046, -974, -901, -828, -756, -683, -610, -537, -465, -392, -319, -247, -174, -101, -29, 117, 189, 262, 335, 407, 480, 553, 626, 698, 771, 844, 916, 989, 1062, 1134, 1207, 1280, 1352, 1425, 1498, 1570, 1643, 1716, 1789, 1861, 1934, 2007, 2079, 2152, 2225, 2297, 2370, 2443, 2515, 2588, 2661, 2733, 2806, 2879, 2951, 3024, 3097, 3170, 3242, 3315, 3388, 3460, 3533]<br>[1280], [-3590, -3518, -3445, -3372, -3300, -3227, -3154, -3082, -3009, -2936, -2863, -2791, -2718, -2645, -2573, -2500, -2427, -2355, -2282, -2209, -2137, -2064, -1991, -1919, -1846, -1773, -1700, -1628, -1555, -1482, -1410, -1337, -1264, -1192, -1119, -1046, -974, -901, -828, -756, -683, -610, -537, -465, -392, -319, 117, 189, 262, 335, 407, 480, 553, 625, 698, 771, 844, 916, 989, 1062, 1134, 1207, 1280, 1352, 1425, 1498, 1570, 1643, 1716, 1788, 1861, 1934, 2007, 2079, 2152, 2225, 2297, 2370, 2443, 2515, 2588, 2661, 2733, 2806, 2879, 2951, 3024, 3097, 3169, 3242, 3315, 3388, 3460, 3533] |

| | |
|---|---|
| | [2560], [-3590, -3518, -3445, -3372, -3300, -3227, -3154, -3082, -3009, -2936, -2864, -2791, -2718, -2645, -2573, -2500, -2427, -2355, -2282, -2209, -2137, -2064, -1991, -1919, -1846, -1773, -1701, -1628, -1555, -1482, -1410, -1337, -1264, -1192, -1119, -1046, -974, -901, -828, -756, -683, -610, -538, -465, -392, -319, -247, -174, -29, 44, 117, 335, 407, 480, 553, 625, 698, 771, 843, 916, 989, 1062, 1134, 1207, 1280, 1352, 1425, 1498, 1570, 1643, 1716, 1788, 1861, 1934, 2006, 2079, 2152, 2225, 2297, 2370, 2443, 2515, 2588, 2661, 2733, 2806, 2879, 2951, 3024, 3097, 3169, 3242, 3315, 3387, 3460, 3533] |
| GAC, 10ºC<br>$T_{EX}$ = 100 ms<br>600 MHz | [20], [-3564, -3491, -3419, -3346, -3273, -3201, -3128, -3055, -2983, -2910, -2837, -2765, -2692, -2619, -2547, -2474, -2401, -2329, -2256, -2183, -2110, -2038, -1965, -1892, -1820, -1747, -1674, -1602, -1529, -1456, -1384, -1311, -1238, -1166, -1093, -1020, -947, -875, -802, -729, -657, -584, -511, -439, -366, -293, -221, -148, -75, -3, 70, 143, 215, 288, 361, 434, 506, 579, 652, 724, 797, 870, 942, 1015, 1088, 1160, 1233, 1306, 1378, 1451, 1524, 1597, 1669, 1742, 1815, 1887, 1960, 2033, 2105, 2178, 2251, 2323, 2396, 2469, 2541, 2614, 2687, 2760, 2832, 2905, 2978, 3050, 3123, 3196, 3268, 3341, 3414, 3486, 3559]<br>[40], [-3564, -3491, -3419, -3346, -3273, -3201, -3128, -3055, -2983, -2910, -2837, -2765, -2692, -2619, -2547, -2474, -2401, -2328, -2256, -2183, -2110, -2038, -1965, -1892, -1820, -1747, -1674, -1602, -1529, -1456, -1384, -1311, -1238, -1166, -1093, -1020, -947, -875, -802, -729, -657, -584, -511, -439, -366, -293, -221, -148, -75, -3, 70, 143, 216, 288, 361, 434, 506, 579, 652, 724, 797, 870, 942, 1015, 1088, 1160, 1233, 1306, 1379, 1451, 1524, 1597, 1669, 1742, 1815, 1887, 1960, 2033, 2105, 2178, 2251, 2323, 2396, 2469, 2541, 2614, 2687, 2760, 2832, 2905, 2978, 3050, 3123, 3196, 3268, 3341, 3414, 3486, 3559]<br>[80], [-3564, -3492, -3419, -3346, -3273, -3201, -3128, -3055, -2983, -2910, -2837, -2765, -2692, -2619, -2547, -2474, -2401, -2329, -2256, -2183, -2111, -2038, -1965, -1892, -1820, -1747, -1674, -1602, -1529, -1456, -1384, -1311, -1238, -1166, -1093, -1020, -948, -875, -802, -729, -657, -584, -511, -439, -366, -293, -221, -148, -75, 70, 143, 215, 288, 361, 433, 506, 579, 652, 724, 797, 870, 942, 1015, 1088, 1160, 1233, 1306, 1378, 1451, 1524, 1596, 1669, 1742, 1815, 1887, 1960, 2033, 2105, 2178, 2251, 2323, 2396, 2469, 2541, 2614, 2687, 2759, 2832, 2905, 2978, 3050, 3123, 3196, 3268, 3341, 3414, 3486, 3559]<br>[160], [-3564, -3492, -3419, -3346, -3273, -3201, -3128, -3055, -2983, -2910, -2837, -2765, -2692, -2619, -2547, -2474, -2401, -2329, -2256, -2183, -2110, -2038, -1965, -1892, -1820, -1747, -1674, -1602, -1529, -1456, -1384, -1311, -1238, -1166, -1093, -1020, -948, -875, -802, -729, -657, -584, -511, -439, -366, -293, -221, -148, -75, 70, 143, 215, 288, 361, 434, 506, 579, 652, 724, 797, 870, 942, 1015, 1088, 1160, 1233, 1306, 1378, 1451, 1524, 1596, 1669, 1742, 1815, 1887, 1960, 2033, 2105, 2178, 2251, 2323, 2396, 2469, 2541, 2614, 2687, 2759, 2832, 2905, 2978, 3050, 3123, 3196, 3268, 3341, 3414, 3486, 3559]<br>[320], [-3564, -3491, -3419, -3346, -3273, -3201, -3128, -3055, -2983, -2910, -2837, -2765, -2692, -2619, -2547, -2474, -2401, -2328, -2256, -2183, -2110, - |

| | |
|---|---|
| | 2038, -1965, -1892, -1820, -1747, -1674, -1602, -1529, -1456, -1384, -1311, -1238, -1165, -1093, -1020, -947, -875, -802, -729, -657, -584, -511, -439, -366, -293, -221, -75, 70, 143, 216, 288, 361, 434, 506, 579, 652, 724, 797, 870, 942, 1015, 1088, 1160, 1233, 1306, 1379, 1451, 1524, 1597, 1669, 1742, 1815, 1887, 1960, 2033, 2105, 2178, 2251, 2323, 2396, 2469, 2542, 2614, 2687, 2760, 2832, 2905, 2978, 3050, 3123, 3196, 3268, 3341, 3414, 3486, 3559]<br>[640], [-3564, -3491, -3419, -3346, -3273, -3201, -3128, -3055, -2983, -2910, -2837, -2765, -2692, -2619, -2546, -2474, -2401, -2328, -2256, -2183, -2110, -2038, -1965, -1892, -1820, -1747, -1674, -1602, -1529, -1456, -1384, -1311, -1238, -1165, -1093, -1020, -947, -875, -802, -729, -657, -584, -511, -439, -366, -293, -221, -148, -75, 143, 216, 288, 361, 434, 506, 579, 652, 724, 797, 870, 942, 1015, 1088, 1160, 1233, 1306, 1379, 1451, 1524, 1597, 1669, 1742, 1815, 1887, 1960, 2033, 2105, 2178, 2251, 2323, 2396, 2469, 2542, 2614, 2687, 2760, 2832, 2905, 2978, 3050, 3123, 3196, 3268, 3341, 3414, 3486, 3559] |
| CAC, 25ºC<br>$T_{EX}$ = 100 ms<br>900 MHz | [100], [-5287, -5042, -4797, -4551, -4306, -4060, -3815, -3569, -3324, -3078, -2833, -2735, -2636, -2538, -2440, -2342, -2244, -2146, -2047, -1949, -1851, -1753, -1655, -1556, -1458, -1360, -1262, -1164, -1065, -967, -869, -771, -673, -575, -476, -378, -280, -182, -84, 113, 211, 309, 407, 505, 604, 702, 800, 898, 996, 1095, 1193, 1291, 1389, 1487, 1586, 1684, 1782, 1880, 1978, 2076, 2175, 2273, 2371, 2469, 2567, 2813, 3058, 3304, 3549, 3795, 4040, 4286, 4531, 4777, 5022, 5267]<br>[250], [-5287, -5042, -4797, -4551, -4306, -4060, -3815, -3569, -3324, -3078, -2833, -2735, -2636, -2538, -2440, -2342, -2244, -2146, -2047, -1949, -1851, -1753, -1655, -1556, -1458, -1360, -1262, -1164, -1065, -967, -869, -771, -673, -575, -476, -378, -280, -182, -84, 113, 211, 309, 407, 505, 604, 702, 800, 898, 996, 1095, 1193, 1291, 1389, 1487, 1586, 1684, 1782, 1880, 1978, 2076, 2175, 2273, 2371, 2469, 2567, 2813, 3058, 3304, 3549, 3795, 4040, 4286, 4531, 4777, 5022, 5267]<br>[500], [-5288, -5042, -4797, -4551, -4306, -4060, -3815, -3569, -3324, -3079, -2833, -2735, -2637, -2539, -2440, -2342, -2244, -2146, -2048, -1949, -1851, -1753, -1655, -1557, -1458, -1360, -1262, -1164, -1066, -968, -869, -771, -673, -575, -477, -378, -280, -182, -84, 14, 112, 211, 309, 407, 505, 603, 702, 800, 898, 996, 1094, 1193, 1291, 1389, 1487, 1585, 1683, 1782, 1880, 1978, 2076, 2174, 2273, 2371, 2469, 2567, 2813, 3058, 3303, 3549, 3794, 4040, 4285, 4531, 4776, 5022, 5267]<br>[1000], [-5287, -5042, -4796, -4551, -4305, -4060, -3814, -3569, -3323, -3078, -2833, -2734, -2636, -2538, -2440, -2342, -2243, -2145, -2047, -1949, -1851, -1752, -1654, -1556, -1458, -1360, -1262, -1163, -1065, -967, -869, -771, -672, -574, -476, -378, -280, -182, -83, 15, 113, 211, 309, 408, 506, 604, 702, 800, 899, 997, 1095, 1193, 1291, 1389, 1488, 1586, 1684, 1782, 1880, 1979, 2077, 2175, 2273, 2371, 2469, 2568, 2813, 3059, 3304, 3550, 3795, 4040, 4286, 4531, 4777, 5022, 5268]<br>[2000], [-5287, -5042, -4796, -4551, -4305, -4060, -3814, -3569, -3324, -3078, -2833, -2734, -2636, -2538, -2440, -2342, -2243, -2145, -2047, -1949, -1851, - |

| | |
|---|---|
| | 1753, -1654, -1556, -1458, -1360, -1262, -1163, -1065, -967, -869, -771, -673, -574, -378, -280, -83, 113, 309, 408, 506, 604, 702, 800, 898, 997, 1095, 1193, 1291, 1389, 1488, 1586, 1684, 1782, 1880, 1978, 2077, 2175, 2273, 2371, 2469, 2568, 2813, 3059, 3304, 3549, 3795, 4040, 4286, 4531, 4777, 5022, 5268]<br>[4000], [-5288, -5042, -4797, -4551, -4306, -4060, -3815, -3569, -3324, -3078, -2833, -2735, -2637, -2538, -2440, -2342, -2244, -2146, -2048, -1949, -1851, -1753, -1655, -1557, -1458, -1360, -1262, -1164, -1066, -967, -869, -771, -673, -575, -477, -280, -182, -84, 14, 113, 211, 309, 407, 603, 702, 800, 898, 996, 1094, 1193, 1291, 1389, 1487, 1585, 1684, 1782, 1880, 1978, 2076, 2174, 2273, 2371, 2469, 2567, 2813, 3058, 3304, 3549, 3794, 4040, 4285, 4531, 4776, 5022, 5267] |
| CAC, 20ºC<br>$T_{EX}$ = 100 ms<br>900 MHz | [100], [-5259, -4959, -4659, -4359, -4059, -3759, -3459, -3159, -2859, -2759, -2659, -2559, -2459, -2359, -2259, -2159, -2059, -1959, -1859, -1759, -1659, -1559, -1459, -1359, -1259, -1159, -1059, -959, -859, -759, -659, -559, -459, -359, -259, -159, -59, 141, 241, 341, 441, 541, 641, 741, 841, 941, 1041, 1141, 1241, 1341, 1441, 1541, 1641, 1741, 1841, 1941, 2041, 2141, 2241, 2341, 2441, 2541, 2841, 3141, 3441, 3741, 4041, 4341, 4641, 4941, 5241]<br>[500], [-5260, -4960, -4660, -4360, -4060, -3760, -3460, -3160, -2860, -2760, -2660, -2560, -2460, -2360, -2260, -2160, -2060, -1960, -1860, -1760, -1660, -1560, -1460, -1360, -1260, -1160, -1060, -960, -860, -760, -660, -560, -460, -360, -260, -160, -60, 40, 140, 240, 340, 440, 540, 640, 740, 840, 940, 1040, 1140, 1240, 1340, 1440, 1540, 1640, 1740, 1840, 1940, 2040, 2140, 2240, 2340, 2440, 2540, 2840, 3140, 3440, 3740, 4040, 4340, 4640, 4940, 5240]<br>[1000], [-5260, -4960, -4660, -4360, -4060, -3760, -3460, -3160, -2860, -2760, -2660, -2559, -2459, -2359, -2259, -2159, -2059, -1959, -1859, -1759, -1659, -1559, -1459, -1359, -1259, -1159, -1059, -959, -859, -759, -659, -559, -459, -359, -259, 41, 141, 341, 441, 541, 641, 741, 841, 941, 1041, 1141, 1241, 1341, 1441, 1541, 1641, 1741, 1841, 1941, 2041, 2141, 2241, 2341, 2441, 2541, 2841, 3141, 3441, 3741, 4041, 4341, 4641, 4941, 5241]<br>[1500], [-5260, -4960, -4660, -4360, -4060, -3760, -3460, -3160, -2860, -2760, -2660, -2560, -2460, -2360, -2260, -2160, -2060, -1960, -1860, -1760, -1660, -1560, -1460, -1360, -1260, -1160, -1060, -960, -860, -760, -660, -560, -460, -360, -260, -160, 40, 240, 340, 440, 540, 640, 740, 840, 940, 1040, 1140, 1240, 1340, 1440, 1540, 1640, 1740, 1840, 1940, 2040, 2140, 2240, 2340, 2440, 2540, 2840, 3140, 3440, 3740, 4040, 4340, 4640, 4940, 5240]<br>[2500], [-5260, -4960, -4660, -4360, -4060, -3760, -3460, -3160, -2860, -2760, -2660, -2560, -2460, -2360, -2260, -2160, -2060, -1960, -1860, -1760, -1660, -1560, -1460, -1360, -1260, -1160, -1060, -960, -860, -760, -660, -560, -460, -360, -260, -160, 241, 341, 441, 541, 641, 741, 841, 941, 1041, 1141, 1241, 1341, 1441, 1541, 1641, 1741, 1841, 1941, 2041, 2141, 2241, 2341, 2441, 2541, 2841, 3141, 3441, 3741, 4041, 4341, 4641, 4941, 5241]<br>[4000], [-5259, -4959, -4659, -4359, -4059, -3759, -3459, -3159, -2859, -2759, -2659, -2559, -2459, -2359, -2259, -2159, -2059, -1959, -1859, -1759, -1659, - |

| | |
|---|---|
| | 1559, -1459, -1359, -1259, -1159, -1059, -959, -859, -759, -659, -559, -459, -259, 41, 441, 541, 641, 741, 841, 941, 1041, 1141, 1241, 1341, 1441, 1541, 1641, 1741, 1841, 1941, 2041, 2141, 2241, 2341, 2441, 2541, 2841, 3141, 3441, 3741, 4041, 4341, 4641, 4941, 5241] |
| CAC, 15ºC<br>$T_{EX}$ = 100 ms<br>900 MHz | [100], [-5394, -5201, -5009, -4816, -4623, -4430, -4237, -4044, -3851, -3659, -3466, -3273, -3080, -2887, -2796, -2704, -2612, -2521, -2429, -2338, -2246, -2155, -2063, -1972, -1880, -1789, -1697, -1606, -1514, -1423, -1331, -1240, -1148, -1056, -965, -873, -782, -690, -599, -507, -416, -324, -233, -141, -50, 133, 225, 316, 408, 499, 591, 683, 774, 866, 957, 1049, 1140, 1232, 1323, 1415, 1506, 1598, 1689, 1781, 1872, 1964, 2055, 2147, 2239, 2330, 2422, 2513, 2706, 2899, 3092, 3285, 3477, 3670, 3863, 4056, 4249, 4442, 4635, 4827, 5020, 5213]<br>[250], [-5395, -5202, -5009, -4816, -4623, -4430, -4237, -4045, -3852, -3659, -3466, -3273, -3080, -2887, -2796, -2704, -2613, -2521, -2430, -2338, -2247, -2155, -2064, -1972, -1881, -1789, -1698, -1606, -1514, -1423, -1331, -1240, -1148, -1057, -965, -874, -782, -691, -599, -508, -416, -325, -233, -142, -50, 42, 133, 225, 316, 408, 499, 591, 682, 774, 865, 957, 1048, 1140, 1231, 1323, 1414, 1506, 1597, 1689, 1781, 1872, 1964, 2055, 2147, 2238, 2330, 2421, 2513, 2706, 2899, 3091, 3284, 3477, 3670, 3863, 4056, 4249, 4441, 4634, 4827, 5020, 5213]<br>[500], [-5395, -5202, -5009, -4816, -4623, -4430, -4238, -4045, -3852, -3659, -3466, -3273, -3080, -2888, -2796, -2705, -2613, -2521, -2430, -2338, -2247, -2155, -2064, -1972, -1881, -1789, -1698, -1606, -1515, -1423, -1332, -1240, -1149, -1057, -965, -874, -782, -691, -599, -508, -416, -325, -233, -142, -50, 133, 224, 316, 407, 499, 591, 682, 774, 865, 957, 1048, 1140, 1231, 1323, 1414, 1506, 1597, 1689, 1780, 1872, 1963, 2055, 2147, 2238, 2330, 2421, 2513, 2705, 2898, 3091, 3284, 3477, 3670, 3863, 4056, 4248, 4441, 4634, 4827, 5020, 5213]<br>[1000], [-5394, -5201, -5009, -4816, -4623, -4430, -4237, -4044, -3851, -3659, -3466, -3273, -3080, -2887, -2796, -2704, -2613, -2521, -2429, -2338, -2246, -2155, -2063, -1972, -1880, -1789, -1697, -1606, -1514, -1423, -1331, -1240, -1148, -1057, -965, -873, -782, -690, -599, -507, -416, -324, -233, -141, 42, 133, 225, 316, 408, 499, 591, 683, 774, 866, 957, 1049, 1140, 1232, 1323, 1415, 1506, 1598, 1689, 1781, 1872, 1964, 2055, 2147, 2239, 2330, 2422, 2513, 2706, 2899, 3092, 3285, 3477, 3670, 3863, 4056, 4249, 4442, 4635, 4827, 5020, 5213]<br>[1500], [-5395, -5202, -5009, -4816, -4623, -4430, -4237, -4045, -3852, -3659, -3466, -3273, -3080, -2887, -2796, -2704, -2613, -2521, -2430, -2338, -2247, -2155, -2064, -1972, -1881, -1789, -1697, -1606, -1514, -1423, -1331, -1240, -1148, -1057, -965, -874, -782, -691, -599, -508, -416, -325, -233, -141, -50, 42, 133, 225, 316, 408, 499, 591, 682, 774, 865, 957, 1048, 1140, 1231, 1323, 1414, 1506, 1598, 1689, 1781, 1872, 1964, 2055, 2147, 2238, 2330, 2421, 2513, 2706, 2899, 3091, 3284, 3477, 3670, 3863, 4056, 4249, 4441, 4634, 4827, 5020, 5213] |

| | |
|---|---|
| | [2000], [-5394, -5201, -5009, -4816, -4623, -4430, -4237, -4044, -3851, -3659, -3466, -3273, -3080, -2887, -2796, -2704, -2613, -2521, -2429, -2338, -2246, -2155, -2063, -1972, -1880, -1789, -1697, -1606, -1514, -1423, -1331, -1240, -1148, -1057, -965, -873, -782, -690, -599, -507, -416, -324, -141, -50, 42, 133, 225, 316, 408, 499, 591, 683, 774, 866, 957, 1049, 1140, 1232, 1323, 1415, 1506, 1598, 1689, 1781, 1872, 1964, 2055, 2147, 2238, 2330, 2422, 2513, 2706, 2899, 3092, 3285, 3477, 3670, 3863, 4056, 4249, 4442, 4635, 4827, 5020, 5213] |
| CAC, 7ºC<br>$T_{EX}$ = 100 ms<br>900 MHz | [25], [-5299, -5053, -4808, -4562, -4317, -4071, -3826, -3581, -3335, -3090, -2844, -2746, -2648, -2550, -2451, -2353, -2255, -2157, -2059, -1960, -1862, -1764, -1666, -1568, -1470, -1371, -1273, -1175, -1077, -979, -880, -782, -684, -586, -488, -389, -291, -193, -95, 3, 101, 200, 298, 396, 494, 592, 691, 789, 887, 985, 1083, 1181, 1280, 1378, 1476, 1574, 1672, 1771, 1869, 1967, 2065, 2163, 2262, 2360, 2458, 2556, 2802, 3047, 3292, 3538, 3783, 4029, 4274, 4520, 4765, 5011, 5256]<br>[100], [-5298, -5053, -4807, -4562, -4316, -4071, -3825, -3580, -3334, -3089, -2844, -2745, -2647, -2549, -2451, -2353, -2254, -2156, -2058, -1960, -1862, -1764, -1665, -1567, -1469, -1371, -1273, -1174, -1076, -978, -880, -782, -683, -585, -487, -389, -291, -193, -94, 4, 102, 200, 298, 397, 495, 593, 691, 789, 887, 986, 1084, 1182, 1280, 1378, 1477, 1575, 1673, 1771, 1869, 1968, 2066, 2164, 2262, 2360, 2458, 2557, 2802, 3048, 3293, 3538, 3784, 4029, 4275, 4520, 4766, 5011, 5257]<br>[400], [-5299, -5053, -4808, -4562, -4317, -4071, -3826, -3581, -3335, -3090, -2844, -2746, -2648, -2550, -2451, -2353, -2255, -2157, -2059, -1960, -1862, -1764, -1666, -1568, -1470, -1371, -1273, -1175, -1077, -979, -880, -782, -684, -586, -488, -389, -291, -193, 101, 200, 298, 396, 494, 592, 691, 789, 887, 985, 1083, 1181, 1280, 1378, 1476, 1574, 1672, 1771, 1869, 1967, 2065, 2163, 2262, 2360, 2458, 2556, 2802, 3047, 3292, 3538, 3783, 4029, 4274, 4520, 4765, 5011, 5256]<br>[800], [-5299, -5053, -4808, -4562, -4317, -4071, -3826, -3581, -3335, -3090, -2844, -2746, -2648, -2550, -2451, -2353, -2255, -2157, -2059, -1960, -1862, -1764, -1666, -1568, -1470, -1371, -1273, -1175, -1077, -979, -880, -782, -684, -586, -488, -389, -291, -193, 3, 200, 298, 396, 494, 592, 691, 789, 887, 985, 1083, 1181, 1280, 1378, 1476, 1574, 1672, 1771, 1869, 1967, 2065, 2163, 2262, 2360, 2458, 2556, 2802, 3047, 3292, 3538, 3783, 4029, 4274, 4520, 4765, 5011, 5256]<br>[1200], [-5298, -5053, -4807, -4562, -4317, -4071, -3826, -3580, -3335, -3089, -2844, -2746, -2647, -2549, -2451, -2353, -2255, -2156, -2058, -1960, -1862, -1764, -1666, -1567, -1469, -1371, -1273, -1175, -1076, -978, -880, -782, -684, -585, -487, -389, -291, -193, 102, 200, 298, 396, 495, 593, 691, 789, 887, 985, 1084, 1182, 1280, 1378, 1476, 1575, 1673, 1771, 1869, 1967, 2066, 2164, 2262, 2360, 2458, 2556, 2802, 3047, 3293, 3538, 3784, 4029, 4275, 4520, 4766, 5011, 5257] |

| | |
|---|---|
| | [1500], [-5298, -5053, -4807, -4562, -4316, -4071, -3825, -3580, -3335, -3089, -2844, -2745, -2647, -2549, -2451, -2353, -2255, -2156, -2058, -1960, -1862, -1764, -1665, -1567, -1469, -1371, -1273, -1174, -1076, -978, -880, -782, -684, -585, -487, -389, -291, 102, 298, 396, 495, 593, 691, 789, 887, 986, 1084, 1182, 1280, 1378, 1477, 1575, 1673, 1771, 1869, 1967, 2066, 2164, 2262, 2360, 2458, 2557, 2802, 3047, 3293, 3538, 3784, 4029, 4275, 4520, 4766, 5011, 5257] |
| hpGAG, 25ºC $T_{EX}$ = 100 ms 600 MHz | [30], [-3216, -3143, -3070, -2997, -2925, -2852, -2779, -2707, -2634, -2561, -2489, -2416, -2343, -2271, -2198, -2125, -2053, -1980, -1907, -1834, -1762, -1689, -1616, -1544, -1471, -1398, -1326, -1253, -1180, -1108, -1035, -962, -890, -817, -744, -672, -599, -526, -453, -381, -308, -235, -163, -90, -17, 55, 128, 201, 273, 346, 419, 491, 564, 637, 710, 782, 855, 928, 1000, 1073, 1146, 1218, 1291, 1364, 1436, 1509, 1582, 1654, 1727, 1800, 1872, 1945, 2018, 2091, 2163, 2236, 2309, 2381, 2454, 2527, 2599, 2672, 2745, 2817, 2890, 2963, 3035, 3108, 3181, 3254, 3326, 3399, 3472, 3544]<br>[90], [-3216, -3143, -3070, -2997, -2925, -2852, -2779, -2707, -2634, -2561, -2489, -2416, -2343, -2271, -2198, -2125, -2053, -1980, -1907, -1834, -1762, -1689, -1616, -1544, -1471, -1398, -1326, -1253, -1180, -1108, -1035, -962, -890, -817, -744, -672, -599, -526, -453, -381, -308, -235, -163, -90, -17, 55, 128, 201, 273, 346, 419, 491, 564, 637, 710, 782, 855, 928, 1000, 1073, 1146, 1218, 1291, 1364, 1436, 1509, 1582, 1654, 1727, 1800, 1873, 1945, 2018, 2091, 2163, 2236, 2309, 2381, 2454, 2527, 2599, 2672, 2745, 2817, 2890, 2963, 3035, 3108, 3181, 3254, 3326, 3399, 3472, 3544]<br>[270], [-3216, -3143, -3070, -2997, -2925, -2852, -2779, -2707, -2634, -2561, -2489, -2416, -2343, -2271, -2198, -2125, -2053, -1980, -1907, -1835, -1762, -1689, -1616, -1544, -1471, -1398, -1326, -1253, -1180, -1108, -1035, -962, -890, -817, -744, -672, -599, -526, -453, -381, -308, -235, -163, -17, 128, 201, 273, 346, 419, 491, 564, 637, 710, 782, 855, 928, 1000, 1073, 1146, 1218, 1291, 1364, 1436, 1509, 1582, 1654, 1727, 1800, 1872, 1945, 2018, 2091, 2163, 2236, 2309, 2381, 2454, 2527, 2599, 2672, 2745, 2817, 2890, 2963, 3035, 3108, 3181, 3254, 3326, 3399, 3472, 3544]<br>[810], [-3215, -3143, -3070, -2997, -2925, -2852, -2779, -2707, -2634, -2561, -2489, -2416, -2343, -2271, -2198, -2125, -2053, -1980, -1907, -1834, -1762, -1689, -1616, -1544, -1471, -1398, -1326, -1253, -1180, -1108, -1035, -962, -890, -817, -744, -671, -599, -526, -453, -381, -308, -235, -163, -90, 201, 273, 346, 419, 492, 564, 637, 710, 782, 855, 928, 1000, 1073, 1146, 1218, 1291, 1364, 1436, 1509, 1582, 1654, 1727, 1800, 1873, 1945, 2018, 2091, 2163, 2236, 2309, 2381, 2454, 2527, 2599, 2672, 2745, 2817, 2890, 2963, 3036, 3108, 3181, 3254, 3326, 3399, 3472, 3544]<br>[2430], [-3216, -3143, -3070, -2997, -2925, -2852, -2779, -2707, -2634, -2561, -2489, -2416, -2343, -2271, -2198, -2125, -2053, -1980, -1907, -1834, -1762, -1689, -1616, -1544, -1471, -1398, -1326, -1253, -1180, -1108, -1035, -962, -890, -817, -744, -672, -599, -526, -453, -308, -17, 128, 273, 346, 419, 491, 564, 637, 710, 782, 855, 928, 1000, 1073, 1146, 1218, 1291, 1364, 1436, 1509, 1582, 1654, 1727, 1800, 1873, 1945, 2018, 2091, 2163, 2236, 2309, 2381, 2454, |

| | |
|---|---|
| | 2527, 2599, 2672, 2745, 2817, 2890, 2963, 3035, 3108, 3181, 3254, 3326,<br>3399, 3472, 3544]<br>[5000], [-3215, -3143, -3070, -2997, -2925, -2852, -2779, -2707, -2634, -2561, -<br>2489, -2416, -2343, -2271, -2198, -2125, -2053, -1980, -1907, -1834, -1762, -<br>1689, -1616, -1544, -1471, -1398, -1326, -1253, -1180, -1108, -1035, -962, -890,<br>-817, -744, -671, -599, -453, -381, -308, -235, -163, 128, 201, 419, 491, 564,<br>710, 782, 855, 928, 1000, 1073, 1146, 1218, 1291, 1364, 1436, 1509, 1582,<br>1654, 1727, 1800, 1873, 1945, 2018, 2091, 2163, 2236, 2309, 2381, 2454,<br>2527, 2599, 2672, 2745, 2817, 2890, 2963, 3036, 3108, 3181, 3254, 3326,<br>3399, 3472, 3544] |
| hpCAC,<br>25ºC<br>$T_{EX}$ = 100<br>ms<br>700 MHz | [100], [-3955, -3861, -3766, -3672, -3578, -3483, -3389, -3295, -3200, -3106, -<br>3012, -2917, -2823, -2728, -2634, -2540, -2445, -2351, -2257, -2162, -2068, -<br>1973, -1879, -1785, -1690, -1596, -1502, -1407, -1313, -1218, -1124, -1030, -<br>935, -841, -747, -652, -558, -463, -369, -275, -180, -86, 103, 197, 292, 386, 480,<br>575, 669, 763, 858, 952, 1047, 1141, 1235, 1330, 1424, 1518, 1613, 1707, 1802,<br>1896, 1990, 2085, 2179, 2273, 2368, 2462, 2557, 2651, 2745, 2840, 2934,<br>3028, 3123, 3217, 3311, 3406, 3500, 3595, 3689, 3783, 3878, 3972, 4066, 4161,<br>4255, 4350, 4444]<br>[250], [-3955, -3861, -3767, -3672, -3578, -3483, -3389, -3295, -3200, -3106, -<br>3012, -2917, -2823, -2728, -2634, -2540, -2445, -2351, -2257, -2162, -2068, -<br>1974, -1879, -1785, -1690, -1596, -1502, -1407, -1313, -1219, -1124, -1030, -<br>935, -841, -747, -652, -558, -464, -369, -275, -180, -86, 103, 197, 291, 386, 480,<br>575, 669, 763, 858, 952, 1046, 1141, 1235, 1330, 1424, 1518, 1613, 1707, 1801,<br>1896, 1990, 2085, 2179, 2273, 2368, 2462, 2556, 2651, 2745, 2840, 2934,<br>3028, 3123, 3217, 3311, 3406, 3500, 3595, 3689, 3783, 3878, 3972, 4066, 4161,<br>4255, 4349, 4444]<br>[600], [-3955, -3861, -3767, -3672, -3578, -3483, -3389, -3295, -3200, -3106, -<br>3012, -2917, -2823, -2728, -2634, -2540, -2445, -2351, -2257, -2162, -2068, -<br>1973, -1879, -1785, -1690, -1596, -1502, -1407, -1313, -1219, -1124, -1030, -<br>935, -841, -747, -652, -558, -464, -369, -275, -180, -86, 8, 103, 197, 291, 386,<br>480, 575, 669, 763, 858, 952, 1046, 1141, 1235, 1330, 1424, 1518, 1613, 1707,<br>1801, 1896, 1990, 2085, 2179, 2273, 2368, 2462, 2556, 2651, 2745, 2840,<br>2934, 3028, 3123, 3217, 3311, 3406, 3500, 3595, 3689, 3783, 3878, 3972, 4066,<br>4161, 4255, 4350, 4444]<br>[1000], [-3955, -3861, -3767, -3672, -3578, -3483, -3389, -3295, -3200, -3106, -<br>3012, -2917, -2823, -2728, -2634, -2540, -2445, -2351, -2257, -2162, -2068, -<br>1973, -1879, -1785, -1690, -1596, -1502, -1407, -1313, -1219, -1124, -1030, -<br>935, -841, -747, -652, -558, -464, -369, -275, -180, -86, 197, 291, 386, 480, 575,<br>669, 763, 858, 952, 1046, 1141, 1235, 1330, 1424, 1518, 1613, 1707, 1801,<br>1896, 1990, 2085, 2179, 2273, 2368, 2462, 2556, 2651, 2745, 2840, 2934,<br>3028, 3123, 3217, 3311, 3406, 3500, 3595, 3689, 3783, 3878, 3972, 4066, 4161,<br>4255, 4350, 4444] |

**Table S6.** RF powers and offsets used in [1]H CEST experiments performed at 100 mM NaCl, pH 8 and T=25ºC.

| Sample | [RF field power] [offset frequencies] |
|---|---|
| | [ω/2π (Hz)] [Ω/2π (Hz)] |
| AAA, 25ºC<br>$T_{EX}$ = 100 ms<br>600 MHz | [10], [-3154, -3082, -3009, -2936, -2864, -2791, -2718, -2645, -2573, -2500, -2427, -2355, -2282, -2209, -2137, -2064, -1991, -1919, -1846, -1773, -1701, -1628, -1555, -1482, -1410, -1337, -1264, -1192, -1119, -1046, -974, -901, -828, -756, -683, -610, -538, -465, -392, -319, -247, -174, -101, -29, 44, 117, 189, 262, 335, 407, 480, 553, 625, 698, 771, 843, 916, 989, 1062, 1134, 1207, 1280, 1352, 1425, 1498, 1570, 1643, 1716, 1788, 1861, 1934, 2006, 2079, 2152, 2225, 2297, 2370, 2443, 2515, 2588, 2661, 2733, 2806, 2879, 2951, 3024, 3097, 3169, 3242, 3315, 3387, 3460, 3533]<br>[30], [-3154, -3082, -3009, -2936, -2863, -2791, -2718, -2645, -2573, -2500, -2427, -2355, -2282, -2209, -2137, -2064, -1991, -1919, -1846, -1773, -1701, -1628, -1555, -1482, -1410, -1337, -1264, -1192, -1119, -1046, -974, -901, -828, -756, -683, -610, -538, -465, -392, -319, -247, -174, -101, -29, 44, 117, 189, 262, 335, 407, 480, 553, 625, 698, 771, 843, 916, 989, 1062, 1134, 1207, 1280, 1352, 1425, 1498, 1570, 1643, 1716, 1788, 1861, 1934, 2006, 2079, 2152, 2225, 2297, 2370, 2443, 2515, 2588, 2661, 2733, 2806, 2879, 2951, 3024, 3097, 3169, 3242, 3315, 3388, 3460, 3533]<br>[90], [-3154, -3081, -3009, -2936, -2863, -2791, -2718, -2645, -2573, -2500, -2427, -2355, -2282, -2209, -2137, -2064, -1991, -1918, -1846, -1773, -1700, -1628, -1555, -1482, -1410, -1337, -1264, -1192, -1119, -1046, -974, -901, -828, -756, -683, -610, -537, -465, -392, -319, -247, -174, -101, 44, 117, 189, 262, 335, 407, 480, 553, 626, 698, 771, 844, 916, 989, 1062, 1134, 1207, 1280, 1352, 1425, 1498, 1570, 1643, 1716, 1788, 1861, 1934, 2007, 2079, 2152, 2225, 2297, 2370, 2443, 2515, 2588, 2661, 2733, 2806, 2879, 2951, 3024, 3097, 3170, 3242, 3315, 3388, 3460, 3533]<br>[270], [-3154, -3082, -3009, -2936, -2863, -2791, -2718, -2645, -2573, -2500, -2427, -2355, -2282, -2209, -2137, -2064, -1991, -1919, -1846, -1773, -1701, -1628, -1555, -1482, -1410, -1337, -1264, -1192, -1119, -1046, -974, -901, -828, -756, -683, -610, -538, -465, -392, -319, -247, -174, -101, 189, 262, 335, 407, 480, 553, 625, 698, 771, 844, 916, 989, 1062, 1134, 1207, 1280, 1352, 1425, 1498, 1570, 1643, 1716, 1788, 1861, 1934, 2006, 2079, 2152, 2225, 2297, 2370, 2443, 2515, 2588, 2661, 2733, 2806, 2879, 2951, 3024, 3097, 3169, 3242, 3315, 3388, 3460, 3533]<br>[810], [-3154, -3081, -3009, -2936, -2863, -2791, -2718, -2645, -2573, -2500, -2427, -2355, -2282, -2209, -2137, -2064, -1991, -1919, -1846, -1773, -1700, -1628, -1555, -1482, -1410, -1337, -1264, -1192, -1119, -1046, -974, -901, -828, -756, -683, -610, -537, -465, -392, -319, -247, -29, 44, 189, 262, 335, 407, 480, 553, 626, 698, 771, 844, 916, 989, 1062, 1134, 1207, 1280, 1352, 1425, 1498, 1570, 1643, 1716, 1788, 1861, 1934, 2007, 2079, 2152, 2225, 2297, 2370, |

| | |
|---|---|
| | 2443, 2515, 2588, 2661, 2733, 2806, 2879, 2951, 3024, 3097, 3170, 3242, 3315, 3388, 3460, 3533]<br>[2430], [-3154, -3082, -3009, -2936, -2864, -2791, -2718, -2645, -2573, -2500, -2427, -2355, -2282, -2209, -2137, -2064, -1991, -1919, -1846, -1773, -1701, -1628, -1555, -1482, -1410, -1337, -1264, -1192, -1119, -1046, -974, -901, -828, -756, -683, -610, -538, -465, -392, -319, 44, 117, 189, 335, 407, 480, 553, 625, 698, 771, 843, 916, 989, 1062, 1134, 1207, 1280, 1352, 1425, 1498, 1570, 1643, 1716, 1788, 1861, 1934, 2006, 2079, 2152, 2225, 2297, 2370, 2443, 2515, 2588, 2661, 2733, 2806, 2879, 2951, 3024, 3097, 3169, 3242, 3315, 3388, 3460, 3533]<br>[5000], [-3154, -3081, -3009, -2936, -2863, -2791, -2718, -2645, -2573, -2500, -2427, -2355, -2282, -2209, -2137, -2064, -1991, -1918, -1846, -1773, -1700, -1628, -1555, -1482, -1410, -1337, -1264, -1192, -1119, -1046, -901, -828, -756, -683, -610, -537, -392, -174, -101, 407, 480, 553, 626, 698, 771, 916, 989, 1062, 1134, 1207, 1280, 1352, 1425, 1498, 1570, 1643, 1716, 1789, 1861, 1934, 2007, 2079, 2152, 2225, 2297, 2370, 2443, 2515, 2588, 2661, 2733, 2806, 2879, 2951, 3024, 3097, 3170, 3242, 3315, 3388, 3460, 3533] |
| AAG, 25ºC<br>$T_{EX}$ = 100 ms<br>600 MHz | [10], [-3157, -3084, -3012, -2939, -2866, -2794, -2721, -2648, -2576, -2503, -2430, -2357, -2285, -2212, -2139, -2067, -1994, -1921, -1849, -1776, -1703, -1631, -1558, -1485, -1413, -1340, -1267, -1195, -1122, -1049, -976, -904, -831, -758, -686, -613, -540, -468, -395, -322, -250, -177, -104, -32, 41, 114, 187, 259, 332, 405, 477, 550, 623, 695, 768, 841, 913, 986, 1059, 1131, 1204, 1277, 1350, 1422, 1495, 1568, 1640, 1713, 1786, 1858, 1931, 2004, 2076, 2149, 2222, 2294, 2367, 2440, 2512, 2585, 2658, 2731, 2803, 2876, 2949, 3021, 3094, 3167, 3239, 3312, 3385, 3457, 3530]<br>[30], [-3157, -3084, -3012, -2939, -2866, -2794, -2721, -2648, -2576, -2503, -2430, -2357, -2285, -2212, -2139, -2067, -1994, -1921, -1849, -1776, -1703, -1631, -1558, -1485, -1413, -1340, -1267, -1195, -1122, -1049, -976, -904, -831, -758, -686, -613, -540, -468, -395, -322, -250, -177, -104, -32, 41, 114, 187, 259, 332, 405, 477, 550, 623, 695, 768, 841, 913, 986, 1059, 1131, 1204, 1277, 1350, 1422, 1495, 1568, 1640, 1713, 1786, 1858, 1931, 2004, 2076, 2149, 2222, 2294, 2367, 2440, 2512, 2585, 2658, 2731, 2803, 2876, 2949, 3021, 3094, 3167, 3239, 3312, 3385, 3457, 3530]<br>[90], [-3157, -3084, -3012, -2939, -2866, -2794, -2721, -2648, -2576, -2503, -2430, -2357, -2285, -2212, -2139, -2067, -1994, -1921, -1849, -1776, -1703, -1631, -1558, -1485, -1413, -1340, -1267, -1194, -1122, -1049, -976, -904, -831, -758, -686, -613, -540, -468, -395, -322, -250, -177, -104, -32, 114, 187, 259, 332, 405, 477, 550, 623, 695, 768, 841, 913, 986, 1059, 1131, 1204, 1277, 1350, 1422, 1495, 1568, 1640, 1713, 1786, 1858, 1931, 2004, 2076, 2149, 2222, 2294, 2367, 2440, 2512, 2585, 2658, 2731, 2803, 2876, 2949, 3021, 3094, 3167, 3239, 3312, 3385, 3457, 3530]<br>[270], [-3157, -3084, -3012, -2939, -2866, -2794, -2721, -2648, -2576, -2503, -2430, -2358, -2285, -2212, -2139, -2067, -1994, -1921, -1849, -1776, -1703, -1631, -1558, -1485, -1413, -1340, -1267, -1195, -1122, -1049, -976, -904, -831, |

| | |
|---|---|
| | -758, -686, -613, -540, -468, -395, -322, -250, -177, -32, 41, 114, 187, 259, 332, 405, 477, 550, 623, 695, 768, 841, 913, 986, 1059, 1131, 1204, 1277, 1349, 1422, 1495, 1568, 1640, 1713, 1786, 1858, 1931, 2004, 2076, 2149, 2222, 2294, 2367, 2440, 2512, 2585, 2658, 2731, 2803, 2876, 2949, 3021, 3094, 3167, 3239, 3312, 3385, 3457, 3530]<br>[810], [-3157, -3084, -3012, -2939, -2866, -2794, -2721, -2648, -2575, -2503, -2430, -2357, -2285, -2212, -2139, -2067, -1994, -1921, -1849, -1776, -1703, -1631, -1558, -1485, -1412, -1340, -1267, -1194, -1122, -1049, -976, -904, -831, -758, -686, -613, -540, -468, -395, -322, -249, -177, 41, 114, 187, 259, 332, 405, 477, 550, 623, 695, 768, 841, 913, 986, 1059, 1132, 1204, 1277, 1350, 1422, 1495, 1568, 1640, 1713, 1786, 1858, 1931, 2004, 2076, 2149, 2222, 2295, 2367, 2440, 2513, 2585, 2658, 2731, 2803, 2876, 2949, 3021, 3094, 3167, 3239, 3312, 3385, 3457, 3530]<br>[2430], [-3157, -3084, -3012, -2939, -2866, -2794, -2721, -2648, -2576, -2503, -2430, -2357, -2285, -2212, -2139, -2067, -1994, -1921, -1849, -1776, -1703, -1631, -1558, -1485, -1413, -1340, -1267, -1194, -1122, -1049, -976, -904, -831, -758, -686, -613, -540, -468, -395, -177, -32, 259, 332, 405, 477, 550, 623, 695, 768, 841, 913, 986, 1059, 1131, 1204, 1277, 1350, 1422, 1495, 1568, 1640, 1713, 1786, 1858, 1931, 2004, 2076, 2149, 2222, 2294, 2367, 2440, 2513, 2585, 2658, 2731, 2803, 2876, 2949, 3021, 3094, 3167, 3239, 3312, 3385, 3457, 3530]<br>[5000], [-3157, -3084, -3012, -2939, -2866, -2794, -2721, -2648, -2576, -2503, -2430, -2357, -2285, -2212, -2139, -2067, -1994, -1921, -1849, -1776, -1703, -1631, -1558, -1485, -1413, -1340, -1267, -1194, -1049, -976, -686, -468, -322, -250, -177, -104, 187, 332, 550, 623, 695, 768, 913, 1059, 1131, 1277, 1350, 1422, 1495, 1568, 1640, 1713, 1786, 1858, 1931, 2004, 2076, 2149, 2222, 2294, 2367, 2440, 2513, 2585, 2658, 2731, 2803, 2876, 2949, 3021, 3094, 3167, 3239, 3312, 3385, 3457, 3530] |
| AAC, 25ºC<br>$T_{EX}$ = 100 ms<br>600 MHz | [10], [-3191, -3118, -3045, -2972, -2900, -2827, -2754, -2682, -2609, -2536, -2464, -2391, -2318, -2246, -2173, -2100, -2028, -1955, -1882, -1810, -1737, -1664, -1591, -1519, -1446, -1373, -1301, -1228, -1155, -1083, -1010, -937, -865, -792, -719, -647, -574, -501, -428, -356, -283, -210, -138, -65, 8, 80, 153, 226, 298, 371, 444, 516, 589, 662, 734, 807, 880, 953, 1025, 1098, 1171, 1243, 1316, 1389, 1461, 1534, 1607, 1679, 1752, 1825, 1897, 1970, 2043, 2116, 2188, 2261, 2334, 2406, 2479, 2552, 2624, 2697, 2770, 2842, 2915, 2988, 3060, 3133, 3206, 3279, 3351, 3424, 3497, 3569]<br>[30], [-3190, -3118, -3045, -2972, -2900, -2827, -2754, -2682, -2609, -2536, -2464, -2391, -2318, -2246, -2173, -2100, -2028, -1955, -1882, -1809, -1737, -1664, -1591, -1519, -1446, -1373, -1301, -1228, -1155, -1083, -1010, -937, -865, -792, -719, -646, -574, -501, -428, -356, -283, -210, -138, -65, 8, 80, 153, 226, 298, 371, 444, 517, 589, 662, 735, 807, 880, 953, 1025, 1098, 1171, 1243, 1316, 1389, 1461, 1534, 1607, 1679, 1752, 1825, 1898, 1970, 2043, 2116, 2188, 2261, 2334, 2406, 2479, 2552, 2624, 2697, 2770, 2842, 2915, 2988, 3061, 3133, 3206, 3279, 3351, 3424, 3497, 3569] |

| | |
|---|---|
| | [90], [-3190, -3118, -3045, -2972, -2900, -2827, -2754, -2682, -2609, -2536, -2464, -2391, -2318, -2246, -2173, -2100, -2027, -1955, -1882, -1809, -1737, -1664, -1591, -1519, -1446, -1373, -1301, -1228, -1155, -1083, -1010, -937, -864, -792, -719, -646, -574, -501, -428, -356, -283, -210, -138, -65, 8, 80, 153, 226, 299, 371, 444, 517, 589, 662, 735, 807, 880, 953, 1025, 1098, 1171, 1243, 1316, 1389, 1461, 1534, 1607, 1680, 1752, 1825, 1898, 1970, 2043, 2116, 2188, 2261, 2334, 2406, 2479, 2552, 2624, 2697, 2770, 2843, 2915, 2988, 3061, 3133, 3206, 3279, 3351, 3424, 3497, 3569]<br>[270], [-3190, -3118, -3045, -2972, -2900, -2827, -2754, -2682, -2609, -2536, -2464, -2391, -2318, -2246, -2173, -2100, -2028, -1955, -1882, -1809, -1737, -1664, -1591, -1519, -1446, -1373, -1301, -1228, -1155, -1083, -1010, -937, -865, -792, -719, -646, -574, -501, -428, -356, -283, -210, -138, 8, 80, 153, 226, 298, 371, 444, 517, 589, 662, 735, 807, 880, 953, 1025, 1098, 1171, 1243, 1316, 1389, 1461, 1534, 1607, 1679, 1752, 1825, 1898, 1970, 2043, 2116, 2188, 2261, 2334, 2406, 2479, 2552, 2624, 2697, 2770, 2842, 2915, 2988, 3061, 3133, 3206, 3279, 3351, 3424, 3497, 3569]<br>[810], [-3190, -3118, -3045, -2972, -2900, -2827, -2754, -2682, -2609, -2536, -2464, -2391, -2318, -2246, -2173, -2100, -2027, -1955, -1882, -1809, -1737, -1664, -1591, -1519, -1446, -1373, -1301, -1228, -1155, -1083, -1010, -937, -864, -792, -719, -646, -574, -501, -428, -356, -283, -210, -138, -65, 153, 226, 298, 371, 444, 517, 589, 662, 735, 807, 880, 953, 1025, 1098, 1171, 1243, 1316, 1389, 1461, 1534, 1607, 1680, 1752, 1825, 1898, 1970, 2043, 2116, 2188, 2261, 2334, 2406, 2479, 2552, 2624, 2697, 2770, 2843, 2915, 2988, 3061, 3133, 3206, 3279, 3351, 3424, 3497, 3569]<br>[2430], [-3190, -3118, -3045, -2972, -2900, -2827, -2754, -2682, -2609, -2536, -2464, -2391, -2318, -2246, -2173, -2100, -2028, -1955, -1882, -1809, -1737, -1664, -1591, -1519, -1446, -1373, -1301, -1228, -1155, -1083, -1010, -937, -865, -792, -719, -646, -574, -501, -428, -356, -65, 153, 298, 371, 444, 662, 735, 807, 880, 953, 1025, 1098, 1171, 1243, 1316, 1389, 1461, 1534, 1607, 1679, 1752, 1825, 1898, 1970, 2043, 2116, 2188, 2261, 2334, 2406, 2479, 2552, 2624, 2697, 2770, 2842, 2915, 2988, 3061, 3133, 3206, 3279, 3351, 3424, 3497, 3569]<br>[5000], [-3190, -3118, -3045, -2972, -2900, -2827, -2754, -2682, -2609, -2536, -2464, -2391, -2318, -2246, -2173, -2100, -2027, -1955, -1882, -1809, -1737, -1664, -1591, -1519, -1446, -1373, -1301, -1228, -1155, -1083, -1010, -937, -864, -646, -501, -356, 8, 299, 735, 807, 880, 953, 1098, 1171, 1243, 1316, 1389, 1461, 1534, 1607, 1680, 1752, 1825, 1898, 1970, 2043, 2116, 2188, 2261, 2334, 2406, 2479, 2552, 2624, 2697, 2770, 2843, 2915, 2988, 3061, 3133, 3206, 3279, 3351, 3424, 3497, 3569] |
| CAT, 25ºC<br>$T_{EX}$ = 100 ms<br>600 MHz | [30], [-3458, -3386, -3313, -3240, -3168, -3095, -3022, -2950, -2877, -2804, -2732, -2659, -2586, -2514, -2441, -2368, -2295, -2223, -2150, -2077, -2005, -1932, -1859, -1787, -1714, -1641, -1569, -1496, -1423, -1351, -1278, -1205, -1133, -1060, -987, -914, -842, -769, -696, -624, -551, -478, -406, -333, -260, -188, -115, -42, 30, 103, 176, 249, 321, 394, 467, 539, 612, 685, 757, 830, 903, 975, 1048, 1121, 1193, 1266, 1339, 1411, 1484, 1557, 1630, 1702, 1775, 1848, |

| | |
|---|---|
| | 1920, 1993, 2066, 2138, 2211, 2284, 2356, 2429, 2502, 2574, 2647, 2720, 2793, 2865, 2938, 3011, 3083, 3156, 3229, 3301, 3374, 3447, 3519, 3592]<br>[90], [-3458, -3386, -3313, -3240, -3168, -3095, -3022, -2950, -2877, -2804, -2731, -2659, -2586, -2513, -2441, -2368, -2295, -2223, -2150, -2077, -2005, -1932, -1859, -1787, -1714, -1641, -1568, -1496, -1423, -1350, -1278, -1205, -1132, -1060, -987, -914, -842, -769, -696, -624, -551, -478, -406, -333, -260, -187, -115, -42, 103, 176, 249, 321, 394, 467, 539, 612, 685, 757, 830, 903, 976, 1048, 1121, 1194, 1266, 1339, 1412, 1484, 1557, 1630, 1702, 1775, 1848, 1920, 1993, 2066, 2139, 2211, 2284, 2357, 2429, 2502, 2575, 2647, 2720, 2793, 2865, 2938, 3011, 3083, 3156, 3229, 3301, 3374, 3447, 3520, 3592]<br>[270], [-3458, -3386, -3313, -3240, -3168, -3095, -3022, -2950, -2877, -2804, -2732, -2659, -2586, -2513, -2441, -2368, -2295, -2223, -2150, -2077, -2005, -1932, -1859, -1787, -1714, -1641, -1569, -1496, -1423, -1350, -1278, -1205, -1132, -1060, -987, -914, -842, -769, -696, -624, -551, -478, -406, -333, -260, -188, -115, -42, 31, 176, 249, 321, 394, 467, 539, 612, 685, 757, 830, 903, 975, 1048, 1121, 1194, 1266, 1339, 1412, 1484, 1557, 1630, 1702, 1775, 1848, 1920, 1993, 2066, 2138, 2211, 2284, 2357, 2429, 2502, 2575, 2647, 2720, 2793, 2865, 2938, 3011, 3083, 3156, 3229, 3301, 3374, 3447, 3519, 3592]<br>[810], [-3458, -3386, -3313, -3240, -3168, -3095, -3022, -2949, -2877, -2804, -2731, -2659, -2586, -2513, -2441, -2368, -2295, -2223, -2150, -2077, -2005, -1932, -1859, -1786, -1714, -1641, -1568, -1496, -1423, -1350, -1278, -1205, -1132, -1060, -987, -914, -842, -769, -696, -624, -551, -478, -405, -333, -115, 103, 249, 321, 394, 467, 539, 612, 685, 758, 830, 903, 976, 1048, 1121, 1194, 1266, 1339, 1412, 1484, 1557, 1630, 1702, 1775, 1848, 1921, 1993, 2066, 2139, 2211, 2284, 2357, 2429, 2502, 2575, 2647, 2720, 2793, 2865, 2938, 3011, 3083, 3156, 3229, 3302, 3374, 3447, 3520, 3592]<br>[2430], [-3458, -3386, -3313, -3240, -3168, -3095, -3022, -2950, -2877, -2804, -2731, -2659, -2586, -2513, -2441, -2368, -2295, -2223, -2150, -2077, -2005, -1932, -1859, -1787, -1714, -1641, -1568, -1496, -1423, -1350, -1278, -1205, -1132, -1060, -987, -914, -842, -769, -696, -624, -551, -478, -333, -260, -115, -42, 31, 321, 394, 539, 612, 685, 757, 830, 903, 976, 1048, 1121, 1194, 1266, 1339, 1412, 1484, 1557, 1630, 1702, 1775, 1848, 1920, 1993, 2066, 2139, 2211, 2284, 2357, 2429, 2502, 2575, 2647, 2720, 2793, 2865, 2938, 3011, 3083, 3156, 3229, 3301, 3374, 3447, 3520, 3592]<br>[5000], [-3458, -3386, -3313, -3240, -3168, -3095, -3022, -2949, -2877, -2804, -2731, -2659, -2586, -2513, -2441, -2368, -2295, -2223, -2150, -2077, -2005, -1932, -1859, -1786, -1714, -1641, -1568, -1423, -1350, -1278, -1205, -1132, -1060, -987, -914, -842, -769, -551, -405, -333, -260, 31, 103, 176, 249, 685, 758, 830, 903, 976, 1048, 1121, 1194, 1266, 1339, 1412, 1484, 1557, 1630, 1702, 1775, 1848, 1921, 1993, 2066, 2139, 2211, 2284, 2357, 2429, 2502, 2575, 2647, 2720, 2793, 2865, 2938, 3011, 3083, 3156, 3229, 3302, 3374, 3447, 3520, 3592] |
| GAT, 25ºC $T_{EX}$ = 100 ms | [10], [-3415, -3342, -3269, -3197, -3124, -3051, -2979, -2906, -2833, -2761, -2688, -2615, -2542, -2470, -2397, -2324, -2252, -2179, -2106, -2034, -1961, -1888, -1816, -1743, -1670, -1598, -1525, -1452, -1380, -1307, -1234, -1161, - |

| | |
|---|---|
| 600 MHz | 1089, -1016, -943, -871, -798, -725, -653, -580, -507, -435, -362, -289, -217, -144, -71, 2, 74, 147, 220, 292, 365, 438, 510, 583, 656, 728, 801, 874, 946, 1019, 1092, 1164, 1237, 1310, 1383, 1455, 1528, 1601, 1673, 1746, 1819, 1891, 1964, 2037, 2109, 2182, 2255, 2327, 2400, 2473, 2546, 2618, 2691, 2764, 2836, 2909, 2982, 3054, 3127, 3200, 3272, 3345, 3418, 3490, 3563]<br>[30], [-3415, -3342, -3269, -3197, -3124, -3051, -2979, -2906, -2833, -2761, -2688, -2615, -2542, -2470, -2397, -2324, -2252, -2179, -2106, -2034, -1961, -1888, -1816, -1743, -1670, -1598, -1525, -1452, -1379, -1307, -1234, -1161, -1089, -1016, -943, -871, -798, -725, -653, -580, -507, -435, -362, -289, -217, -144, -71, 2, 74, 147, 220, 292, 365, 438, 510, 583, 656, 728, 801, 874, 946, 1019, 1092, 1165, 1237, 1310, 1383, 1455, 1528, 1601, 1673, 1746, 1819, 1891, 1964, 2037, 2255, 2328, 2400, 2473, 2546, 2618, 2691, 2764, 2836, 2909, 2982, 3054, 3127, 3200, 3272, 3345, 3418, 3490, 3563]<br>[90], [-3415, -3342, -3269, -3197, -3124, -3051, -2979, -2906, -2833, -2760, -2688, -2615, -2542, -2470, -2397, -2324, -2252, -2179, -2106, -2034, -1961, -1888, -1816, -1743, -1670, -1597, -1525, -1452, -1379, -1307, -1234, -1161, -1089, -1016, -943, -871, -798, -725, -653, -580, -507, -434, -362, -289, -216, -144, -71, 74, 147, 220, 292, 365, 438, 510, 583, 656, 728, 801, 874, 947, 1019, 1092, 1165, 1237, 1310, 1383, 1455, 1528, 1601, 1673, 1746, 1819, 1891, 1964, 2037, 2110, 2182, 2255, 2328, 2400, 2473, 2546, 2618, 2691, 2764, 2836, 2909, 2982, 3054, 3127, 3200, 3272, 3345, 3418, 3491, 3563]<br>[270], [-3415, -3342, -3269, -3196, -3124, -3051, -2978, -2906, -2833, -2760, -2688, -2615, -2542, -2470, -2397, -2324, -2252, -2179, -2106, -2033, -1961, -1888, -1815, -1743, -1670, -1597, -1525, -1452, -1379, -1307, -1234, -1161, -1089, -1016, -943, -871, -798, -725, -652, -580, -507, -434, -362, -289, -216, -144, 147, 220, 292, 365, 438, 511, 583, 656, 729, 801, 874, 947, 1019, 1092, 1165, 1237, 1310, 1383, 1455, 1528, 1601, 1674, 1746, 1819, 1892, 1964, 2037, 2110, 2182, 2255, 2328, 2400, 2473, 2546, 2618, 2691, 2764, 2836, 2909, 2982, 3055, 3127, 3200, 3273, 3345, 3418, 3491, 3563]<br>[810], [-3415, -3342, -3269, -3196, -3124, -3051, -2978, -2906, -2833, -2760, -2688, -2615, -2542, -2470, -2397, -2324, -2252, -2179, -2106, -2034, -1961, -1888, -1815, -1743, -1670, -1597, -1525, -1452, -1379, -1307, -1234, -1161, -1089, -1016, -943, -871, -798, -725, -652, -580, -507, -434, -362, -289, -216, -71, 74, 147, 220, 292, 365, 438, 510, 583, 656, 729, 801, 874, 947, 1019, 1092, 1165, 1237, 1310, 1383, 1455, 1528, 1601, 1673, 1746, 1819, 1892, 1964, 2037, 2110, 2182, 2255, 2328, 2400, 2473, 2546, 2618, 2691, 2764, 2836, 2909, 2982, 3055, 3127, 3200, 3273, 3345, 3418, 3491, 3563]<br>[2430], [-3415, -3342, -3269, -3197, -3124, -3051, -2979, -2906, -2833, -2760, -2688, -2615, -2542, -2470, -2397, -2324, -2252, -2179, -2106, -2034, -1961, -1888, -1816, -1743, -1670, -1597, -1525, -1452, -1379, -1307, -1234, -1161, -1089, -1016, -943, -871, -798, -725, -653, -580, -507, -362, -289, -216, 2, 74, 147, 220, 292, 365, 438, 510, 583, 656, 728, 801, 874, 947, 1019, 1092, 1165, 1237, 1310, 1383, 1455, 1528, 1601, 1673, 1746, 1819, 1891, 1964, 2037, |

| | |
|---|---|
| | 2109, 2182, 2255, 2328, 2400, 2473, 2546, 2618, 2691, 2764, 2836, 2909, 2982, 3054, 3127, 3200, 3272, 3345, 3418, 3491, 3563]<br>[5000], [-3415, -3342, -3269, -3197, -3124, -3051, -2978, -2906, -2833, -2760, -2688, -2615, -2542, -2470, -2397, -2324, -2252, -2179, -2106, -2034, -1961, -1888, -1815, -1743, -1670, -1597, -1525, -1452, -1379, -1307, -1234, -1089, -1016, -943, -871, -798, -580, -362, -71, 74, 147, 292, 365, 510, 583, 656, 801, 947, 1019, 1165, 1237, 1310, 1383, 1455, 1528, 1601, 1673, 1746, 1819, 1892, 1964, 2037, 2110, 2182, 2255, 2328, 2400, 2473, 2546, 2618, 2691, 2764, 2836, 2909, 2982, 3054, 3127, 3200, 3273, 3345, 3418, 3491, 3563] |
| TAC, 25ºC<br>$T_{EX}$ = 100 ms<br>600 MHz | [10], [-3501, -3428, -3355, -3283, -3210, -3137, -3065, -2992, -2919, -2847, -2774, -2701, -2629, -2556, -2483, -2410, -2338, -2265, -2192, -2120, -2047, -1974, -1902, -1829, -1756, -1684, -1611, -1538, -1466, -1393, -1320, -1247, -1175, -1102, -1029, -957, -884, -811, -739, -666, -593, -521, -448, -375, -303, -230, -157, -85, -12, 61, 134, 206, 279, 352, 424, 497, 570, 642, 715, 788, 860, 933, 1006, 1078, 1151, 1224, 1297, 1369, 1442, 1515, 1587, 1660, 1733, 1805, 1878, 1951, 2023, 2096, 2169, 2241, 2314, 2387, 2460, 2532, 2605, 2678, 2750, 2823, 2896, 2968, 3041, 3114, 3186, 3259, 3332, 3404, 3477, 3550]<br>[30], [-3501, -3428, -3355, -3283, -3210, -3137, -3065, -2992, -2919, -2847, -2774, -2701, -2629, -2556, -2483, -2411, -2338, -2265, -2193, -2120, -2047, -1974, -1902, -1829, -1756, -1684, -1611, -1538, -1466, -1393, -1320, -1248, -1175, -1102, -1030, -957, -884, -811, -739, -666, -593, -521, -448, -375, -303, -230, -157, -85, 61, 133, 206, 279, 352, 424, 497, 570, 642, 715, 788, 860, 933, 1006, 1078, 1151, 1224, 1296, 1369, 1442, 1514, 1587, 1660, 1733, 1805, 1878, 1951, 2023, 2096, 2169, 2241, 2314, 2387, 2459, 2532, 2605, 2677, 2750, 2823, 2896, 2968, 3041, 3114, 3186, 3259, 3332, 3404, 3477, 3550]<br>[90], [-3501, -3428, -3356, -3283, -3210, -3138, -3065, -2992, -2919, -2847, -2774, -2701, -2629, -2556, -2483, -2411, -2338, -2265, -2193, -2120, -2047, -1975, -1902, -1829, -1756, -1684, -1611, -1538, -1466, -1393, -1320, -1248, -1175, -1102, -1030, -957, -884, -812, -739, -666, -593, -521, -448, -375, -303, -230, -157, -85, 61, 133, 206, 279, 351, 424, 497, 569, 642, 715, 788, 860, 933, 1006, 1078, 1151, 1224, 1296, 1369, 1442, 1514, 1587, 1660, 1732, 1805, 1878, 1951, 2023, 2096, 2169, 2241, 2314, 2387, 2459, 2532, 2605, 2677, 2750, 2823, 2895, 2968, 3041, 3113, 3186, 3259, 3332, 3404, 3477, 3550]<br>[270], [-3501, -3428, -3355, -3283, -3210, -3137, -3065, -2992, -2919, -2847, -2774, -2701, -2629, -2556, -2483, -2410, -2338, -2265, -2192, -2120, -2047, -1974, -1902, -1829, -1756, -1684, -1611, -1538, -1466, -1393, -1320, -1247, -1175, -1102, -1029, -957, -884, -811, -739, -666, -593, -521, -448, -375, -303, -230, -157, -85, -12, 134, 206, 279, 352, 424, 497, 570, 642, 715, 788, 860, 933, 1006, 1078, 1151, 1224, 1297, 1369, 1442, 1515, 1587, 1660, 1733, 1805, 1878, 1951, 2023, 2096, 2169, 2241, 2314, 2387, 2459, 2532, 2605, 2678, 2750, 2823, 2896, 2968, 3041, 3114, 3186, 3259, 3332, 3404, 3477, 3550]<br>[810], [-3501, -3428, -3355, -3283, -3210, -3137, -3065, -2992, -2919, -2847, -2774, -2701, -2629, -2556, -2483, -2411, -2338, -2265, -2192, -2120, -2047, -1974, -1902, -1829, -1756, -1684, -1611, -1538, -1466, -1393, -1320, -1248, - |

| | |
|---|---|
| | 1175, -1102, -1030, -957, -884, -811, -739, -666, -593, -521, -448, -375, -303, -230, -157, -85, 61, 133, 206, 279, 352, 424, 497, 570, 642, 715, 788, 860, 933, 1006, 1078, 1151, 1224, 1296, 1369, 1442, 1514, 1587, 1660, 1733, 1805, 1878, 1951, 2023, 2096, 2169, 2241, 2314, 2387, 2459, 2532, 2605, 2677, 2750, 2823, 2896, 2968, 3041, 3114, 3186, 3259, 3332, 3404, 3477, 3550]<br>[2430], [-3501, -3428, -3356, -3283, -3210, -3138, -3065, -2992, -2919, -2847, -2774, -2701, -2629, -2556, -2483, -2411, -2338, -2265, -2193, -2120, -2047, -1975, -1902, -1829, -1757, -1684, -1611, -1538, -1466, -1393, -1320, -1248, -1175, -1102, -1030, -957, -884, -812, -739, -666, -594, -303, -85, -12, 61, 279, 351, 424, 497, 715, 860, 933, 1006, 1078, 1151, 1224, 1296, 1369, 1442, 1514, 1587, 1660, 1732, 1805, 1878, 1950, 2023, 2096, 2169, 2241, 2314, 2387, 2459, 2532, 2605, 2677, 2750, 2823, 2895, 2968, 3041, 3113, 3186, 3259, 3332, 3404, 3477, 3550]<br>[5000], [-3501, -3428, -3356, -3283, -3210, -3138, -3065, -2992, -2919, -2847, -2774, -2701, -2629, -2556, -2483, -2411, -2338, -2265, -2193, -2120, -2047, -1975, -1902, -1829, -1756, -1684, -1611, -1538, -1393, -1248, -1175, -1102, -957, -375, -303, -230, -85, 61, 133, 206, 351, 497, 569, 642, 715, 788, 860, 1078, 1296, 1369, 1442, 1514, 1587, 1732, 1805, 1878, 1951, 2023, 2096, 2169, 2241, 2314, 2387, 2459, 2532, 2605, 2677, 2750, 2823, 2895, 2968, 3041, 3113, 3186, 3259, 3332, 3404, 3477, 3550] |
| GAG, 25ºC<br>$T_{EX}$ = 100 ms<br>600 MHz | [10], [-3265, -3174, -3083, -2992, -2901, -2810, -2719, -2628, -2537, -2446, -2354, -2263, -2172, -2081, -1990, -1899, -1808, -1717, -1626, -1535, -1444, -1352, -1261, -1170, -1079, -988, -897, -806, -715, -624, -533, -442, -351, -259, -168, -77, 14, 105, 196, 287, 378, 469, 560, 651, 743, 834, 925, 1016, 1107, 1198, 1289, 1380, 1471, 1562, 1653, 1744, 1836, 1927, 2018, 2109, 2200, 2291, 2382, 2473, 2564, 2655, 2746, 2838, 2929, 3020, 3111, 3202, 3293, 3384, 3475, 3566]<br>[30], [-3265, -3174, -3083, -2992, -2901, -2810, -2719, -2628, -2537, -2445, -2354, -2263, -2172, -2081, -1990, -1899, -1808, -1717, -1626, -1535, -1443, -1352, -1261, -1170, -1079, -988, -897, -806, -715, -624, -533, -442, -350, -259, -168, -77, 14, 105, 196, 287, 378, 469, 560, 652, 743, 834, 925, 1016, 1107, 1198, 1289, 1380, 1471, 1562, 1653, 1745, 1836, 1927, 2018, 2109, 2200, 2291, 2382, 2473, 2564, 2655, 2747, 2838, 2929, 3020, 3111, 3202, 3293, 3384, 3475, 3566]<br>[90], [-3265, -3174, -3083, -2992, -2901, -2810, -2719, -2628, -2536, -2445, -2354, -2263, -2172, -2081, -1990, -1899, -1808, -1717, -1626, -1534, -1443, -1352, -1261, -1170, -1079, -988, -897, -806, -715, -624, -533, -441, -350, -259, -168, -77, 105, 196, 287, 378, 469, 561, 652, 743, 834, 925, 1016, 1107, 1198, 1289, 1380, 1471, 1562, 1654, 1745, 1836, 1927, 2018, 2109, 2200, 2291, 2382, 2473, 2564, 2656, 2747, 2838, 2929, 3020, 3111, 3202, 3293, 3384, 3475, 3566]<br>[270], [-3265, -3174, -3083, -2992, -2901, -2810, -2719, -2628, -2537, -2445, -2354, -2263, -2172, -2081, -1990, -1899, -1808, -1717, -1626, -1535, -1444, -1352, -1261, -1170, -1079, -988, -897, -806, -715, -624, -533, -442, -350, -259, |

| | |
|---|---|
| | -168, -77, 14, 105, 196, 287, 378, 469, 560, 651, 743, 834, 925, 1016, 1107, 1198, 1289, 1380, 1471, 1562, 1653, 1745, 1836, 1927, 2018, 2109, 2200, 2291, 2382, 2473, 2564, 2655, 2746, 2838, 2929, 3020, 3111, 3202, 3293, 3384, 3475, 3566]<br>[810], [-3265, -3174, -3083, -2992, -2901, -2810, -2719, -2628, -2536, -2445, -2354, -2263, -2172, -2081, -1990, -1899, -1808, -1717, -1626, -1535, -1443, -1352, -1261, -1170, -1079, -988, -897, -806, -715, -624, -533, -441, -350, -259, -168, -77, 105, 196, 287, 378, 469, 561, 652, 743, 834, 925, 1016, 1107, 1198, 1289, 1380, 1471, 1562, 1654, 1745, 1836, 1927, 2018, 2109, 2200, 2291, 2382, 2473, 2564, 2656, 2747, 2838, 2929, 3020, 3111, 3202, 3293, 3384, 3475, 3566]<br>[2430], [-3265, -3174, -3083, -2992, -2901, -2810, -2719, -2628, -2537, -2446, -2354, -2263, -2172, -2081, -1990, -1899, -1808, -1717, -1626, -1535, -1444, -1352, -1261, -1170, -1079, -988, -897, -806, -715, -624, -533, -442, -351, -168, 287, 378, 469, 560, 651, 743, 834, 925, 1016, 1107, 1198, 1289, 1380, 1471, 1562, 1653, 1744, 1836, 1927, 2018, 2109, 2200, 2291, 2382, 2473, 2564, 2655, 2746, 2838, 2929, 3020, 3111, 3202, 3293, 3384, 3475, 3566]<br>[5000], [-3265, -3174, -3083, -2992, -2901, -2810, -2719, -2628, -2537, -2445, -2354, -2263, -2172, -2081, -1990, -1899, -1808, -1717, -1626, -1535, -1444, -1352, -1261, -1170, -1079, -897, -806, -442, -350, -168, 14, 105, 196, 378, 651, 743, 834, 1016, 1107, 1198, 1289, 1380, 1471, 1562, 1653, 1745, 1836, 1927, 2018, 2109, 2200, 2291, 2382, 2473, 2564, 2655, 2747, 2838, 2929, 3020, 3111, 3202, 3293, 3384, 3475, 3566] |
| CAG, 25ºC<br>$T_{EX}$ = 100 ms<br>600 MHz | [10], [-3235, -3162, -3090, -3017, -2944, -2872, -2799, -2726, -2653, -2581, -2508, -2435, -2363, -2290, -2217, -2145, -2072, -1999, -1927, -1854, -1781, -1709, -1636, -1563, -1491, -1418, -1345, -1272, -1200, -1127, -1054, -982, -909, -836, -764, -691, -618, -546, -473, -400, -328, -255, -182, -109, -37, 36, 109, 181, 254, 327, 399, 472, 545, 617, 690, 763, 835, 908, 981, 1053, 1126, 1199, 1272, 1344, 1417, 1490, 1562, 1635, 1708, 1780, 1853, 1926, 1998, 2071, 2144, 2216, 2289, 2362, 2435, 2507, 2580, 2653, 2725, 2798, 2871, 2943, 3016, 3089, 3161, 3234, 3307, 3379, 3452, 3525, 3598]<br>[30], [-3235, -3162, -3090, -3017, -2944, -2872, -2799, -2726, -2653, -2581, -2508, -2435, -2363, -2290, -2217, -2145, -2072, -1999, -1927, -1854, -1781, -1709, -1636, -1563, -1490, -1418, -1345, -1272, -1200, -1127, -1054, -982, -909, -836, -764, -691, -618, -546, -473, -400, -327, -255, -182, -109, -37, 36, 109, 181, 254, 327, 399, 472, 545, 617, 690, 763, 835, 908, 981, 1054, 1126, 1199, 1272, 1344, 1417, 1490, 1562, 1635, 1708, 1780, 1853, 1926, 1998, 2071, 2144, 2217, 2289, 2362, 2435, 2507, 2580, 2653, 2725, 2798, 2871, 2943, 3016, 3089, 3161, 3234, 3307, 3379, 3452, 3525, 3598]<br>[90], [-3235, -3162, -3089, -3017, -2944, -2871, -2799, -2726, -2653, -2581, -2508, -2435, -2363, -2290, -2217, -2145, -2072, -1999, -1927, -1854, -1781, -1708, -1636, -1563, -1490, -1418, -1345, -1272, -1200, -1127, -1054, -982, -909, -836, -764, -691, -618, -545, -473, -400, -327, -255, -182, -109, -37, 36, 109, 181, 254, 327, 399, 472, 545, 617, 690, 763, 836, 908, 981, 1054, 1126, 1199, |

| | |
|---|---|
| | 1272, 1344, 1417, 1490, 1562, 1635, 1708, 1780, 1853, 1926, 1999, 2071, 2144, 2217, 2289, 2362, 2435, 2507, 2580, 2653, 2725, 2798, 2871, 2943, 3016, 3089, 3162, 3234, 3307, 3380, 3452, 3525, 3598]<br>[270], [-3235, -3162, -3090, -3017, -2944, -2872, -2799, -2726, -2653, -2581, -2508, -2435, -2363, -2290, -2217, -2145, -2072, -1999, -1927, -1854, -1781, -1709, -1636, -1563, -1490, -1418, -1345, -1272, -1200, -1127, -1054, -982, -909, -836, -764, -691, -618, -546, -473, -400, -328, -255, -182, -109, 109, 181, 254, 327, 399, 472, 545, 617, 690, 763, 835, 908, 981, 1054, 1126, 1199, 1272, 1344, 1417, 1490, 1562, 1635, 1708, 1780, 1853, 1926, 1998, 2071, 2144, 2216, 2289, 2362, 2435, 2507, 2580, 2653, 2725, 2798, 2871, 2943, 3016, 3089, 3161, 3234, 3307, 3379, 3452, 3525, 3598]<br>[810], [-3235, -3162, -3090, -3017, -2944, -2871, -2799, -2726, -2653, -2581, -2508, -2435, -2363, -2290, -2217, -2145, -2072, -1999, -1927, -1854, -1781, -1708, -1636, -1563, -1490, -1418, -1345, -1272, -1200, -1127, -1054, -982, -909, -836, -764, -691, -618, -546, -473, -400, -327, -255, -182, -37, 36, 109, 181, 254, 327, 399, 472, 545, 617, 690, 763, 836, 908, 981, 1054, 1126, 1199, 1272, 1344, 1417, 1490, 1562, 1635, 1708, 1780, 1853, 1926, 1999, 2071, 2144, 2217, 2289, 2362, 2435, 2507, 2580, 2653, 2725, 2798, 2871, 2943, 3016, 3089, 3161, 3234, 3307, 3380, 3452, 3525, 3598]<br>[2430], [-3235, -3162, -3090, -3017, -2944, -2872, -2799, -2726, -2654, -2581, -2508, -2435, -2363, -2290, -2217, -2145, -2072, -1999, -1927, -1854, -1781, -1709, -1636, -1563, -1491, -1418, -1345, -1272, -1200, -1127, -1054, -982, -909, -836, -764, -691, -618, -546, -473, -400, -255, -37, 327, 472, 545, 617, 690, 763, 835, 908, 981, 1053, 1126, 1199, 1272, 1344, 1417, 1490, 1562, 1635, 1708, 1780, 1853, 1926, 1998, 2071, 2144, 2216, 2289, 2362, 2435, 2507, 2580, 2653, 2725, 2798, 2871, 2943, 3016, 3089, 3161, 3234, 3307, 3379, 3452, 3525, 3597]<br>[5000], [-3235, -3162, -3090, -3017, -2944, -2871, -2799, -2726, -2653, -2581, -2508, -2435, -2363, -2290, -2217, -2145, -2072, -1999, -1927, -1854, -1781, -1708, -1636, -1563, -1490, -1418, -1345, -1272, -1200, -1127, -1054, -982, -909, -764, -691, -618, -473, -182, -109, -37, 181, 254, 399, 472, 617, 690, 763, 836, 908, 981, 1054, 1126, 1199, 1272, 1344, 1417, 1490, 1562, 1635, 1708, 1780, 1853, 1926, 1998, 2071, 2144, 2217, 2289, 2362, 2435, 2507, 2580, 2653, 2725, 2798, 2871, 2943, 3016, 3089, 3161, 3234, 3307, 3380, 3452, 3525, 3598] |
| GAC, 25ºC<br>$T_{EX}$ = 100 ms<br>600 MHz | [30], [-3343, -3270, -3197, -3125, -3052, -2979, -2907, -2834, -2761, -2689, -2616, -2543, -2470, -2398, -2325, -2252, -2180, -2107, -2034, -1962, -1889, -1816, -1744, -1671, -1598, -1526, -1453, -1380, -1308, -1235, -1162, -1089, -1017, -944, -871, -799, -726, -653, -581, -508, -435, -363, -290, -217, -145, -72, 1, 74, 146, 219, 292, 364, 437, 510, 582, 655, 728, 800, 873, 946, 1018, 1091, 1164, 1236, 1309, 1382, 1455, 1527, 1600, 1673, 1745, 1818, 1891, 1963, 2036, 2109, 2181, 2254, 2327, 2399, 2472, 2545, 2618, 2690, 2763, 2836, 2908, 2981, 3054, 3126, 3199, 3272, 3344, 3417, 3490, 3562] |

| | |
|---|---|
| | [90], [-3343, -3270, -3197, -3125, -3052, -2979, -2907, -2834, -2761, -2689, -2616, -2543, -2470, -2398, -2325, -2252, -2180, -2107, -2034, -1962, -1889, -1816, -1744, -1671, -1598, -1526, -1453, -1380, -1307, -1235, -1162, -1089, -1017, -944, -871, -799, -726, -653, -581, -508, -435, -363, -290, -217, -145, -72, 1, 74, 146, 219, 292, 364, 437, 510, 582, 655, 728, 800, 873, 946, 1018, 1091, 1164, 1237, 1309, 1382, 1455, 1527, 1600, 1673, 1745, 1818, 1891, 1963, 2036, 2109, 2181, 2254, 2327, 2400, 2472, 2545, 2618, 2690, 2763, 2836, 2908, 2981, 3054, 3126, 3199, 3272, 3344, 3417, 3490, 3562]<br>[270], [-3344, -3271, -3198, -3126, -3053, -2980, -2908, -2835, -2762, -2689, -2617, -2544, -2471, -2399, -2326, -2253, -2181, -2108, -2035, -1963, -1890, -1817, -1745, -1672, -1599, -1526, -1454, -1381, -1308, -1236, -1163, -1090, -1018, -945, -872, -800, -727, -654, -582, -509, -436, -364, -291, -218, -145, 145, 218, 291, 363, 436, 509, 581, 654, 727, 799, 872, 945, 1018, 1090, 1163, 1236, 1308, 1381, 1454, 1526, 1599, 1672, 1744, 1817, 1890, 1962, 2035, 2108, 2180, 2253, 2326, 2399, 2471, 2544, 2617, 2689, 2762, 2835, 2907, 2980, 3053, 3125, 3198, 3271, 3343, 3416, 3489, 3562]<br>[810], [-3343, -3270, -3197, -3125, -3052, -2979, -2907, -2834, -2761, -2689, -2616, -2543, -2471, -2398, -2325, -2253, -2180, -2107, -2034, -1962, -1889, -1816, -1744, -1671, -1598, -1526, -1453, -1380, -1308, -1235, -1162, -1090, -1017, -944, -871, -799, -726, -653, -581, -508, -435, -363, -290, -217, -145, 1, 73, 146, 219, 291, 364, 437, 510, 582, 655, 728, 800, 873, 946, 1018, 1091, 1164, 1236, 1309, 1382, 1454, 1527, 1600, 1673, 1745, 1818, 1891, 1963, 2036, 2109, 2181, 2254, 2327, 2399, 2472, 2545, 2617, 2690, 2763, 2835, 2908, 2981, 3054, 3126, 3199, 3272, 3344, 3417, 3490, 3562]<br>[2430], [-3343, -3270, -3197, -3125, -3052, -2979, -2907, -2834, -2761, -2689, -2616, -2543, -2470, -2398, -2325, -2252, -2180, -2107, -2034, -1962, -1889, -1816, -1744, -1671, -1598, -1526, -1453, -1380, -1307, -1235, -1162, -1089, -1017, -944, -871, -799, -726, -653, -581, -508, -363, -217, 1, 74, 146, 219, 510, 582, 655, 728, 800, 873, 946, 1018, 1091, 1164, 1237, 1309, 1382, 1455, 1527, 1600, 1673, 1745, 1818, 1891, 1963, 2036, 2109, 2181, 2254, 2327, 2400, 2472, 2545, 2618, 2690, 2763, 2836, 2908, 2981, 3054, 3126, 3199, 3272, 3344, 3417, 3490, 3562] |
| CAC, 25ºC<br>$T_{EX}$ = 100 ms<br>600 MHz | [30], [-3414, -3310, -3206, -3101, -2997, -2893, -2789, -2684, -2580, -2476, -2371, -2267, -2163, -2059, -1954, -1850, -1746, -1641, -1537, -1433, -1329, -1224, -1120, -1016, -911, -807, -703, -599, -494, -390, -286, -181, -77, 27, 131, 236, 340, 444, 549, 653, 757, 861, 966, 1070, 1174, 1279, 1383, 1487, 1592, 1696, 1800, 1904, 2009, 2113, 2217, 2322, 2426, 2530, 2634, 2739, 2843, 2947, 3052, 3156, 3260, 3364, 3469, 3573]<br>[90], [-3414, -3310, -3206, -3101, -2997, -2893, -2789, -2684, -2580, -2476, -2371, -2267, -2163, -2059, -1954, -1850, -1746, -1641, -1537, -1433, -1329, -1224, -1120, -1016, -911, -807, -703, -598, -494, -390, -286, -181, -77, 132, 236, 340, 444, 549, 653, 757, 862, 966, 1070, 1174, 1279, 1383, 1487, 1592, 1696, 1800, 1904, 2009, 2113, 2217, 2322, 2426, 2530, 2634, 2739, 2843, 2947, 3052, 3156, 3260, 3364, 3469, 3573] |

| | |
|---|---|
| | [270], [-3414, -3310, -3206, -3101, -2997, -2893, -2789, -2684, -2580, -2476, -2371, -2267, -2163, -2059, -1954, -1850, -1746, -1641, -1537, -1433, -1329, -1224, -1120, -1016, -911, -807, -703, -599, -494, -390, -286, -181, 27, 131, 236, 340, 444, 549, 653, 757, 862, 966, 1070, 1174, 1279, 1383, 1487, 1592, 1696, 1800, 1904, 2009, 2113, 2217, 2322, 2426, 2530, 2634, 2739, 2843, 2947, 3052, 3156, 3260, 3364, 3469, 3573]<br>[810], [-3414, -3310, -3206, -3101, -2997, -2893, -2789, -2684, -2580, -2476, -2371, -2267, -2163, -2059, -1954, -1850, -1746, -1641, -1537, -1433, -1328, -1224, -1120, -1016, -911, -807, -703, -598, -494, -390, -286, -181, -77, 27, 132, 236, 340, 444, 549, 653, 757, 862, 966, 1070, 1174, 1279, 1383, 1487, 1592, 1696, 1800, 1904, 2009, 2113, 2217, 2322, 2426, 2530, 2634, 2739, 2843, 2947, 3052, 3156, 3260, 3364, 3469, 3573]<br>[2430], [-3414, -3310, -3206, -3101, -2997, -2893, -2789, -2684, -2580, -2476, -2371, -2267, -2163, -2059, -1954, -1850, -1746, -1641, -1537, -1433, -1329, -1224, -1120, -1016, -911, -807, -703, -599, -494, -77, 27, 132, 236, 340, 444, 549, 653, 757, 862, 966, 1070, 1174, 1279, 1383, 1487, 1592, 1696, 1800, 1904, 2009, 2113, 2217, 2322, 2426, 2530, 2634, 2739, 2843, 2947, 3052, 3156, 3260, 3364, 3469, 3573]<br>[5000], [-3414, -3310, -3206, -3101, -2997, -2893, -2789, -2684, -2580, -2476, -2371, -2267, -2163, -2059, -1954, -1850, -1746, -1641, -1537, -1433, -1329, -1224, -1120, -1016, -911, -703, -599, -286, -181, 131, 340, 444, 549, 653, 757, 862, 966, 1070, 1174, 1279, 1383, 1487, 1592, 1696, 1800, 1904, 2009, 2113, 2217, 2322, 2426, 2530, 2634, 2739, 2843, 2947, 3052, 3156, 3260, 3364, 3469, 3573] |

**Table S7.** Spin-lock powers and offsets used in $^{13}$C $R_{1\rho}$ measurements performed for the indicated nuclei at 25 mM NaCl, pH 6.8 and at the indicated temperatures.

| **Sample** | **[RF field power] [offset frequencies]** |
|---|---|
| | [$\omega_1/2\pi$ (Hz)] [$\Omega_{eff}/2\pi$ (Hz)] |
| TAA, 25ºC<br>A-C8<br>600 MHz | [200], [-704.0, -640.0, -576.0, -512.0, -448.0, -384.0, -320.0, -256.0, -192.0, -128.0, -64.0, -10.0, 10.0, 64.0, 128.0, 192.0, 256.0, 320.0, 384.0, 448.0, 512.0, 576.0, 640.0, 704.0]<br>[500], [-1749.0, -1590.0, -1431.0, -1272.0, -1113.0, -954.0, -795.0, -636.0, -477.0, -318.0, -159.0, -10.0, 10.0, 159.0, 318.0, 477.0, 636.0, 795.0, 954.0, 1113.0, 1272.0, 1431.0, 1590.0, 1749.0]<br>[1000], [-3498.0, -3180.0, -2862.0, -2544.0, -2226.0, -1908.0, -1590.0, -1272.0, -954.0, -636.0, -318.0, -10.0, 10.0, 318.0, 636.0, 954.0, 1272.0, 1590.0, 1908.0, 2226.0, 2544.0, 2862.0, 3180.0, 3498.0]<br>[1500], [-5247.0, -4770.0, -4293.0, -3816.0, -3339.0, -2862.0, -2385.0, -1908.0, -1431.0, -954.0, -477.0, -10.0, 10.0, 477.0, 954.0, 1431.0, 1908.0, 2385.0, 2862.0, 3339.0, 3816.0, 4293.0, 4770.0, 5247.0]<br>[2000], [-6996.0, -6360.0, -5724.0, -5088.0, -4452.0, -3816.0, -3180.0, -2544.0, -1908.0, -1272.0, -636.0, -10.0, 10.0, 1272.0, 1908.0, 2544.0, 3180.0, 3816.0, 4452.0, 5088.0, 5724.0, 6360.0, 6996.0] |
| TAG, 25ºC<br>A-C8<br>700 MHz | [500], [-1749.0, -1590.0, -1431.0, -1272.0, -1113.0, -954.0, -795.0, -636.0, -477.0, -318.0, -159.0, -10.0, 10.0, 159.0, 318.0, 477.0, 636.0, 795.0, 954.0, 1113.0, 1272.0, 1431.0, 1590.0, 1749.0]<br>[1000], [-3498.0, -3180.0, -2862.0, -2544.0, -2226.0, -1908.0, -1590.0, -1272.0, -954.0, -636.0, -318.0, -10.0, 10.0, 318.0, 636.0, 954.0, 1272.0, 1590.0, 1908.0, 2226.0, 2544.0, 2862.0, 3180.0, 3498.0]<br>[1500], [-5247.0, -4770.0, -4293.0, -3816.0, -3339.0, -2862.0, -1908.0, -1431.0, -954.0, -477.0, -10.0, 10.0, 477.0, 954.0, 1431.0, 1908.0, 2385.0, 2862.0, 3339.0, 3816.0, 4293.0, 4770.0, 5247.0]<br>[2000], [-6996.0, -6360.0, -5724.0, -5088.0, -4452.0, -3816.0, -3180.0, -2544.0, -1908.0, -1272.0, -636.0, -10.0, 10.0, 636.0, 1272.0, 1908.0, 2544.0, 3180.0, 3816.0, 4452.0, 5088.0, 5724.0, 6360.0, 6996.0]<br>[2500], [-7950.0, -7155.0, -6360.0, -5565.0, -4770.0, -3975.0, -3180.0, -2385.0, -1590.0, -795.0, -10.0, 10.0, 1590.0, 2385.0, 3180.0, 3975.0, 4770.0, 5565.0, 6360.0, 7155.0, 7950.0] |
| TAG, 25ºC<br>A-C1'<br>700 MHz | [200], [-704.0, -640.0, -576.0, -512.0, -448.0, -384.0, -320.0, -256.0, -192.0, -128.0, -64.0, -10.0, 10.0, 64.0, 128.0, 192.0, 256.0, 320.0, 384.0, 448.0, 512.0, 576.0, 640.0, 704.0]<br>[500], [-1749.0, -1590.0, -1431.0, -1272.0, -1113.0, -954.0, -795.0, -636.0, -477.0, -318.0, -159.0, -10.0, 10.0, 159.0, 318.0, 477.0, 636.0, 795.0, 954.0, 1113.0, 1272.0, 1431.0, 1590.0, 1749.0] |

| | |
|---|---|
| | [750], [-2629.0, -2390.0, -2151.0, -1912.0, -1673.0, -1434.0, -1195.0, -956.0, -717.0, -478.0, -239.0, -10.0, 10.0, 239.0, 478.0, 717.0, 956.0, 1195.0, 1434.0, 1673.0, 1912.0, 2151.0, 2390.0, 2629.0]<br>[1000], [-3498.0, -3180.0, -2862.0, -2544.0, -2226.0, -1908.0, -1590.0, -1272.0, -954.0, -636.0, -318.0, 10.0, 318.0, 636.0, 954.0, 1272.0, 1590.0, 1908.0, 2226.0, 2544.0, 2862.0, 3180.0, 3498.0]<br>[2000], [-6996.0, -6360.0, -5724.0, -5088.0, -4452.0, -3816.0, -3180.0, -2544.0, -1908.0, -1272.0, -636.0, -10.0, 636.0, 1272.0, 1908.0, 2544.0, 3180.0, 3816.0, 4452.0, 5088.0, 5724.0, 6360.0, 6996.0]<br>[2500], [-7950.0, -7155.0, -6360.0, -5565.0, -4770.0, -3975.0, -3180.0, -2385.0, -1590.0, -795.0, 795.0, 1590.0, 2385.0, 3180.0, 3975.0, 4770.0, 5565.0, 6360.0, 7155.0, 7950.0] |
| TAT, 25ºC<br>A-C8<br>700 MHz | [200], [-704.0, -640.0, -576.0, -512.0, -448.0, -384.0, -320.0, -256.0, -192.0, -128.0, -64.0, -10.0, 10.0, 64.0, 128.0, 192.0, 256.0, 320.0, 384.0, 448.0, 512.0, 576.0, 640.0, 704.0]<br>[500], [-1749.0, -1590.0, -1431.0, -1272.0, -1113.0, -954.0, -795.0, -636.0, -477.0, -318.0, -159.0, -10.0, 10.0, 159.0, 318.0, 477.0, 636.0, 795.0, 954.0, 1113.0, 1272.0, 1431.0, 1590.0, 1749.0]<br>[750], [-2629.0, -2390.0, -2151.0, -1912.0, -1673.0, -1434.0, -1195.0, -956.0, -717.0, -478.0, -239.0, -10.0, 10.0, 239.0, 478.0, 717.0, 956.0, 1195.0, 1434.0, 1673.0, 1912.0, 2151.0, 2390.0, 2629.0]<br>[1000], [-3498.0, -3180.0, -2862.0, -2544.0, -2226.0, -1908.0, -1590.0, -1272.0, -954.0, -636.0, -318.0, -10.0, 10.0, 318.0, 636.0, 954.0, 1272.0, 1590.0, 1908.0, 2226.0, 2544.0, 2862.0, 3180.0, 3498.0]<br>[2000], [-6996.0, -6360.0, -5724.0, -5088.0, -4452.0, -3816.0, -3180.0, -2544.0, -1908.0, -1272.0, -636.0, -10.0, 10.0, 636.0, 1272.0, 1908.0, 2544.0, 3180.0, 3816.0, 4452.0, 5088.0, 5724.0, 6360.0, 6996.0]<br>[2500], [-7950.0, -7155.0, -6360.0, -5565.0, -4770.0, -3975.0, -3180.0, -2385.0, -1590.0, -795.0, -10.0, 10.0, 795.0, 1590.0, 2385.0, 3180.0, 3975.0, 4770.0, 5565.0, 6360.0, 7155.0, 7950.0] |
| TAT, 25ºC<br>A-C1'<br>700 MHz | [200], [-704.0, -640.0, -576.0, -512.0, -448.0, -384.0, -320.0, -256.0, -192.0, -128.0, -64.0, -10.0, 10.0, 64.0, 128.0, 192.0, 256.0, 320.0, 384.0, 448.0, 512.0, 576.0, 640.0, 704.0]<br>[500], [-1749.0, -1590.0, -1431.0, -1272.0, -1113.0, -954.0, -795.0, -636.0, -477.0, -318.0, -159.0, -10.0, 10.0, 159.0, 318.0, 477.0, 636.0, 795.0, 954.0, 1113.0, 1272.0, 1431.0, 1590.0, 1749.0]<br>[750], [-2629.0, -2390.0, -2151.0, -1912.0, -1673.0, -1434.0, -1195.0, -956.0, -717.0, -478.0, -239.0, -10.0, 10.0, 239.0, 478.0, 717.0, 956.0, 1195.0, 1434.0, 1673.0, 1912.0, 2151.0, 2390.0, 2629.0]<br>[1000], [-3498.0, -3180.0, -2862.0, -2544.0, -2226.0, -1908.0, -1590.0, -1272.0, -954.0, -636.0, -318.0, -10.0, 10.0, 318.0, 636.0, 954.0, 1272.0, 1590.0, 1908.0, 2226.0, 2544.0, 2862.0, 3180.0, 3498.0] |

| | |
|---|---|
| | [2000], [-6996.0, -6360.0, -5724.0, -5088.0, -4452.0, -3816.0, -3180.0, -2544.0, -1908.0, -1272.0, -636.0, -10.0, 10.0, 636.0, 1272.0, 1908.0, 2544.0, 3180.0, 3816.0, 4452.0, 5088.0, 5724.0, 6360.0, 6996.0]<br>[2500], [-7950.0, -7155.0, -6360.0, -5565.0, -4770.0, -3975.0, -3180.0, -2385.0, -1590.0, -795.0, -10.0, 10.0, 795.0, 1590.0, 2385.0, 3180.0, 3975.0, 4770.0, 5565.0, 6360.0, 7155.0, 7950.0] |
| GAG, 25ºC<br>A-C8<br>800 MHz | [150], [-528.0, -480.0, -432.0, -384.0, -336.0, -288.0, -240.0, -192.0, -144.0, -96.0, -48.0, -10.0, 10.0, 48.0, 96.0, 144.0, 192.0, 240.0, 288.0, 336.0, 384.0, 432.0, 480.0, 528.0]<br>[300], [-1045.0, -950.0, -855.0, -760.0, -665.0, -570.0, -475.0, -380.0, -285.0, -190.0, -95.0, -10.0, 10.0, 95.0, 190.0, 285.0, 380.0, 475.0, 570.0, 665.0, 760.0, 855.0, 950.0, 1045.0]<br>[500], [-1749.0, -1590.0, -1431.0, -1272.0, -1113.0, -954.0, -795.0, -636.0, -477.0, -318.0, -159.0, -10.0, 10.0, 159.0, 318.0, 477.0, 636.0, 795.0, 954.0, 1113.0, 1272.0, 1431.0, 1590.0, 1749.0]<br>[1000], [-3498.0, -3180.0, -2862.0, -2544.0, -2226.0, -1908.0, -1590.0, -1272.0, -954.0, -636.0, -318.0, -10.0, 10.0, 318.0, 636.0, 954.0, 1272.0, 1590.0, 1908.0, 2226.0, 2544.0, 2862.0, 3180.0, 3498.0]<br>[2000], [-6996.0, -6360.0, -5724.0, -5088.0, -4452.0, -3816.0, -3180.0, -2544.0, -1908.0, -1272.0, -636.0, -10.0, 10.0, 636.0, 1272.0, 1908.0, 2544.0, 3180.0, 3816.0, 4452.0, 5088.0, 5724.0, 6360.0, 6996.0]<br>[3000], [-10505.0, -9550.0, -8595.0, -7640.0, -6685.0, -5730.0, -4775.0, -3820.0, -2865.0, -1910.0, -955.0, -10.0, 10.0, 955.0, 1910.0, 2865.0, 3820.0, 4775.0, 5730.0, 6685.0, 7640.0, 8595.0, 9550.0, 10505.0] |
| CAG, 25ºC<br>A-C8<br>700 MHz | [200], [-704.0, -640.0, -576.0, -512.0, -448.0, -384.0, -320.0, -256.0, -192.0, -128.0, -64.0, -10.0, 10.0, 64.0, 128.0, 192.0, 256.0, 320.0, 384.0, 448.0, 512.0, 576.0, 640.0, 704.0]<br>[500], [-1749.0, -1590.0, -1431.0, -1272.0, -1113.0, -954.0, -795.0, -636.0, -477.0, -318.0, -159.0, -10.0, 10.0, 159.0, 318.0, 477.0, 636.0, 795.0, 954.0, 1113.0, 1272.0, 1431.0, 1590.0, 1749.0]<br>[1000], [-3498.0, -3180.0, -2862.0, -2544.0, -2226.0, -1908.0, -1590.0, -1272.0, -954.0, -636.0, -318.0, -10.0, 10.0, 318.0, 636.0, 954.0, 1272.0, 1590.0, 1908.0, 2226.0, 2544.0, 2862.0, 3180.0, 3498.0]<br>[2000], [-6996.0, -6360.0, -5724.0, -5088.0, -4452.0, -3816.0, -3180.0, -2544.0, -1908.0, -1272.0, -636.0, -10.0, 10.0, 636.0, 1272.0, 1908.0, 2544.0, 3180.0, 3816.0, 4452.0, 5088.0, 5724.0, 6360.0, 6996.0]<br>[3000], [-10505.0, -9550.0, -7640.0, -6685.0, -5730.0, -4775.0, -3820.0, -2865.0, -1910.0, -955.0, -10.0, 10.0, 955.0, 1910.0, 2865.0, 3820.0, 5730.0, 6685.0, 7640.0, 9550.0, 10505.0] |
| CAG, 15ºC<br>A-C8<br>700 MHz | [150], [-528.0, -480.0, -432.0, -384.0, -336.0, -288.0, -240.0, -192.0, -144.0, -96.0, -48.0, -10.0, 10.0, 48.0, 96.0, 144.0, 192.0, 240.0, 288.0, 336.0, 384.0, 432.0, 480.0, 528.0] |

| | |
|---|---|
| | [300], [-1045.0, -950.0, -855.0, -760.0, -665.0, -570.0, -475.0, -380.0, -285.0, -190.0, -95.0, -10.0, 10.0, 95.0, 190.0, 285.0, 380.0, 475.0, 570.0, 665.0, 760.0, 855.0, 950.0, 1045.0]<br>[500], [-1749.0, -1590.0, -1431.0, -1272.0, -1113.0, -954.0, -795.0, -636.0, -477.0, -318.0, -159.0, -10.0, 10.0, 159.0, 318.0, 477.0, 636.0, 795.0, 954.0, 1113.0, 1272.0, 1431.0, 1590.0, 1749.0]<br>[1000], [-3498.0, -3180.0, -2862.0, -2544.0, -2226.0, -1908.0, -1590.0, -1272.0, -954.0, -636.0, -318.0, -10.0, 10.0, 318.0, 636.0, 954.0, 1272.0, 1590.0, 1908.0, 2226.0, 2544.0, 2862.0, 3180.0, 3498.0]<br>[2000], [-6996.0, -6360.0, -5724.0, -5088.0, -4452.0, -3816.0, -3180.0, -2544.0, -1908.0, -1272.0, -636.0, -10.0, 10.0, 636.0, 1272.0, 1908.0, 2544.0, 3180.0, 3816.0, 4452.0, 5088.0, 5724.0, 6360.0, 6996.0]<br>[3000], [-10505.0, -9550.0, -7640.0, -6685.0, -5730.0, -4775.0, -3820.0, -2865.0, -1910.0, -955.0, -10.0, 10.0, 955.0, 1910.0, 2865.0, 3820.0, 4775.0, 5730.0, 6685.0, 7640.0, 9550.0, 10505.0] |
| CAG, 25ºC<br>A-C1'<br>700 MHz | [200], [-704.0, -640.0, -576.0, -512.0, -448.0, -384.0, -320.0, -256.0, -192.0, -128.0, -64.0, -10.0, 10.0, 64.0, 128.0, 192.0, 256.0, 320.0, 384.0, 448.0, 512.0, 576.0, 640.0, 704.0]<br>[500], [-1749.0, -1590.0, -1431.0, -1272.0, -1113.0, -954.0, -795.0, -636.0, -477.0, -318.0, -159.0, -10.0, 10.0, 159.0, 318.0, 477.0, 636.0, 795.0, 954.0, 1113.0, 1272.0, 1431.0, 1590.0, 1749.0]<br>[1000], [-3498.0, -3180.0, -2862.0, -2544.0, -2226.0, -1908.0, -1590.0, -1272.0, -954.0, -636.0, -318.0, -10.0, 10.0, 318.0, 636.0, 954.0, 1272.0, 1590.0, 1908.0, 2226.0, 2544.0, 2862.0, 3180.0, 3498.0]<br>[2000], [-6996.0, -6360.0, -5724.0, -5088.0, -4452.0, -3816.0, -3180.0, -2544.0, -1908.0, -1272.0, -636.0, -10.0, 10.0, 636.0, 1272.0, 1908.0, 2544.0, 3180.0, 3816.0, 4452.0, 5088.0, 5724.0, 6360.0, 6996.0]<br>[3000], [-10505.0, -9550.0, -7640.0, -6685.0, -5730.0, -4775.0, -3820.0, -2865.0, -1910.0, -955.0, -10.0, 10.0, 955.0, 1910.0, 2865.0, 3820.0, 4775.0, 5730.0, 6685.0, 7640.0, 9550.0, 10505.0] |
| CAG, 15ºC<br>A-C1'<br>700 MHz | [150], [-528.0, -480.0, -432.0, -384.0, -336.0, -288.0, -240.0, -192.0, -144.0, -96.0, -48.0, -10.0, 10.0, 48.0, 96.0, 144.0, 192.0, 240.0, 288.0, 336.0, 384.0, 432.0, 480.0, 528.0]<br>[300], [-1045.0, -950.0, -855.0, -760.0, -665.0, -570.0, -475.0, -380.0, -285.0, -190.0, -95.0, -10.0, 10.0, 190.0, 285.0, 380.0, 475.0, 570.0, 665.0, 760.0, 855.0, 950.0, 1045.0]<br>[500], [-1749.0, -1590.0, -1431.0, -1272.0, -1113.0, -954.0, -795.0, -636.0, -477.0, -318.0, -159.0, -10.0, 10.0, 159.0, 318.0, 477.0, 636.0, 795.0, 954.0, 1113.0, 1272.0, 1431.0, 1590.0, 1749.0]<br>[1000], [-3498.0, -3180.0, -2862.0, -2544.0, -2226.0, -1908.0, -1590.0, -1272.0, -954.0, -636.0, -318.0, -10.0, 318.0, 636.0, 954.0, 1272.0, 1590.0, 1908.0, 2226.0, 2544.0, 2862.0, 3180.0, 3498.0] |

| | |
|---|---|
| | [2000], [-6996.0, -6360.0, -5724.0, -5088.0, -4452.0, -3816.0, -3180.0, -2544.0, -1908.0, -1272.0, -636.0, -10.0, 10.0, 636.0, 1908.0, 2544.0, 3180.0, 3816.0, 4452.0, 5088.0, 5724.0, 6360.0, 6996.0]<br>[3000], [-10505.0, -9550.0, -7640.0, -6685.0, -5730.0, -4775.0, -3820.0, -2865.0, -1910.0, -955.0, 10.0, 955.0, 1910.0, 2865.0, 3820.0, 4775.0, 5730.0, 6685.0, 7640.0, 8595.0, 9550.0, 10505.0] |
| GAC, 25ºC<br>A-C8<br>800 MHz | [200], [-704.0, -640.0, -576.0, -512.0, -448.0, -384.0, -320.0, -256.0, -192.0, -128.0, -64.0, -10.0, 10.0, 64.0, 128.0, 192.0, 256.0, 320.0, 384.0, 448.0, 512.0, 576.0, 640.0, 704.0]<br>[500], [-1749.0, -1590.0, -1431.0, -1272.0, -1113.0, -954.0, -795.0, -636.0, -477.0, -318.0, -159.0, -10.0, 10.0, 159.0, 318.0, 477.0, 636.0, 795.0, 954.0, 1113.0, 1272.0, 1431.0, 1590.0, 1749.0]<br>[1000], [-3498.0, -3180.0, -2862.0, -2544.0, -2226.0, -1908.0, -1590.0, -1272.0, -954.0, -636.0, -318.0, -10.0, 10.0, 318.0, 636.0, 954.0, 1272.0, 1590.0, 1908.0, 2226.0, 2544.0, 2862.0, 3180.0, 3498.0]<br>[2000], [-6996.0, -6360.0, -5724.0, -5088.0, -4452.0, -3816.0, -3180.0, -2544.0, -1908.0, -1272.0, -636.0, -10.0, 10.0, 636.0, 1272.0, 1908.0, 2544.0, 3180.0, 3816.0, 4452.0, 5088.0, 5724.0, 6360.0, 6996.0]<br>[3000], [-10505.0, -9550.0, -8595.0, -6685.0, -5730.0, -4775.0, -3820.0, -2865.0, -1910.0, -955.0, -10.0, 10.0, 955.0, 1910.0, 2865.0, 3820.0, 4775.0, 5730.0, 6685.0, 8595.0, 9550.0, 10505.0] |
| GAC, 25ºC<br>A-C1'<br>800 MHz | [150], [-528.0, -480.0, -432.0, -384.0, -336.0, -288.0, -240.0, -192.0, -144.0, -96.0, -48.0, -10.0, 10.0, 48.0, 96.0, 144.0, 192.0, 240.0, 288.0, 336.0, 384.0, 432.0, 480.0, 528.0]<br>[300], [-1045.0, -950.0, -855.0, -760.0, -665.0, -570.0, -475.0, -380.0, -285.0, -190.0, -95.0, -10.0, 10.0, 95.0, 190.0, 285.0, 380.0, 475.0, 570.0, 665.0, 760.0, 855.0, 950.0, 1045.0]<br>[500], [-1749.0, -1590.0, -1431.0, -1272.0, -1113.0, -954.0, -795.0, -636.0, -477.0, -318.0, -159.0, -10.0, 10.0, 159.0, 318.0, 477.0, 636.0, 795.0, 954.0, 1113.0, 1272.0, 1431.0, 1590.0, 1749.0]<br>[1000], [-3498.0, -3180.0, -2862.0, -2544.0, -2226.0, -1908.0, -1590.0, -1272.0, -954.0, -636.0, -318.0, -10.0, 10.0, 318.0, 636.0, 954.0, 1272.0, 1590.0, 1908.0, 2226.0, 2544.0, 2862.0, 3180.0, 3498.0]<br>[2000], [-6996.0, -6360.0, -5724.0, -5088.0, -4452.0, -3816.0, -3180.0, -2544.0, -1908.0, -1272.0, -636.0, -10.0, 10.0, 636.0, 1272.0, 1908.0, 2544.0, 3180.0, 3816.0, 4452.0, 5088.0, 5724.0, 6360.0, 6996.0]<br>[3000], [-10505.0, -9550.0, -8595.0, -7640.0, -6685.0, -5730.0, -4775.0, -3820.0, -2865.0, -1910.0, -955.0, -10.0, 10.0, 955.0, 1910.0, 2865.0, 3820.0, 4775.0, 5730.0, 6685.0, 7640.0, 8595.0, 9550.0, 10505.0] |
| CAC, 25ºC<br>A-C8<br>700 MHz | [200], [-704.0, -640.0, -576.0, -512.0, -448.0, -384.0, -320.0, -256.0, -192.0, -128.0, -64.0, -10.0, 10.0, 64.0, 128.0, 192.0, 256.0, 320.0, 384.0, 448.0, 512.0, 576.0, 640.0, 704.0] |

| | |
| --- | --- |
| | [500], [-1749.0, -1590.0, -1431.0, -1272.0, -1113.0, -954.0, -795.0, -636.0, -477.0, -318.0, -159.0, -10.0, 10.0, 159.0, 318.0, 477.0, 636.0, 795.0, 954.0, 1113.0, 1272.0, 1431.0, 1590.0, 1749.0]<br>[1000], [-3498.0, -3180.0, -2862.0, -2544.0, -2226.0, -1908.0, -1590.0, -1272.0, -954.0, -636.0, -318.0, -10.0, 10.0, 318.0, 636.0, 954.0, 1272.0, 1590.0, 1908.0, 2226.0, 2544.0, 2862.0, 3180.0, 3498.0]<br>[2000], [-6996.0, -6360.0, -5724.0, -5088.0, -4452.0, -3816.0, -3180.0, -2544.0, -1908.0, -1272.0, -636.0, -10.0, 10.0, 636.0, 1272.0, 1908.0, 2544.0, 3180.0, 3816.0, 4452.0, 5088.0, 5724.0, 6360.0, 6996.0]<br>[3000], [-10505.0, -9550.0, -7640.0, -6685.0, -5730.0, -4775.0, -3820.0, -2865.0, -1910.0, -955.0, -10.0, 10.0, 955.0, 1910.0, 2865.0, 3820.0, 4775.0, 5730.0, 6685.0, 7640.0, 9550.0, 10505.0] |
| CAC, 25ºC<br>A-C1'<br>700 MHz | [200], [-704.0, -640.0, -576.0, -512.0, -448.0, -384.0, -320.0, -256.0, -192.0, -128.0, -64.0, -10.0, 10.0, 64.0, 128.0, 192.0, 256.0, 320.0, 384.0, 448.0, 512.0, 576.0, 640.0, 704.0]<br>[500], [-1749.0, -1590.0, -1431.0, -1272.0, -1113.0, -954.0, -795.0, -636.0, -477.0, -318.0, -159.0, -10.0, 10.0, 159.0, 318.0, 477.0, 636.0, 795.0, 954.0, 1113.0, 1272.0, 1431.0, 1590.0, 1749.0]<br>[1000], [-3498.0, -3180.0, -2862.0, -2544.0, -2226.0, -1908.0, -1590.0, -1272.0, -954.0, -636.0, -318.0, -10.0, 10.0, 318.0, 636.0, 954.0, 1272.0, 1590.0, 1908.0, 2226.0, 2544.0, 2862.0, 3180.0, 3498.0]<br>[2000], [-6996.0, -6360.0, -5724.0, -5088.0, -4452.0, -3816.0, -3180.0, -2544.0, -1908.0, -1272.0, -636.0, -10.0, 10.0, 636.0, 1272.0, 1908.0, 2544.0, 3180.0, 3816.0, 4452.0, 5088.0, 5724.0, 6360.0, 6996.0]<br>[3000], [-10505.0, -9550.0, -7640.0, -6685.0, -5730.0, -4775.0, -3820.0, -2865.0, -1910.0, -955.0, -10.0, 10.0, 955.0, 1910.0, 2865.0, 3820.0, 4775.0, 5730.0, 6685.0, 7640.0, 9550.0, 10505.0] |
| sl-TAG,<br>25ºC<br>A-C8<br>800 MHz | [150], [-528.0, -480.0, -432.0, -384.0, -336.0, -288.0, -240.0, -192.0, -144.0, -96.0, -48.0, -10.0, 10.0, 48.0, 96.0, 144.0, 192.0, 240.0, 288.0, 336.0, 384.0, 432.0, 480.0, 528.0]<br>[300], [-1045.0, -950.0, -855.0, -760.0, -665.0, -570.0, -475.0, -380.0, -285.0, -190.0, -95.0, -10.0, 10.0, 95.0, 190.0, 285.0, 380.0, 475.0, 570.0, 665.0, 760.0, 855.0, 950.0, 1045.0]<br>[500], [-1749.0, -1590.0, -1431.0, -1272.0, -1113.0, -954.0, -795.0, -636.0, -477.0, -318.0, -159.0, -10.0, 10.0, 159.0, 318.0, 477.0, 636.0, 795.0, 954.0, 1113.0, 1272.0, 1431.0, 1590.0, 1749.0]<br>[1000], [-3498.0, -3180.0, -2862.0, -2544.0, -2226.0, -1908.0, -1590.0, -1272.0, -954.0, -636.0, -318.0, -10.0, 10.0, 318.0, 636.0, 954.0, 1272.0, 1590.0, 1908.0, 2226.0, 2544.0, 2862.0, 3180.0, 3498.0]<br>[2000], [-6996.0, -6360.0, -5724.0, -5088.0, -4452.0, -3816.0, -3180.0, -2544.0, -1908.0, -1272.0, -636.0, -10.0, 10.0, 636.0, 1272.0, 1908.0, 2544.0, 3180.0, 3816.0, 4452.0, 5088.0, 5724.0, 6360.0, 6996.0] |
| sl-TAG,<br>25ºC | [200], [-702.0, -624.0, -546.0, -468.0, -390.0, -312.0, -234.0, -156.0, -78.0, -10.0, 10.0, 78.0, 156.0, 234.0, 312.0, 390.0, 468.0, 546.0, 624.0, 702.0] |

| | |
|---|---|
| A-C1'<br>800 MHz | [500], [-1746.0, -1552.0, -1358.0, -1164.0, -970.0, -776.0, -582.0, -388.0, -194.0, -10.0, 10.0, 194.0, 388.0, 582.0, 776.0, 970.0, 1164.0, 1358.0, 1552.0, 1746.0]<br>[1000], [-3501.0, -3112.0, -2723.0, -2334.0, -1945.0, -1556.0, -1167.0, -778.0, -389.0, -10.0, 10.0, 389.0, 778.0, 1167.0, 1556.0, 1945.0, 2334.0, 2723.0, 3112.0, 3501.0]<br>[2000], [-7002.0, -6224.0, -5446.0, -4668.0, -3890.0, -3112.0, -2334.0, -1556.0, -778.0, -10.0, 10.0, 778.0, 1556.0, 2334.0, 3112.0, 3890.0, 4668.0, 5446.0, 6224.0, 7002.0] |
